# The RNA helicase UPF1 associates with mRNAs co-transcriptionally and is required for the release of mRNAs from transcription sites

**DOI:** 10.1101/395863

**Authors:** Anand K. Singh, Subhendu Roy Choudhury, Sandip De, Jie Zhang, Stephen Kissane, Vibha Dwivedi, Preethi Ramanathan, Luisa Orsini, Daniel Hebenstreit, Saverio Brogna

## Abstract

UPF1 is an RNA helicase that is required for efficient nonsense-mediated mRNA decay (NMD) in eukaryotes, and the predominant view is that UPF1 mainly operates on the 3’UTRs of mRNAs that are directed for NMD in the cytoplasm. Here we offer evidence, obtained from *Drosophila*, that UPF1 constantly moves between the nucleus and cytoplasm and that it has multiple functions in the nucleus. It is associated, genome-wide, with nascent RNAs at most of the active Pol II transcription sites and at some Pol III-transcribed genes, as demonstrated microscopically on the polytene chromosomes of salivary gland and by ChIP-seq analysis in S2 cells. Intron recognition seems to interfere with association and translocation of UPF1 on nascent pre-mRNA transcripts, and cells depleted of UPF1 show defects in several nuclear processes essential to correct gene expression – most strikingly, the release of mRNAs from transcription sites and mRNA export from the nucleus.

## Introduction

UPF1 (UP-Frameshift-1) is a universally conserved eukaryotic protein that was first identified in a *Saccharomyces cerevisiae* genetic screen for mutations that enhance up-frameshift tRNA suppression (Culbertson et al., 1980; Leeds et al., 1992), and gained other names – including *NAM7* (*S. cerevisiae*) and *SMG2* (*Caenorhabditis elegans*) – from other genetic screens (Altamura et al., 1992; Hodgkin et al., 1989; Pulak and Anderson, 1993). Cells that lack active UPF1, accumulate mRNAs with nonsense, frameshift or other mutant alleles that introduce a premature translation termination codon (PTC) (Leeds *et al.*, 1991; Pulak and Anderson, 1993).

These observations are generally interpreted as evidence that UPF1 and related proteins are primarily required for nonsense-mediated mRNA decay (NMD), a conserved mRNA surveillance mechanism of eukaryotes that detects and destroys mRNAs at which translation terminates prematurely (Fatscher *et al*., 2015; He and Jacobson, 2015; Karousis *et al*., 2016; Kurosaki and Maquat, 2016). NMD is mainly regarded as a quality control mechanism that prevents cells from wastefully making truncated (and potentially toxic) proteins and that regulates the selective expression of specific mRNA isoforms during cell homeostasis and differentiation (Goetz and Wilkinson, 2017; Lykke-Andersen and Jensen, 2015).

Standard NMD models postulate that UPF1 monitors translation termination on ribosomes by interacting with a peptide release factor (eRF1 or eRF3). However, recent reports on mammalian translation systems have suggested, in contrast to earlier reports on other organisms (Czaplinski *et al*., 1998; Ivanov *et al.*, 2008; Kashima *et al.*, 2006; Keeling *et al.*, 2004; Singh *et al.*, 2008; Wang *et al.*, 2001), that UPF1 does not bind to either of these. They suggested, instead, that UPF3B may contact release factors, slow the termination of translation and facilitate post-termination release of ribosomes – and so fulfil the termination monitoring role that has been assigned to UPF1 (Gao and Wilkinson, 2017; Muhlemann and Karousis, 2017; Neu-Yilik *et al.,* 2017).

UPF1 is an ATP-driven helicase that unwinds RNA secondary structures and so can displace RNA-bound proteins (Bhattacharya *et al.*, 2000; Chakrabarti *et al.*, 2011; Czaplinski *et al.*, 1995; Fiorini *et al.*, 2015). Its helicase activity is required for NMD, but how this helps to target particular transcripts for NMD is not clear (Brogna *et al.*, 2016; Brogna and Wen, 2009). UPF1 is predominantly associated with 3’UTRs of cytoplasmic mRNAs and it might be selectively recruited to or activated on NMD targets with abnormally long 3’UTRs (Karousis et al., 2016; Kurosaki and Maquat, 2016). However, UPF1 appears to bind mRNAs fairly indiscriminately, whatever the position of their stop codon or PTC and whether or not they include NMD-inducing features such as an abnormally long 3’UTR or an exon junction downstream of the stop codon (Hogg and Goff, 2010; Hurt *et al.*, 2013; Zund *et al.*, 2013).

UPF1 is most abundant in the cytoplasm and its roles discussed above depend on ribosomal translation and occur on cytoplasmic mRNAs. However, UPF1 traffics in and out of the nucleus, it interacts with chromatin, it co-purifies with the catalytic subunits of DNA polymerase δ, and UPF1 depletion impairs DNA replication and telomere maintenance (Ajamian et al., 2015; Azzalin and Lingner, 2006; Azzalin et al., 2007; Carastro et al., 2002; Chawla et al., 2011; Mendell et al., 2002). Moreover, there is evidence that UPF1 might contribute directly to RNA processing, at least in specific instances, and is required for nuclear export of HIV-1 genomic RNAs in HeLa cells (Ajamian et al., 2015; Brogna et al., 2016; de Turris et al., 2011; Flury et al., 2014; Varsally and Brogna, 2012).

In the present study we show direct evidence that UPF1 is globally involved in the formation and nuclear processing of mRNAs in *Drosophila*. First, we demonstrate that UPF1 is a highly mobile protein that constantly shuttles between the nucleus and cytoplasm, and its distribution in the cell, with more in the cytoplasm than the nucleus, approximately reflects that of mRNA. UPF1 associates with nascent transcripts on chromosomes – mostly with Pol II transcripts, but also with some Pol III-transcribed genes – and more of the transcript-associated UPF1 is bound with exons than with introns, suggesting that 5’ splice sites might act as a roadblock to the 5’-to-3’ transit of UPF1 along the pre-mRNA. Most strikingly, UPF1 is needed for the efficient release of polyadenylated mRNA from most chromosomal transcription sites and for its export from nuclei. These observations show that UPF1 starts scanning pre-mRNA transcripts whilst they are still being assembled in ribonucleoprotein (RNP) complexes on chromosomes and suggest that it fulfils previously unrecognised role(s) in facilitating nuclear processes of gene expression and mRNA export. The broad and dynamic association with mRNAs redefines UPF1 from being primarily an NMD-inducing factor to being a global player in mRNA processing in the nucleus as well as in the cytoplasm, and might also explain be the reason why none of the prevailing models satisfactorily explains how UPF1 could target specific transcript to NMD.

## Results

### Drosophila anti-UPF1 antibodies

To explore the functions of UPF1, we generated three monoclonal anti-peptide antibodies that target regions of *Drosophila* UPF1 outside the RNA helicase domain: one epitope in the N-terminal flanking regions (antibody 1C13 against Pep2), and two near the C-terminus (Ab 7D17 *vs.* Pep11; and Ab 7B12 *vs.* Pep12) (see Figure S1 and Supplementary Table S1). Each antibody detected UPF1 as a single band by Western blotting of *Drosophila* S2 cell extracts, with minimal cross-reactivity with other proteins, and also detected a second, larger band of the expected molecular mass in extracts from S2 cells that over-express UPF1-GFP (Figure 1A-1C; Figure S1). Unless otherwise indicated, antibody 7B12 was used in the experiments described below. As expected, *UPF1* RNAi specifically reduced the amount of UPF1 in S2 cells (Figure 1B) without affecting the levels of several other proteins we tested as controls (Figure 1B).

**Figure 1.**
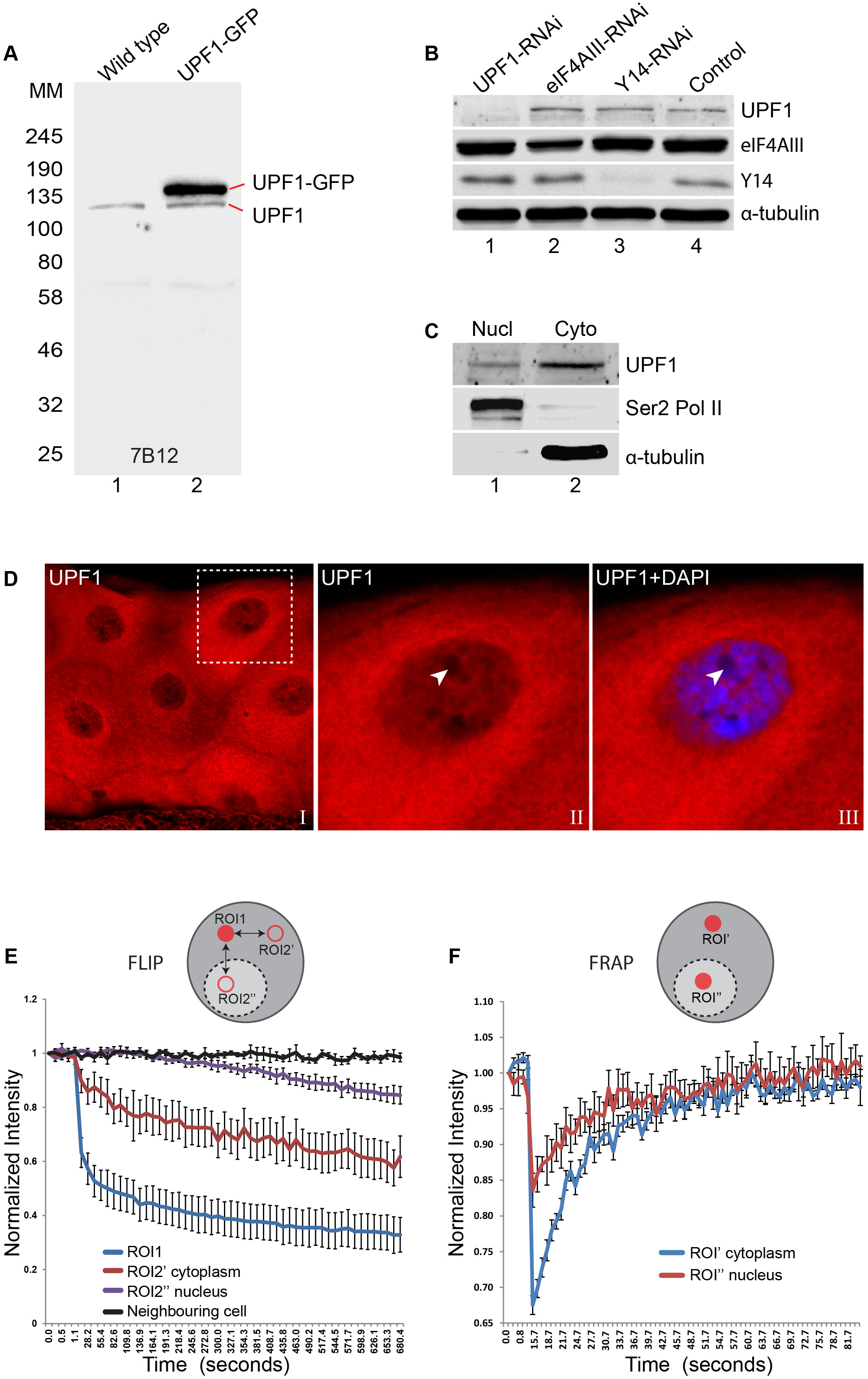
UPF1 continuously shuttles between nucleus and cytoplasm. (**A**) Western blotting of whole-cell lysate from either normal (lane 1) or transfected S2 cells expressing UPF1-GFP (lane 2), probed with the UPF1 monoclonal antibody 7B12. The proteins run according to their expected molecular weights: UPF1 (∼130 kDa), and UPF1-GFP (∼157 kDa) (**B**) Western blotting of S2 cells treated with dsRNA targeting UPF1 or the other RNA binding proteins indicated, used as controls. Separate sections of the membrane were probed with anti-UPF1 (7B12, top row), anti-eIF4AIII (row 2), anti-Y14 (row 3) or anti-α-tubulin (row 4) as a loading control. (**C**) Western blotting of UPF1 following nuclear (Nucl) and cytoplasmic (Cyto) fractionation of S2 cell. RNA Pol II and α-tubulin were detected by the corresponding antibodies using the same blot (shown below). (**D**) Fluorescence immunolocalization of UPF1 (Cy3, red) in 3^rd^ instar larval salivary gland. The arrowheads in panel II and III (magnified view of boxed area in panel I) point to the nucleolus, identified by no DAPI staining, which as other nucleoli shows no UPF1 signal in its centre. (**E**) Plot shows fluorescence loss in photobleaching (FLIP) of UPF1-GFP in salivary gland cells photobleached in ROI1 (red circle, cytoplasm) and then GFP signal measured at the identical time points in two separate ROI2s (red rings), in either cytoplasm or nucleus; both equidistant from ROI1. The different lines show rate of GFP fluorescence loss in either the photobleached ROI1 (blue line), or ROI2’ in the cytoplasm (red line) or ROI2’’ in nucleus (purple line). Change during in fluorescence intensity at equivalent regions in neighbouring cells was measured as a control during the same time-course (black line). Y-axis shows normalized relative fluorescence intensity and X-axis time (seconds) from start of imaging. Quantification based on imaging experiments in 8 different cells. (**F**) Plot shows fluorescence recovery after photobleaching (FRAP) of UPF1-GFP in either cytoplasm (ROI’, blue line) or nucleus (ROI’’, red line) of salivary gland cells. Line values are the average of 8 separate measurements in different cells.

### UPF1 rapidly shuttles between nucleus and cytoplasm

We examined the subcellular localization of immunostained UPF1 in *Drosophila* salivary glands, which are made up of large secretory cells with polytene nuclei. UPF1 was most abundant in the cytoplasm and perinuclear region, and there was also distinct but less intense nuclear staining, mainly around the chromosomes (Figure 1D). Following cell fractionation of S2 cells, α-tubulin and RNA Pol II were, as expected, restricted to the cytoplasmic and nuclear fractions, respectively – and a small proportion of the UPF1 co-purified with nuclei whilst most was in the cytoplasmic fraction (Figure 1C).

Both cytoplasmic and nuclear UPF1 were also present in other larval tissues, with varying relative immunostaining intensities. Perinuclear and intra-nuclear UPF1 were more abundant in Malpighian tubules and gut (Figure S2). In enterocytes (EC), staining was similar in the cytoplasm and within the nucleus, and the most intense UPF1 signal was perinuclear (Figure S2B).

In salivary glands expressing UPF1-GFP (Figure S3A) the perinuclear signals co-localised with binding of wheat germ agglutinin (WGA) – a lectin that predominantly interacts with O-GlcNAc-modified nuclear pore proteins (Mizuguchi-Hata *et al.*, 2013) – and this UPF1-GFP may be associated with components of the nuclear pore complex, as has been proposed for *S. cerevisiae* UPF1 (Nazarenus *et al.*, 2005). It is noteworthy, unexplained though, that *UPF1* RNAi reduced the perinuclear WGA binding in salivary glands (Figure 3SB); cells were also smaller, as would be predicted from its requirement in cell growth during *Drosophila* development (Metzstein and Krasnow, 2006).

**Figure 2.**
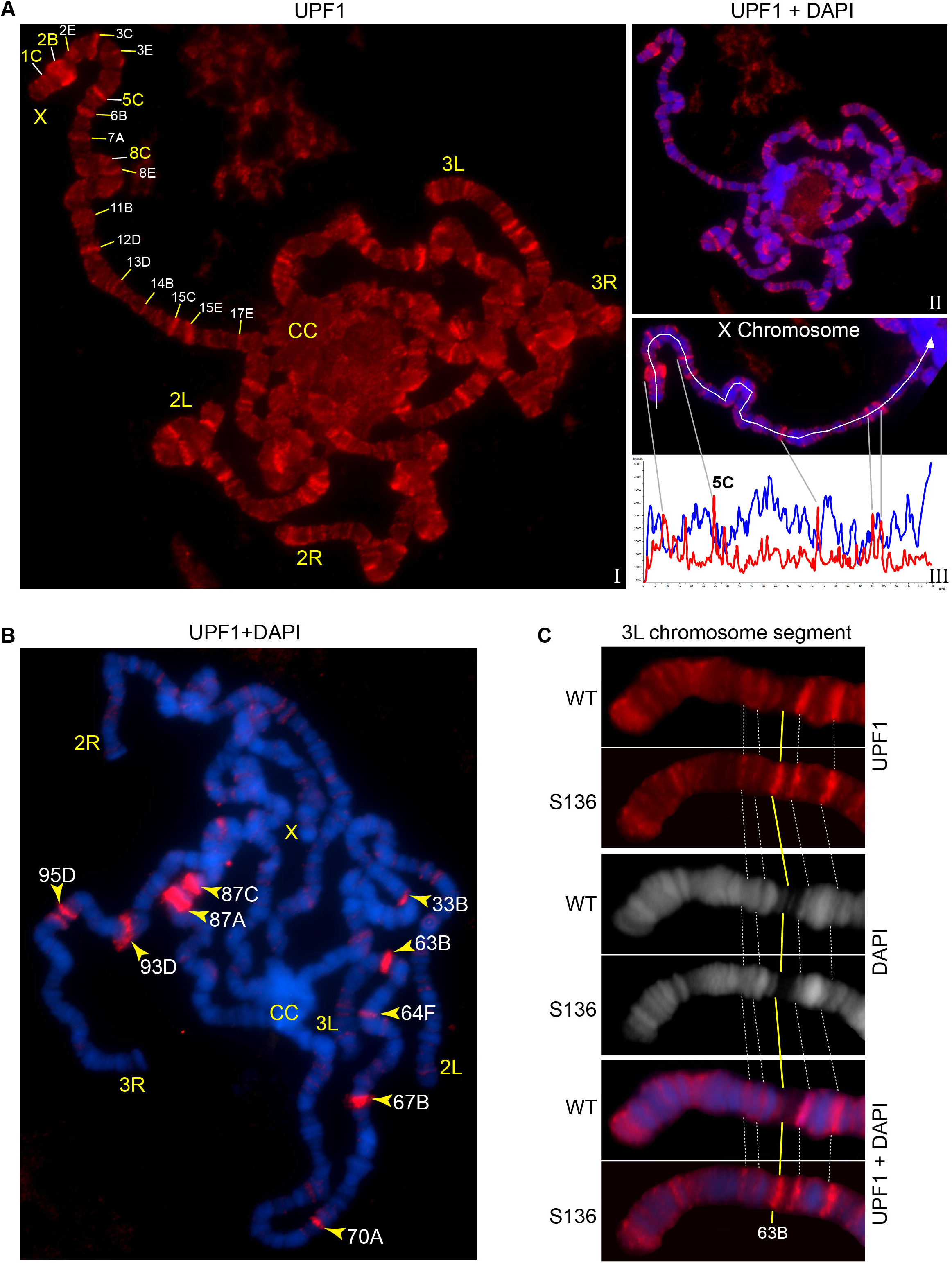
UPF1 binds at transcriptionally active sites on the polytene chromosomes. (**A**) Fluorescence immunolocalization of UPF1 (Cy3, red, I) on polytene chromosomes (DAPI, blue, II). Chromosome arms (X, 2L, 2R, 3L and 3R) and chromocentre (CC) are labelled. The labels indicate cytological locations of interband regions at the X chromosome, presenting apparent UPF1 signal. The line profile (III, white panel) shows signal intensities along the white line drawn on the X chromosome, UPF1 (red) and DAPI (blue). (**B**) Immunolocalization of UPF1 (red) on polytene chromosomes derived from larvae subjected to a 40 min heat shock at 37°C. UPF1 signals are primarily detected at heat shock gene loci, indicated by their cytological locations, using their standard nomenclature. (**C**) Immunolocalization of UPF1 (red) at an ecdysone induced transgene (named S136) located at cytological position 63B (Yellow line) and the same region on the wild type chromosome after ecdysone treatment. The white dotted lines indicate flanking bands as mapping reference. Chromosomes were stained with DAPI (grey in middle panel or blue in bottom panel).

**Figure 3.**
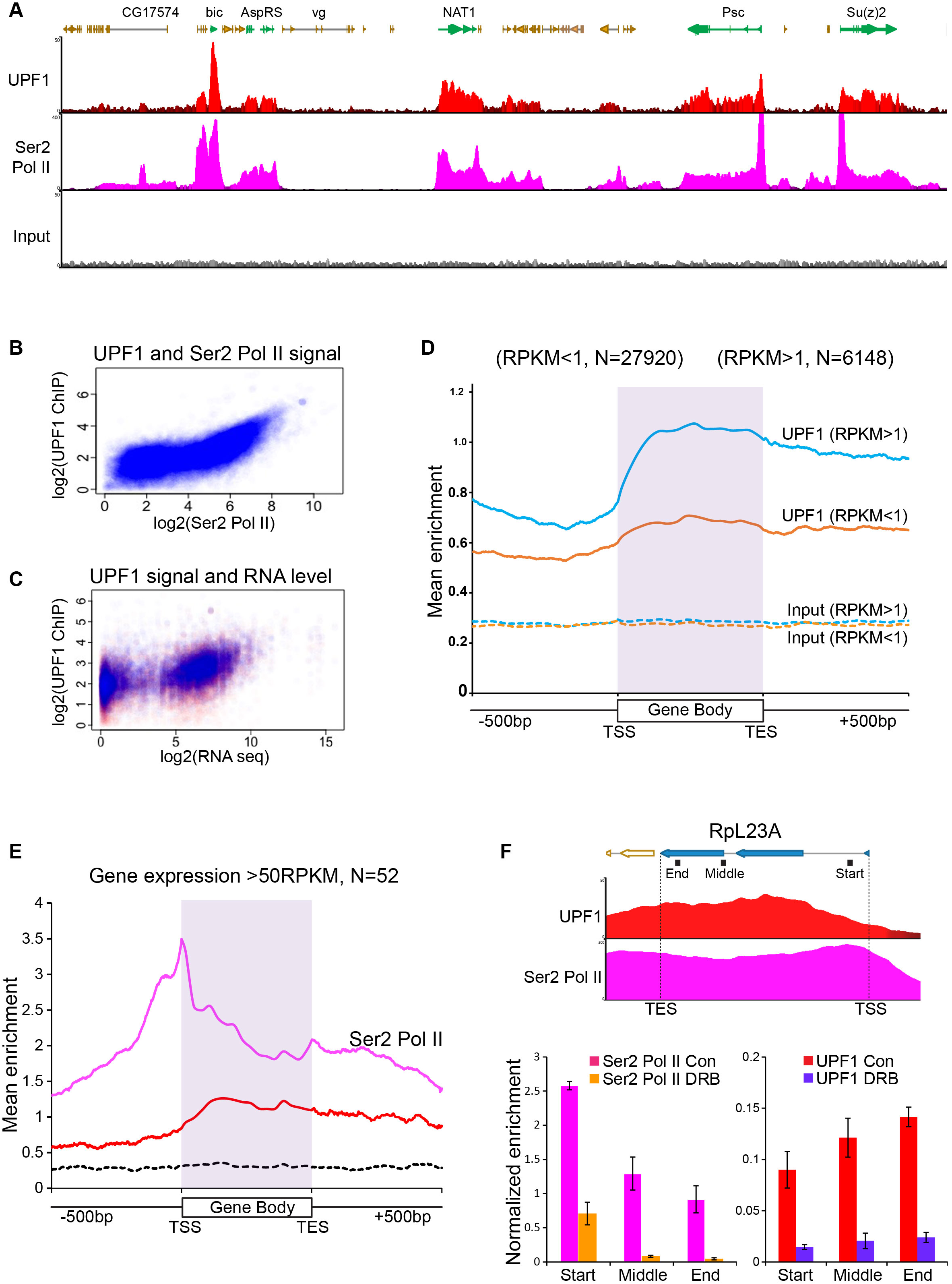
UPF1 associates at Pol II transcription sites. (**A**) Genome browser visualization of UPF1 (red) and Ser2 Pol II (pink) ChIP-seq enrichment profiles at a representative chromosomal region in S2 cells, including highly active genes (green) and low or inactive genes (orange). The input profile (grey) is shown in the bottom panel on the same scale as that of UPF1 (**B**) Scatter plot showing correlation between normalised exon reads in UPF1 and Ser2 Pol II ChIP-seq samples. (**C**) Scatter plot showing relationship between normalised UPF1 ChIP-reads vs. mRNA-seq expression levels; data points corresponding to either exons (blue) or introns (red). (**D**) Metagene profiles showing average UPF1 occupancy at either active (blue, RPKM >1) or inactive/low expressed transcription units (RPKM <1, orange), gene body (scaled to 16 bins of gene full length) plus 500 bp from either end. The number of individual transcription units (N) used for this analysis is given on the top. Corresponding normalised input profiles are shown by dotted lines. (**E**) Superimposed metagene plots of UPF1 (red), Ser2 Pol II (pink) at highly expressed gene loci (RPKM >50). The input enrichment profile for same gene set is shown by the dotted line (black). (**F**) Graph shows ChIP-seq enrichment profiles of UPF1 (red) and Ser2 Pol II (pink) at the RpL23A gene. Bottom, shows real-time PCR quantification of Ser2 Pol II (left) and UPF1 (right) average enrichment at RpL23A gene based on two separate ChIP replicates from either normal or DRB treated S2 cells. The relative position of the three amplicons tested (Start, Middle and End of the gene) are indicated by black boxes underneath the gene schematic on top.

Since UPF1 is present both in cytoplasm and nuclei, with the relative quantities varying between cell-types, we wondered how rapidly UPF1 shuttles between cell compartments. Such trafficking has been reported in HeLa cells, with UPF1 accumulating in the nuclei following treatment with leptomycin B (LMB) (Ajamian et al., 2015; Mendell et al., 2002); this drug selectively inhibits CRM1-mediated protein export from the nucleus in most eukaryotes (Fukuda *et al.*, 1997).

We therefore explored the intracellular localization and dynamics of UPF1 in *Drosophila* salivary glands. Immunostained endogenous UPF1 and UPF1-GFP showed similar intracellular distributions, with an intense cytoplasmic signal and a weaker but still obvious signal in the regions occupied by chromosomes (Figure S4A: the cytoplasmic texture of the salivary cells in these confocal images reflects the fact that they are packed with secretory vesicles at this stage of larval development). In glands treated with LMB for 60 minutes most of the UPF1-GFP was within the nucleus but largely excluded from the nucleolus (Figure S4A, right panels), suggesting that UPF1 exit from the nucleus utilises a CRM1-dependent mechanism. This UPF1 redistribution was rapid in living glands: UPF1 was accumulating in the nucleus by the earliest time we could collect images (within ∼5-6 min), and much of the cell’s UPF1 was in the nucleus within half an hour (Figure S4B).

Heat-shock caused a similar redistribution of much of UPF1 from cytoplasm to nucleus, and this was partially reversed when the tissue was returned to its initial temperature (Figure S4C).

We next used two live cell imaging techniques – Fluorescence Loss in Photo-bleaching (FLIP) and Fluorescence Recovery after Photo-bleaching (FRAP) (Singh and Lakhotia, 2015) – to examine the mobility of UPF1-GFP in salivary gland cells. FLIP revealed that sustained photobleaching of a small area of the cytoplasm led, within the continuously illuminated area, to an initial rapid decrease in UPF1-GFP fluorescence followed by a continued slower reduction. Fluorescence also declined steadily both elsewhere in the cytoplasm, and, more slowly, within the nucleus (Figure 1E). These observations demonstrate ongoing diffusion of UPF1 throughout the cytoplasm, and that nuclear UPF1 can leave the nucleus and enter the photodepletable cytoplasmic UPF1 pool at a fairly steady rate.

The FRAP studies monitored the speed with which unbleached UPF1-GFP diffuses into and repopulates a photobleached region of the cytoplasm or nucleus. Almost all of the UPF1 in each cell compartment was rapidly mobile, and the halftime for repopulation of each bleached area was only a few seconds (Figure 1F).

These observations indicate that UPF1 is freely mobile within cell compartments and that it constantly moves in and out of the nucleus by mechanisms that include the CRM1-dependent nuclear protein export pathway.

### UPF1 associates with transcribing regions of the chromosomes

To gain insight into the role(s) of UPF1 in the nucleus, we used immunostaining to examine whether it associates with the polytene chromosomes of *Drosophila* salivary glands. These well-characterised giant interphase chromosomes are formed after multiple rounds of endoreplication without chromosomal segregation, and they provide a powerful system in which to visualise transcription and pre-mRNA processing at individual gene loci.

UPF1 was present predominantly at interbands and puffs: cytologically distinct chromosome regions in which the chromatin is less condensed and that correspond to transcriptionally active sites (Figure 2A). The immunofluorescence signal appears to be specific, as: a) UPF1-RNAi drastically depletes the endogenous UPF1 chromosomal signal (Figures S5A and S5B); and b) transgenically over-expressed UPF1-GFP, detected either by its fluorescence or with an anti-GFP antibody, shows a similar banding pattern at the chromosomes (Figure S5C).

We then undertook double immunostaining of chromosomes for UPF1 and for Ser2 Pol II – the form of Pol II that transcribes through the main body of genes which is characterised by having the C-terminal domain (CTD) of its largest subunit Ser2-phosphorylated (Boehm *et al.*, 2003). Much of the UPF1 co-localized with Ser2 Pol II, as would be expected from this type of banding pattern (Figure S6A).

The association of UPF1 with the chromosomes depends on transcription. This is illustrated by the changes in UPF1 immunostaining that followed heat-shock, which induces transcription at specific cytological puffs encoding heat-shock proteins and of hsrω lncRNAs at locus 93D (Lakhotia *et al*., 2012). This revealed a pattern of UPF1 association at heat shock puffs and of detachment from most other transcription sites (Figure 2B). UPF1 was recruited to activated heat-shock genes that contained (33B, 63B, 64F, 67B, 70A and 93D) or lacked (87A, 87C and 95D) introns (Figure 2B).

These observations suggested that UPF1 associates with genes that are being transcribed. UPF1 was also recruited to other genes following transcription activation, such as an ecdysone-inducible transgene (S136 at chromosomal position 63B) at normal temperature (Choudhury *et al.*, 2016). No UPF1 was found at this locus while it is inactive, but when ecdysone activated the transgene it produced a cytologically distinguishable transcription puff with which UPF1 was associated (Figure 2C).

### UPF1 mainly associates with Pol II sites that are undergoing transcription and depends on the nascent transcript

We examined the association of UPF1 and of Ser2 Pol II with multiple gene loci by chromatin immunoprecipitation (ChIP) of S2 cell extracts, followed by high-throughput DNA sequencing (ChIP-Seq). UPF1 was associated with many transcriptionally active genes, most of which are Pol II transcription sites. Figure 3A shows enrichment profiles of UPF1 and of Ser2 Pol II across a representative chromosome region. *Actin5C* provided a striking example of correspondence between the ChIP-seq and polytene immunostaining results: it was one of the most UPF1-enriched genes in the ChIP-seq data (Table S2, Figure S9 shows the UPF1 ChIP-seq profile of *Actin5C*) and displayed one of the brightest UPF1 chromosomal signals at the gene locus corresponding to interband 5C on the X chromosome (Figure 2A). The ChIP-seq data also show UPF1 association with a few Pol III genes (Table S2, to be discussed later).

The enrichment profile of UPF1 at Pol II loci closely followed that of Ser2 Pol II, and UPF1 enrichment was greatest at highly expressed genes (Figure 3A; and Figure S8A and S9 show additional examples of UPF1-enriched genes). There was a close correlation between UPF1 and Ser2 Pol II ChIP-seq signals, and also between these and mRNA levels (Figure 3B and 3C). Real-time PCR was used to validate the ChIP-seq data at several genes, both in S2 cells and salivary glands (Figure S7; and other examples are shown below). UPF1-RNAi drastically reduced the UPF1 enrichment at transcription sites, both confirming the specificity of the antibody and validating the ChIP protocol (Fig S7C).

A metagene analysis of the ChIP-seq data shows that UPF1 is associated with genes, and particularly with highly expressed genes (blue trace), throughout their transcription units (Figure 3D) whereas Ser2 Pol II typically shows higher loading around transcription start sites (TSS) – as previously reported in *Drosophila* and other organisms (Adelman and Lis, 2012; Muse et al., 2007). Typically, therefore, most gene-associated UPF1 was further downstream than the TSS-proximal Ser2 Pol II peak, especially at highly expressed genes (Figure 3E: striking examples of this pattern are the *Su(z)2* and *Psc* genes (Figure 3A) and the *α-Tub84B* gene (Figure S8A).

A comparison of the UPF1 loading of genes with different Ser2 Pol II loading profiles suggests that UPF1 association depends on transcription elongation: UPF1 did not associate with genes at which Ser2 Pol II was associated only with the TSS pausing site and which were not being actively transcribed (*e.g. Adam TS-A,* panel 5 in Figure S8A).

The association of UPF1 with Pol II transcription sites is partially sensitive to RNase treatment, suggesting that UPF1 loads onto nascent RNA. This was apparent both for immunostained UPF1 on polytene chromosomes (Figure S6B and C) and when assayed by ChIP at specific genes by qPCR in S2 cells (Figure S7D). UPF1 association was, though, less sensitive to RNase treatment than that of the RNA binding protein hnRNPA1 (Figure S6B-C), which is almost all detached following the same RNase treatment. Some of UPF1 co-purifies with Ser2 Pol II in a standard immunoprecipitation of S2 cell nuclear extracts, the interaction is similarly sensitive to RNase treatment though (Figure S7E): less than that of hnRNPA1, but comparable to that of eIF4AIII, one of the exon junction complex (EJC) proteins that are loaded onto nascent RNAs (Choudhury et al., 2016).

We also examined the effect on salivary glands of 5,6-dichloro-1-beta-D-ribofuranosylbenzimidazole (DRB), a drug that blocks Pol II transcription by inhibiting Ser2 phosphorylation (Bensaude, 2011). In the presence of DRB, unphosphorylated Pol II (Pol II) initiates transcription but does not engage in productive elongation as this would require Ser2-phosphorylated Pol II (Ser2 Pol II) (Adelman and Lis, 2012). DRB treatment left interbands and puffs cytologically unaffected, as expected, but it markedly reduced the amount of UPF1 associated with gene loci (Figure S6D-E), providing further evidence that transcript elongation into the body of the gene is needed for this association to occur. DRB also reduced the association of UPF1 and Ser2 Pol II with genes, such as the highly expressed *RpL23A*, in S2 cells (Figure 3F).

### UPF1 at Pol III transcription sites

UPF1 was found mainly at Pol II transcription sites, most of which are protein-coding genes, but our ChIP-seq data also revealed it at a minority of Pol III genes. The latter included 7SK and both paralogous genes of 7SL snRNAs (Figure S8B) – but not, for example, the much more numerous Pol III-transcribed tRNA genes (Figure S8C, Table S2).

### Intron recognition interferes with UPF1 association with nascent transcripts

UPF1 was recruited both to intron-containing and intronless genes that were undergoing transcription (Figure S9A, and see also the earlier discussion of heat-shock gene activation), so recruitment did not depend on pre-mRNA splicing. Within intron-containing genes, however, more UPF1 was associated with exons than with introns – as can be seen in the ChIP-seq profiles of highly UPF1-enriched genes such as *Xrp1* (Figure 4A; and Figure S9 shows other examples of genes displaying this pattern).

**Figure 4.**
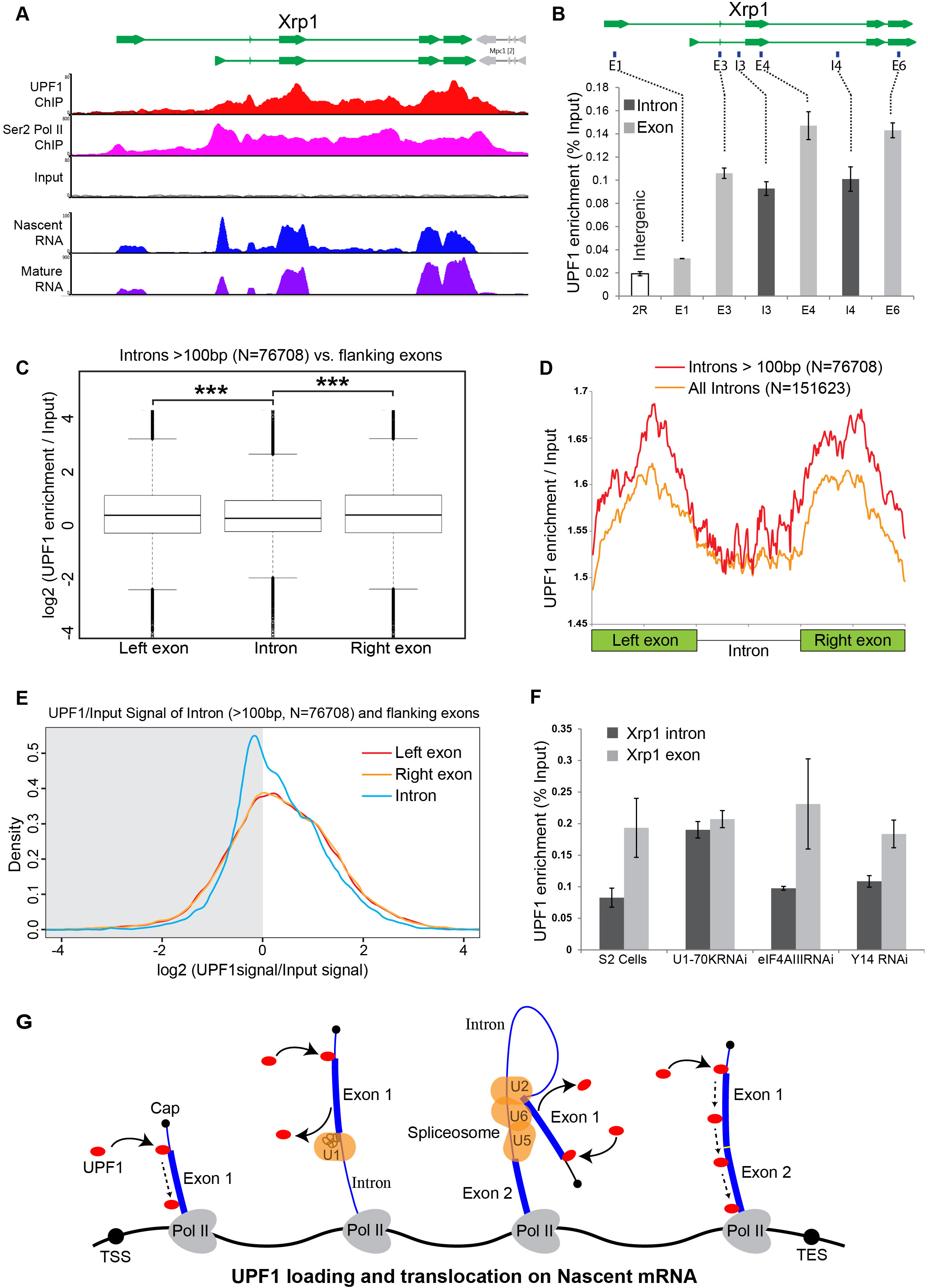
Intron recognition interferes with UPF1 association on nascent transcripts. (**A**) Schematic of the Xrp1 locus (top) showing its two main transcription units. Below, UPF1 (red) and Ser2 Pol II (pink) ChIP-seq profiles at this gene; that of the input is shown below (grey). The bottom two panels show nascent RNA-seq (blue) and poly(A) RNA-seq (purple) profiles. (**B**) Real-time PCR quantification of average enrichment in different regions in either exons (E1, E3, E4 and E6) or introns (I3 or I4) in multiple UPF1 ChIP replicates. (**C**) Box plots of normalised UPF1 ChIP-seq reads mapping at either left exon (shown on left), intron (middle) or right exon (on right). Whiskers correspond to +/- 1.5 interquartile range with respect to quartiles. Wilcoxon rank sum test values are: left exon vs. intron, p-value = 6.737e-08; right exon vs intron, p-value = 2.391e-09; and, left exon vs. right exon p-value = 0.606. *** p < 0.001 for difference in UPF1 signal between intron and its flanking exon. (**D**) Line profile of average UPF1 ChIP-seq/input enrichment expressed as percentage of full length in either exons or intron. Analysis is based on 151623 introns of any length (orange line) or 76708 introns longer than 100 bp (red line) as annotated in the dm6 genome release (**E**) Density plots of UPF1 enrichment at either the exon before (red line), exon after (orange line) or the intron (blue line). The x-axis shows the log2 of the normalized (by input) UPF1 ChIP signal; the right half of the graph shows the density of the values that are enriched, the left half (shadowed) those that are not. (**F**) Real-time PCR quantification of UPF1 ChIP enrichment (two replicates) at intron (I4) or exon (E6) of the Xrp1 gene, in either normal S2 cells or cells RNAi depleted of the proteins indicated. (**G**) Proposed model of how UPF1 scanning of the nascent transcript is connected to intron recognition during spliceosome assembly; spliceosomal snRNPs (U1, U2/U6 and U5) are represented by orange oval shapes, the squiggle drawing within U1 snRNP signifies the base pairing between U1snRNA and the 5’ ss; TSS indicate the transcription start site; and TES, the transcription end site.

This exon-biased UPF1 enrichment was confirmed by real time PCR in multiple ChIP experiments (Figure 4B); and it is genome-wide, as demonstrated by comparing UPF1 association with introns and with their flanking exons in the ChIP-seq data from many genes (Figure 4C, UPF1 enrichment is significantly higher for both the left (*P* = 6.737e-8) and the right flanking exon (2.391e-9); for details of how we corrected for possible bias in chromatin fragmentation or sequencing coverage, see Methods). This pattern is made visually apparent by plotting normalized enrichment in exons and introns, each scaled as a percentage of their full length (Figure 4D), and by comparing the density plots of normalised UPF1 enrichment values in introns and flanking exons, which show more values that are enriched in exons than introns (Fig 4E, compare red and yellow lines *vs.* the blue line in the right half of the graph).

The lower frequency with which UPF1 associated with introns suggested that some features of unspliced transcripts must interfere with the UPF1 interaction. We hypothesised that 5’ splice sites (5’ss) at the starts of introns, where the initial U1 snRNP spliceosome complex would bind, might act as road-blocks to UPF1 translocation along nascent pre-mRNAs and that removal of U1 might allow UPF1 to move on through the intron (Figure 4G). We therefore used ChIP in S2 cells to compare the enrichment of UPF1 at exons and introns in the *Xrp1* gene in cells that had been depleted either of the U1 snRNP protein U1-70K or of Y14 or eIF4III (two of the EJC proteins that bind the nascent pre-mRNA but are not likely to play a direct splicing role in *Drosophila*; see (Choudhury et al., 2016)). The normal bias towards UPF1-exon association in *Xrp1* transcripts was abolished in the U1-70K-depleted cells but persisted in cells depleted of eIF4AIII or Y14 (Figure 4F).

Moreover, genes with the most marked exon-biased UPF1 enrichment, such as *Xrp1*, are efficiently co-transcriptionally spliced (see the Nascent RNA-seq profile in Figure 4A), whereas genes with no detectable exon-biased UPF1 enrichment, such as *CG5059*, are poorly co-transcriptionally spliced (Figure S9C) and are typically expressed at low levels, as reported (Khodor et al., 2011). It seems therefore, that intron recognition interferes with the association of UPF1 with the unspliced nascent transcript.

### Pol II tends to stall near Transcription Start Sites (TSSs) in UPF1-depleted cells

We next determined whether the availability of UPF1 influences the relative amounts of unphosphorylated Pol II and Ser2 Pol II that associate with genes in S2 cells. Both unphosphorylated and, to a lesser extent, Ser2-phosphorylated Pol II were most frequently associated with the start of genes (Figure 5A-5D). Its loading peaked 20 to 60 nucleotides downstream of the TSS (as shown in the expanded depiction in Figure 5B that is based on many more genes), at a position that corresponds to the average distance from TSSs to Pol II pausing sites (Adelman and Lis, 2012). Significantly more unphosphorylated Pol II accumulated there in cells depleted of UPF1 (*P* = 0.011; light blue line in Figure 5B, based on quantifying the aggregate Pol II signal over a +/-100bp span at each of 25440 TSSs): many TSSs showed a 1.2-fold or greater increase in the UPF1-depleted cells (Table S3). Conversely, the amount of Ser2 Pol II associated with these TSSs was unchanged or marginally reduced in UPF1-depleted cells (Figure 5C-5D).

**Figure 5.**
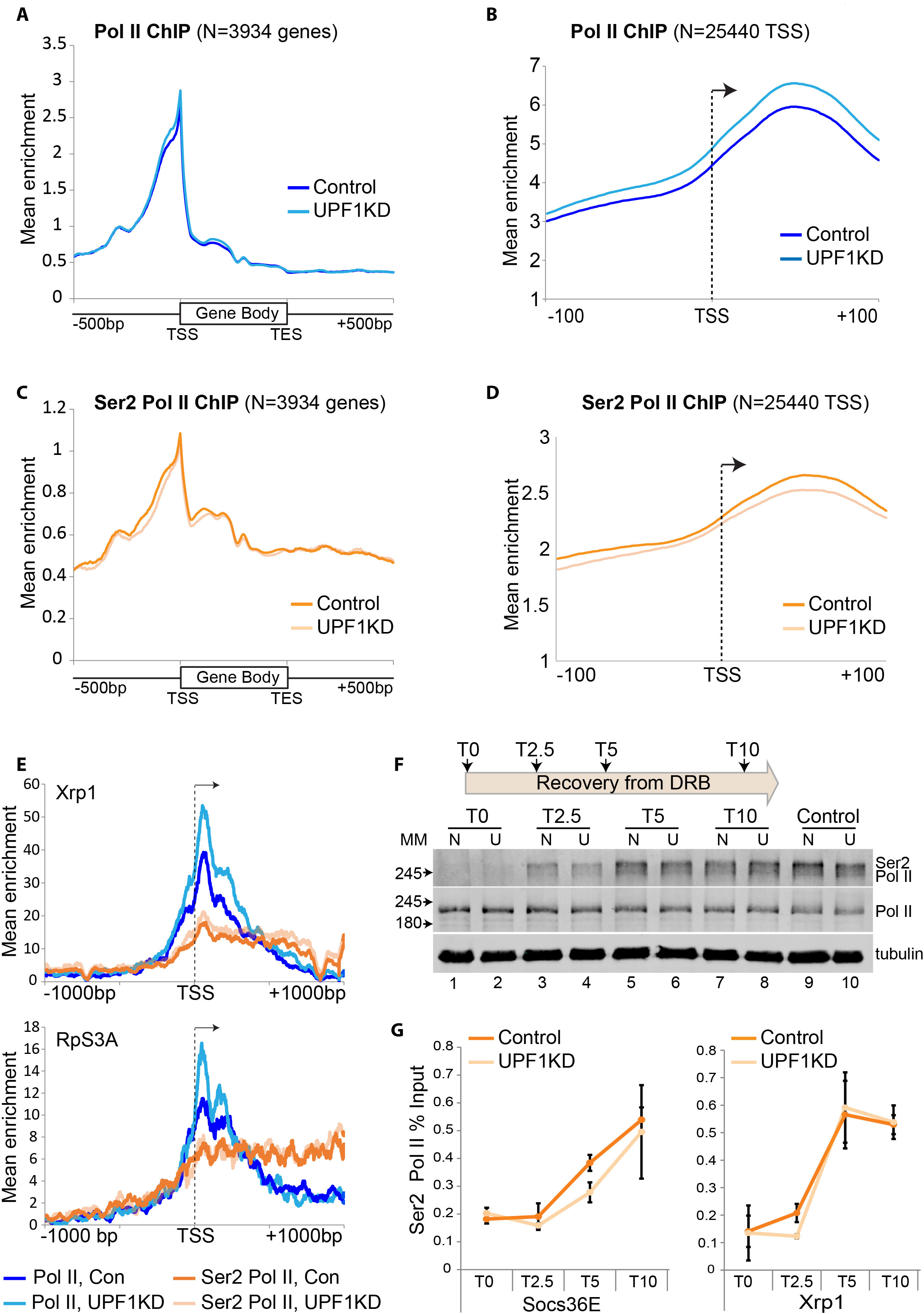
Depletion of UPF1 increases Pol II stalling at TSSs. (**A**) Metagene line plots showing average occupancy of unphosphorylated Pol II at gene bodies (scaled as 16 bins of full length) and ± 500 bp from either end, in normal S2 cells (control, dark blue) or UPF1-RNAi S2 cells (UPF1KD, light blue); analysis based on 3934 (N) transcription units that do not overlap within 500 bp of either ends. (**B**) Metagene plots showing unphosphorylated Pol II at TSS ± 100bp, based on 25440 TSSs that are not closer than 200 bp. (**C**) Metagene plots as in **A** showing Ser2 Pol II occupancy; in normal S2 cells (control, dark orange line) and UPF1-RNAi S2 cells (UPF1KD, light orange line). (**D**) As for B but showing the Ser2 Pol II profiles. (**E**) Normalised ChIP-seq profile of either Pol II (blue lines) or Ser2 Pol II (orange lines) at the Xrp1 gene, ± 1000bp from the TSS (transcript, NM_001275790, the shorter of the two transcripts shown in Figure 3A). The equivalent plots for RpS3A (NM_166714) are shown below. Dark lines refer to normal S2 cells, light lines to UPF1-RNAi, as indicated in the legend below the plots. (**F**) Schematics of the timeframe (top) of the recovery from DRB treatment in S2 cells. Panel below shows Western blot of Ser2 Pol II (top row) and un-phosphorylated Pol II (middle row) at different time points from DRB removal in normal (N) and UPF1-RNAi (U) cells. The α-tubulin was detected as a loading control (bottom row). (**G**) Line graphs show real-time PCR quantifications of average Ser2 Pol II enrichment at either Socs36E gene (on left) or Xrp1 gene (on right) following DRB treatment (T0), at three time points from recovery: 2.5 min (T2.5), 5 minutes (T5) and 10 minutes (T10). Both primer pairs are ∼ 4kb downstream of the TSS (see Table S4).

The increase in unphosphorylated Pol II loading downstream of the TSS in UPF1-depleted cells, alongside fairly constant Ser2 Pol II loading, is illustrated here by *Xrp1* and *RpS3A* (Figure 5E), two genes that are highly transcribed and show strong UPF1 association (Figure 4A, Table S2).

We also assessed transcription by monitoring the repopulation of genes by newly phosphorylated Ser2 Pol II following the withdrawal of DRB treatment. As expected, DRB treatment of S2 cells led to a drastic depletion of Ser2 Pol II and accumulation of unphosphorylated Pol II in cell extracts (Figure 5F, compare Control lanes 9 and 10 with DRB-treated lanes 1 and 2). Ser2 Pol II levels began to recover soon after DRB removal, and were similar to those of untreated control cells within 10 minutes (Figure 5F, compare lanes 7-8 with control lanes 9-10). However, this recovery seemed slower in the UPF1-depleted cells (Figure 5F, compare lanes 3 *vs*. 4 and 5 *vs*. 6). A similarly blunted recovery of gene-associated Ser2 Pol II in UPF1-depleted cells was detected by ChIP at the two gene loci (*Socs36E* and *Xrp1*) that were assayed by real-time PCR (Figure 5G).

Comparable genome-wide observations were made using polytene chromosome spreads. There were no obvious changes in Ser2 Pol II distribution when UPF1 was simply depleted (Figure S10A). When, though, UPF1 was depleted and the glands were also DRB treated, there was then a delay in the recovery of the Ser2 Pol II signal when DRB was removed (Figures S10B, S10C).

Cumulatively, these observations suggest that UPF1 might, by associating with nascent transcripts, influence the phosphorylation of Pol II and hence the transcription of some genes.

### UPF1 depletion leads to nuclear mRNA retention

We also assessed whether depleting UPF1 in the salivary gland cells of 3^rd^ instar larvae would have any effect on mRNA release from transcription sites and its subsequent processing and export from the nucleus.

First we examined the overall cellular distribution of poly(A) RNA – which is referred to from here on simply as poly(A) – by oligo(dT) FISH (fluorescence *in situ* hybridization): this should detect mRNA that has been transcribed, spliced, released from Pol II and polyadenylated. In wild-type cells poly(A) was abundant and fairly evenly distributed throughout the cytoplasm, as would be expected for mature mRNA, and there was little in the nuclei (Figure 6A). By contrast, the nuclei of UPF1-depleted cells retained a substantial amount of poly(A), and the cells appeared to contain less cytoplasmic poly(A) than wild-type cells. Much of the nuclear-retained poly(A) in the UPF1-depleted cells formed large cluster(s) in inter-chromosomal spaces (Figure 6A) that seemed neither to be linked to or in the proximity of any specific chromosomal region(s) or defined transcription site(s).

**Figure 6.**
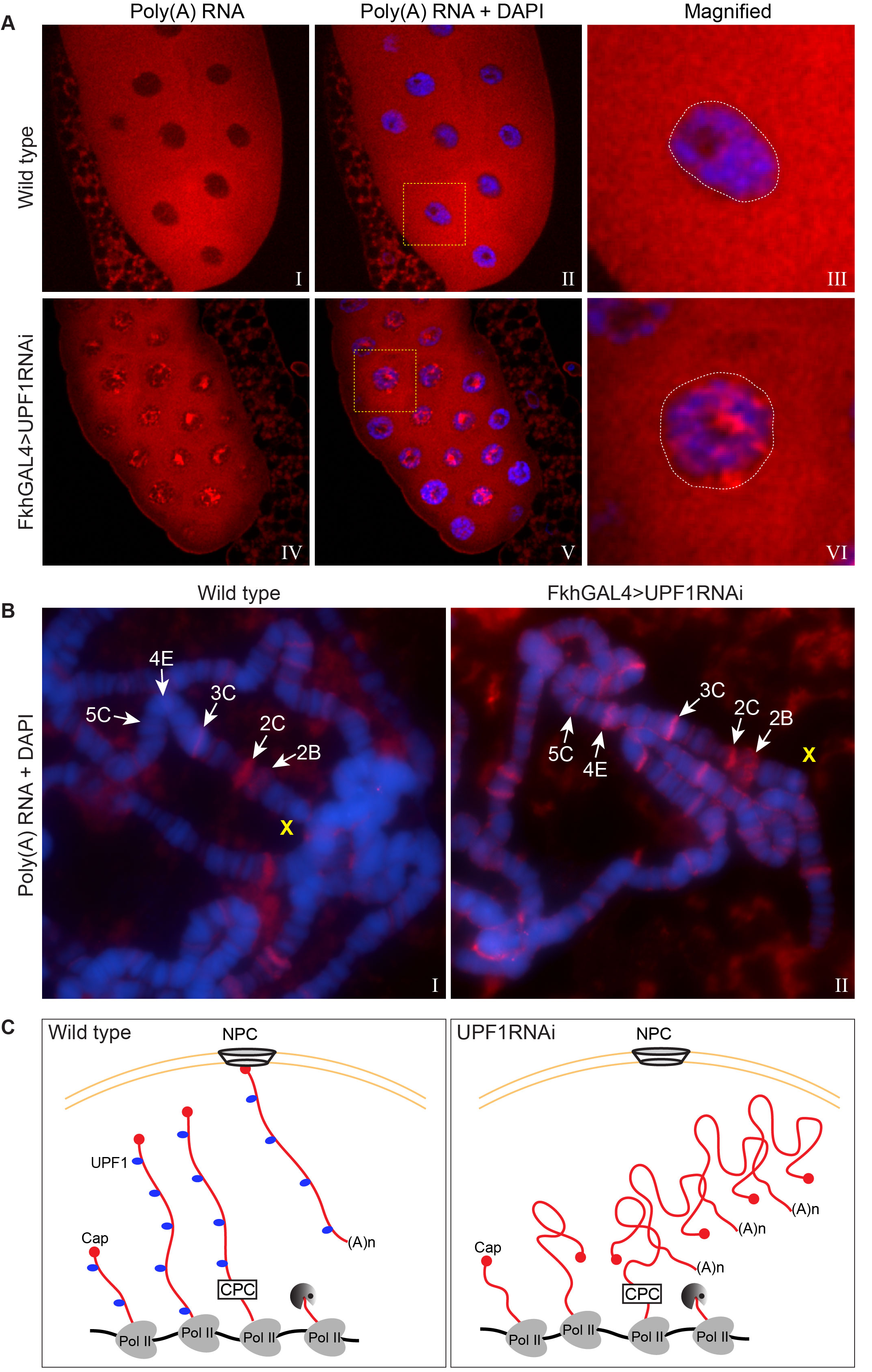
UPF1 knockdown results in nuclear accumulation and transcription sites retention of poly(A) mRNA. (**A)** Fluorescence *in situ* hybridization (FISH) of rhodamine-labelled oligo (dT)45 mer in salivary gland cells of either wild type (top panel) or UPF1-RNAi (bottom panel) 3^rd^ instar larva. Chromosomes were counterstained with DAPI (blue). (**B**) Oligo (dT) 45 mer FISH (as above) of 3^rd^ instar larval salivary gland polytene chromosomes from either wild type or UPF1-RNAi. Chromosomes were counterstained with DAPI (blue). (**C**) Proposed model of accumulation of newly transcribed poly(A) mRNA at the site of transcription in UPF1KD (right) compared with wild type (left). Abbreviations: NPC for nuclear pore complex and CPC for cleavage and polyadenylation complex.

An appreciable amount of poly(A) signal, which was not within clusters, was clearly at the chromosomes though, in the UPF1-depleted cells (Figure 6A, panels III and VI). We therefore used oligo(dT) FISH on polytene chromosome spreads to compare wild-type and UPF1-depleted cells and to assess whether there is retention of poly(A) near transcription sites. There was little poly(A) associated with most of the wild-type chromosomes. However, a few interbands – such as 2C at the distal end of the X chromosome (Figure 6B) – showed clear poly(A) signals (Figure 6B, left panel), suggesting that some completed mRNAs that have been cleaved and polyadenylated remain associated, at least briefly, with transcription sites. Additionally, since UPF1 was obviously not associated with 2C (see Figure 2A), the poly(A) accumulation at 2C in wild-type cells may be a consequence of UPF1 not being normally associated with this transcription site.

Both the number of transcriptional sites showing poly(A) accumulation and the amount of poly(A)RNA associated with these sites were strikingly increased in UPF1-depleted cells (Figure 6B, right panel). For example, there was no visible poly(A) accumulation at site 5C, which corresponds to the highly transcribed *Actin5C* gene, in wild-type, but this band was obviously fluorescent in UPF1-depleted cells. Another example was site 2B – where constitutively expressed *sta* and *rush* are probably the most active genes at this larval stage – which showed a faint poly(A) signal in wild-type glands and a strong signal in UPF1-depleted cells. UPF1 was clearly associated with these transcription sites (2B and 5C) on polytene chromosomes (Fig 2A) and in S2 cells (as detected by ChIP: see Table S2 and Figure S9 for the UPF1 profile of *Actin5C*).

Cumulatively, these data make it clear that UPF1 plays important role(s) both in the release of mRNAs from transcription sites and in their transport out of the nucleus (Figure 6C shows a cartoon of a transcription site of either a wild-type or UPF1 depleted cell with or without mRNA retention).

### Discussion

The RNA helicase UPF1 is usually most abundant in the cytoplasm and is mainly discussed in relation to NMD, leading to the common assumption that it acts mainly on mRNPs that have been exported from the nucleus. In contrast, we present evidence that UPF1 moves constantly within and between cell compartments, that it interacts with mRNA both in the nucleus and the cytoplasm, and that it starts its mRNA association(s) at transcription sites, cotranscriptionally and before pre-mRNA processing is complete.

Within the nucleus we found UPF1 associated with many actively transcribing Pol II sites, to which it seems mainly to be recruited by an interaction with nascent pre-mRNA. More of this transcript-tethered UPF1 is associated with exons than with the introns that separate them. However, this distinction is lost in cells depleted of the spliceosome component U1 snRNP, suggesting that when U1 snRNP is bound to the 5’ss of an intron at the initial stage of splicing, it may hinder UPF1 translocation along the pre-mRNA and cause it to dissociate. It is conceivable that this scanning of pre-mRNAs by UPF1 might influence splice site recognition and pre-mRNA splicing in a manner consistent with reported observations that UPF1 depletion provokes changes in the relative concentrations of many alternatively spliced transcripts in S2 cells (Brooks *et al.*, 2015) – and with the model offered in Figure 4G.

Simple affinity of UPF1 for RNA is not likely to be the primary reason why UPF1 associates with some nascent transcripts, for several reasons: UPF1 does not associate with some highly transcribed Pol II genes, such as spliceosomal snRNAs; nor with snRNA U6 or other highly active Pol III genes; nor with rRNA genes transcribed by Pol I; there would be no differential affinity for introns *vs*. exons within a transcript; and UPF1 appears to be excluded from the nucleolus, including its RNA-packed centre where rRNA genes are transcribed (McLeod *et al*., 2014). What features of some nascent transcripts, most often of Pol II-transcribed genes, dictate that UPF1 becomes associated with them remain to be determined. One obvious candidate would be the 7-methylguanosine (m7G) cap that is added co-transcriptionally to the 5’end of pre-mRNAs but not to Pol I and Pol III transcripts (Ghosh and Lima, 2010). Since the m7G caps added to snRNAs and other small non-mRNA Pol II transcripts are further modified though by hypermethylation to generate 2, 2,7-trimethylguanosine (m(3)G) structures (Mouaikel et al., 2002), it might explain why these classes of transcripts are not UPF1-associated.

The association of UPF1 with nascent transcripts seems to be dynamic, and its putative 5’-to-3’ scanning along RNA seems likely to be fast and, at least on intron-containing pre-mRNA, discontinuous. This pattern also suggests that when it encounters a steric block that cannot be removed UPF1 must be capable of quickly dissociating and re-loading elsewhere on the transcript. *In vitro*, UPF1 can translocate along RNAs over long distances – but only at a maximum scanning velocity of ∼80 base/min (Fiorini *et al.*, 2015), which is much slower than the 2-3 kb/min of Pol II (Fiorini *et al*., 2015; Fukaya *et al*., 2017); possibly UPF1 translocates faster *in vivo*. We did not detect any major impairment of Pol II transcription in UPF1-depleted cells, but Pol II pausing downstream of the TSS was more apparent at some genes – for example, at *Xrp1*, a strikingly UPF1-associated gene. And transcription appears to recover more slowly from DRB inhibition in UPF1-depleted cells.

The most striking effects of UPF1 depletion were retention of poly(A) RNA at transcription sites and then its failure to be exported effectively from the nucleus. Completed mRNA transcripts that have been cleaved and polyadenylated are normally expected to be speedily released from transcription sites, but our data show that this is not always the case. We have both: a) identified some sites on polytene chromosomes that apparently accumulate poly(A) RNA even in wild-type glands; and b) shown that most of the active Pol II genes accumulate poly(A) RNA in UPF1-depleted glands. Poly(A) RNA accumulation in UPF1-depleted cells is most marked at genes with which UPF1 associates strongly in wild-type, such as *Actin5C* (as shown both microscopically and by ChIP-seq). Conversely, those few transcription sites at which poly(A) accumulates even in wild-type cells, may not normally be associated with UPF1, a striking example is transcription site 2C on the polytene chromosomes, which showed the most apparent poly(A) accumulation but no obvious UPF1 association.

Evidence of retention of poly(A) and specific mRNAs in discrete nuclear foci or “dots” has previously been reported in cells defective in RNA processing, initially in mRNA export and processing mutants in yeast (Jensen *et al*., 2001). But whether these foci corresponded to intranuclear mRNP aggregates at sites adjacent to rather than at transcription sites is not clear – and so far these nuclear poly(A) foci have only been reported in cells in which one of several RNA processing reactions are impaired (Abruzzi *et al*., 2006; Paul and Montpetit, 2016). Whether the previously described “dots” correspond to the poly(A) clusters that accumulate in the inter-chromosomal spaces of UPF1-depleted nuclei and/or to accumulations of poly(A) at transcription sites, which we identified here, remains to be determined.

In summary, our results indicate that UPF1 plays an important genome-wide role in nuclear processes of mRNA formation and in their release from transcription sites and export to the cytoplasm, at least in *Drosophila*. Possibly, in the absence of UPF1 function mRNPs acquire or remain in native conformations that hinder their release from the chromosome and make them prone to aggregation and hence nuclear retention. This global role could explain better than NMD why UPF1 is universally conserved in eukaryotes and why its depletion noticeably affects the expression of a large fraction of the genome.

## Materials and Methods

### Antibodies

These antibodies were used for immunostaining: mouse anti-UPF1 (described in this paper 7B12, typically diluted 1:100), mouse IgM anti-Ser2 Pol II (H5, Covance AB_10143905, 1:500), mouse anti-hnRNPA1 (Hrb87F, P11, 1:50)(Hovemann et al., 1991), mouse anti-GFP (B-2, Santa Cruz, SC-9996, 1:200), Tetramethylrhodamine Conjugate Wheat Germ Agglutinin (Thermo Fisher, W7024, 10µg/mL). The antibodies were used in Western blotting: mouse anti-UPF1 (7B12, 1:1000), mouse anti-α-tubulin (Sigma-Aldrich, T5168, 1:2500), rat anti-Ser2 Pol II (3E10, Merck Millipore, 04-1571, 1:5000), mouse anti-Rpb1 (7G5, 1:5000)(Conic et al., 2018); mouse anti-hnRNPA1 (1:200), rabbit anti-eIF4AIII (1:1000), rabbit anti-Y14 (1:1000); the last two antibodies were described previously (Choudhury et al., 2016). The antibodies used in ChIP are mouse anti-UPF1 (7B12, see below, 5-10 µg), rabbit anti-Ser2 Pol II (Abcam, ab5095, 5 µg), mouse anti-Pol II (8WG16, Abcam, ab817, 5µg) and mouse anti-GFP (B-2, Santa Cruz, 5 µg).

### Drosophila Stocks

Flies were reared in standard corn meal fly food media at 24°C. The yw strain was used as wild type. *UAS-UPF1-RNAi* (43144) and *UAS-GFP-UPF1* (24623) were obtained from the Bloomington stock centre. The *forkhead* (*Fkh*) Gal4 has a salivary gland specific expression from early stage of development (Henderson and Andrew, 2000). The transgenes expressing the *lacO*-tagged and ecdysone inducible S136 construct was described before (Choudhury et al., 2016).

### Cell culture and RNA interference

S2 cells were cultured in Insect–XPRESS media (Lonza) supplemented with 10% Fetal Bovine Serum (FBS) and 1% Penicillin-Streptomycin-Glutamine mix (P/S/G, Invitrogen) at 27°C. To make the RNAi constructs for UPF1, eIF4AIII, Y14 and snRNPU1-70k mRNA, the specific sequences were PCR amplified from S2 cell genomic DNA by using corresponding primer pairs (Table S4). Along with the desired gene sequence, all these primer pairs carried the T7 promoter sequence (in bold) at their 5’ end (5’-T**TAATACGACTCACTATAG**GGGAGA-3’). The amplified PCR fragments were purified using Monarch® PCR and DNA Cleanup Kit (T1030S, NEB) and dsRNA was synthesized using the T7 RiboMAX express RNAi system (P1700, Promega). To induce RNAi, a six-well culture dish was seeded with 10^6^ cells/well in serum-free media and mixed with 15 µg of dsRNA/well. Following 1 hr incubation at RT, 2 mL of complete media was added to each well and the cells were incubated for the next three days to knockdown the corresponding RNA and then harvested. The RNAi efficiency of UPF1, eIF4AIII and Y14 was measured by Western blotting while snRNPU1-70k was measured by real time PCR.

### Generation of monoclonal antibodies against Drosophila UPF1

Antigens design, preparation, mice immunization and hybridoma generation were carried out by Abmart (Shanghai). Twelve peptide sequences predicted to be highly immunogenic were selected from *D. melanogaster* UPF1 (Table 1) and cloned in-frame in an expression vector to produce a recombinant protein incorporating all 12 antigens which were used as the immonogen (Abmart, SEAL^TM^ technology). Hybridoma clones were generated and used to induce 18 ascites, which were then screened by Western blotting of S2 cell protein extracts. Out of these, three that showed a single band of the expected size and minimal cross-reactivity were selected and more of the monoclonal antibodies subsequently purified from the corresponding hybridoma cell culture *in vitro*. Unless otherwise specified, 7B12 was used as the anti-UPF1 antibody throughout this study.

### Larval tissue immunostaining

Whole-mount immunostaining was performed as previously described (Choudhury et al., 2016). In brief, the internal organs of 3^rd^ instar larvae were dissected in 1X PBS (13 mM NaCl, 0.7 mM Na2HPO4, 0.3 mM NaH2PO4, pH 7.4) and fixed in 4% formaldehyde for 20 minutes at RT. Tissues were washed in 1XPBS followed by 1% Triton X-100 treatment for 20 minutes. Tissues were washed and incubated in blocking solution (10% Fetal Bovine Serum (FBS), 0.05% Sodium Azide in 1X PBS) for 2 hrs at RT and then incubated in primary antibodies at 4°C overnight. Tissues were washed and further incubated with appropriate fluorescent-tagged secondary antibodies for 2 hrs typically. After washing, tissues were incubated in DAPI (4–6-diamidino-2-phenylindole, Sigma-Aldrich, 1 μg/mL) for 10 minutes and mounted in PromoFluor Antifade Reagent (PK-PF-AFR1, PromoKine) mounting medium and examined using a Leica TCS SP2-AOBS confocal microscope.

### LMB, DRB and larvae heat shock treatment

Wandering 3^rd^ instar larvae were dissected in M3 media and tissues incubated with or without Leptomycin B (LMB, 50 nM) for 1 hr at RT. To examine the real-time effect of LMB treatment in the living cell, salivary glands were dissected in M3 media and incubated with a hanging drop of 50 nM LMB in M3 media in a cavity slide (Singh and Lakhotia, 2015). The fluorescence signal was acquired at 5-minute intervals with a Leica TCS SP2-AOBS confocal microscope. For ecdysone treatment, salivary glands were dissected in M3 media and incubated in 20-hydroxyecdysone (Sigma-Aldrich, H5142, 1µM) for 1 hr at RT. For RNase treatment, salivary glands were dissected in M3 media and incubated in 0.1% Triton X-100 for 2 minutes prior to adding RNase A (Invitrogen, 100µg/mL) a further 1 hr incubation at RT. To examine the effect of 5, 6-Dichlorobenzimidazole 1-β-D-ribofuranoside (DRB) treatment, salivary glands were dissected in M3 media and incubated with DRB (Sigma-Aldrich, 125µM) for 1 hr at RT. For heat shock response, larvae were placed in a pre-warmed microfuge tube lined with moist tissue paper and incubated in water-bath maintained at 37±1°C for 1 hr.

### Live cell imaging (FRAP and FLIP)

Fluorescence recovery after photobleaching (FRAP) and fluorescence loss in photobleaching (FLIP) methods have been previously described (Klonis et al., 2002). Salivary glands expressing UPF1-GFP were dissected from 3^rd^ instar larvae and mounted as a hanging drop in M3 media. For the FRAP the region of interest (ROI, a circle of fixed diameter) was rapidly photobleached with 100 iterations of 100% power Argon laser (488 nm) exposure. Subsequent recovery of fluorescence in the photobleached region was examined at defined time intervals. As a control, fixed cells were examined to confirm irreversible photobleaching. FRAP experiments were carried out on salivary glands at room temperature. The fluorescence signal in ROI was normalized and data analysed following published methods (Phair and Misteli, 2000; Singh and Lakhotia, 2015). FLIP experiments were done as previously described (Phair and Misteli, 2000). Following an acquisition of five control images, GFP fluorescence in ROI1 was continuously photobleached with Argon laser (488 nm) at 100 % power by 50 iterations. The loss in fluorescence in another region of interest, the ROI2 was measured for the same length of time. Fluorescence intensities at ROI1 and ROI2 were normalized and data analysed as described (Nissim-Rafinia and Meshorer, 2011). Both photobleaching experiments have been done using a Leica TCS SP2-AOBS confocal microscope.

### Polytene Chromosomes Immunostaining

Apart from the changes detailed below, the procedure was mostly as previously described (Rugjee et al., 2013). Briefly, actively wandering 3^rd^ instar larvae were dissected in 1X PBS and salivary glands were fixed first with 3.7% formaldehyde in 1X PBS and then with 3.7% formaldehyde in 45% acetic acid for 1 min each (Singh and Lakhotia, 2012). For Pol II immunostaining, salivary glands dissected in 1XPBS were incubated directly with 3.7% formaldehyde in 45% acetic acid for 3 minutes. Salivary glands were squashed in the same solution under the coverslip. Slides were briefly dipped in liquid nitrogen, the coverslips were flipped off with a sharp blade and then immediately immersed in 90% ethanol and stored at 4°C. For immunostaining, the chromosomes were air dried and then rehydrated by incubating the slide with 1XPBS in a plastic Coplin jar. Chromosomes were incubated in blocking solution (as for the tissue immunostaining) for 1 hr at RT and then incubated with primary antibodies diluted in blocking solution in a humid chamber overnight at 4°C. Chromosomes were washed with 1X PBS three times and further incubated with fluorescent tagged appropriate secondary antibodies diluted in blocking solution for 2 hrs at RT in the humid chamber. After washing, chromosomes were counterstained with DAPI and mounted in PromoFluor mounting media. Chromosomes were examined under Nikon Eclipse Ti epifluorescence microscope, equipped with ORCA-R2 camera (Hamamatsu Photonics).

### Fluorescent Oligo (dT) *in situ* hybridization (FISH)

Oligo (dT) FISH was done as previously described for mammalian cells with some adaptations (Tripathi et al., 2015). Salivary glands of 3^rd^ instar larvae were dissected in 1XPBS and fixed in 4% formaldehyde for 15 min at RT. Glands were then washed with 1XPBS and incubated in 0.1% Triton X-100 with 1U/µL Ribolock RNase Inhibitor (ThermoFisher Scientific, EO0381) in 1XPBS for 10 min on ice and then rinsed further with 1XPBS three times with 5 min intervals and then with 2XSSC for 10 min. Salivary glands were incubated with 5ng/µL rhodamine-labelled oligo(dT)45 probe (IDT) in hybridization solution (25% Formamide, 2X SSC pH 7.2, 10% w/v Dextran sulfate (Sigma Aldrich), 1 mg/mL *E. coli* tRNA (Sigma-Aldrich, R1753) for 12 hrs at 42°C. Glands were then washed with freshly made wash buffer (50% Formamide in 2XSSC pH 7.2), followed by 2XSSC, 1XSSC and finally with 1XPBS 3 times each with 5 min interval. Nuclei were counterstained with DAPI and tissues were mounted in PromoFluor Antifade mounting medium and examined under Leica TCS SP2-AOBS confocal microscope.

For polytene chromosomes oligo(dT) FISH, salivary glands were dissected in 1XPBS and incubated with fixing solution (1.85% formaldehyde in 45% acetic acid) for 5 min at RT. Chromosomes were squashed in the same solution and examined immediately under phase-contrast microscope to check if properly spread. Slides with good chromosomes were briefly dipped in liquid nitrogen and the coverslips were flipped off with a sharp blade. Slides were immediately dipped in 90% alcohol and stored at 4°C. Before hybridization, slides were air dried and rehydrated in 1XPBS and then washed and hybridized as described above for whole salivary glands. Chromosomes were counterstained with DAPI and mounted in PromoFluor Antifade mounting medium and examined under Nikon Eclipse Ti epifluorescence microscope.

### Immunoprecipitation

Immunoprecipitation was performed as previously described (Hintermair et al., 2016), with some modifications as detailed below. S2 cells (4 × 10^7^) were harvested and washed with ice-cold 1X PBS containing 1X PhosSTOP (Roche, 04906845001) and 1X cOmplete™, Mini, EDTA-free Protease Inhibitor Cocktail (Roche, 04693159001). Cells were incubated in the hypotonic AT buffer (15 mM HEPES pH 7.6, 10 mM KCl, 5mM MgOAc, 3mM CaCl2, 300 mM Sucrose, 0.1% Triton X-100, 1mM DTT, 1X PhosSTOP, 1X cOmplete™, Mini, EDTA-free Protease Inhibitor Cocktail and 1U/µL Ribolock RNase Inhibitor) for 20 min on ice and lysed with 2mL Dounce homogenizer by 30 strokes with the tight pestle. Lysate was centrifuged at 5000 RPM for 5 min at 4°C in microcentrifuge and the nuclear pellet was resuspended in 500 μL IP buffer (50 mM Tris-HCl pH 8.0, 150 mM NaCl, 1% NP-40 (Roche), 1X PhosSTOP, 1X cOmplete™, Mini, EDTA-free Protease Inhibitor Cocktail, 1U/µL Ribolock RNase Inhibitor) for 20 min on ice. Nuclear lysates were sonicated using a Bioruptor sonicator (Diagenode) for 3 cycles of 30 sec ON and 30 sec OFF with maximum intensity. Following sonication, the lysates were centrifuged at 13000 RPM for 15 min at 4°C in a microfuge and the antibody (5μg) was added to the clear supernatant, with or without addition of RNase A (100µg/mL), and incubated overnight at 4°C on a rocker. Following incubation, 20 μL of prewashed paramagnetic Dynabeads (ThermoFisher Scientific, 10004D) were added and incubated further for 2 hrs at 4°C on a rocker. Beads were washed 5 times with IP buffer using a magnetic rack and proteins were extracted by adding 40μL SDS-PAGE sample buffer.

### ChIP-Seq

S2 cells (2 × 10^7^) were harvested and fixed with 1% formaldehyde (EM grade, Polyscience) for 10 min at RT. Following fixation, cross-linking reaction was stopped by adding 125 mM Glycine for 5 min at RT. Cells were centrifuged at 2000 RPM for 5 min at 4°C, the pellet was washed twice with ice-cold 1X PBS containing 1X cOmplete™, Mini, EDTA-free Protease Inhibitor Cocktail. The cell pellet was resuspended in 1 mL of cell lysis buffer (5mM PIPES pH 8.0, 85mM KCl, 0.5% NP-40) supplemented with 1X cOmplete™, Mini, EDTA-free Protease Inhibitor Cocktail and 1X PhosStop and incubated for 10 min at 4°C. Cells were centrifuged and the pellet was resuspended in 1 mL nuclear lysis buffer (50mM Tris pH 8.0, 10 mM EDTA, 1.0% SDS) supplemented with 1X cOmplete™, Mini, EDTA-free Protease Inhibitor Cocktail and 1X PhosStop and incubated for 10 min at 4°C. The cell suspension was further diluted with 500 µL IP dilution buffer (16.7 mM Tris pH 8.0, 1.2 mM EDTA, 167 mM NaCl, 1.1% Triton X-100, 0.01% SDS) and sonicated for 5 cycles at 30sec ON, 30 sec OFF with maximum intensity by using a Bioruptor sonicator (Diagenode); this produced an average fragment size of ∼500 bp. Samples were centrifuged at 13000 RPM for 20 min in a microcentrifuge and the clear supernatant was transferred to a 15 mL tube. An aliquot of 100 µL supernatant was kept to extract input DNA. The supernatant was further diluted with 5 volume of IP dilution buffer. For each ChIP, typically we added 5 to 10 µg of antibody to this supernatant and incubated overnight at 4°C on a rocker. Prewashed 20 µL Dynabeads were added to the lysate-antibody mix and incubated further for 1 hr at 4°C on a rocker. Beads were washed 6 times with low salt buffer (0.1% SDS, 1% Triton X-100, 2mM EDTA, 20 mM Tris pH 8.0, 150 mM NaCl), once with high salt buffer (0.1% SDS, 1% Triton X 100, 2mM EDTA, 20 mM Tris pH 8.0, 500 mM NaCl) and once with 1X TE buffer (10mM Tris pH 8.0, 1mM EDTA). The beads were then incubated with 250 µL elution buffer (0.1M NaHC03, 1% SDS) at RT for 15 min and eluted chromatin was reverse cross-linked by adding 38 µL de-crosslinking buffer (2M NaCl, 0.1M EDTA, 0.4M Tris pH 7.5) and then incubated at 65°C for overnight on a rotator. Proteins were digested by adding 2 µL Proteinase K (50 mg/mL) and incubated at 50°C for 2 hrs on a rocker. DNA was isolated by using Monarch® PCR https://www.neb.com/products/t1030-monarch-pcr-dna-cleanup-kit-5-ug and DNA Cleanup Kits. Real-time PCR quantification of DNA samples was carried out using the SensiFAST SYBR Hi-ROX Kit (Bioline, BIO-92005) in 96-well plates using an ABI PRISM 7000 system (Applied Biosystems). For NGS sequencing, ChIP and input DNA were further fragmented to 200 bp fragment size using a Bioruptor Pico (Diagenode). All ChIP-DNA libraries were produced using the NEBNext Ultra II DNA Library Prep Kit (New England Biolab E7645L) and NEBnext Multiplex Oligos for Illumina Dual Index Primers (New England Biolabs E7600S), using provided protocols with 10ng of fragmented ChIP DNA. Constructed libraries were assessed for quality using the Tapestation 2200 (Agilent G2964AA) with High Sensitivity D1000 DNA ScreenTape (Agilent 5067-5584). Libraries were tagged with unique barcodes and sequenced simultaneously on a HiSeq4000 sequencer.

### Nascent RNA isolation from S2 cells

Nascent RNA isolation was performed as previously described (Khodor et al., 2011). Briefly, S2 cells (4 × 10^7^) were harvested and washed twice with ice-cold 1X PBS via centrifugation at 2000g for 5 min each. Cells were resuspended in 1 mL ice-cold buffer AT and incubated on ice for 10 minutes. Cells were lysed using a 2mL Dounce homogenizer by 30 strokes with the tight pestle. The lysate was divided into two aliquots and each aliquot of 500 µL was layered over a 1mL cushion of buffer B (15 mM HEPES-KOH at pH 7.6, 10 mM KCl, 5 mM MgOAc, 3 mM CaCl2, 1 M sucrose, 1 mM DTT, 1X cOmplete™, Mini, EDTA-free Protease Inhibitor Cocktail), and centrifuged at 8000 RPM for 15 min at 4°C in a microcentrifuge. The supernatant was removed and the pellet was resuspended in 5 volumes of nuclear lysis buffer (10 mM HEPES-KOH pH 7.6, 100 mM KCl, 0.1 mM EDTA, 10% Glycerol, 0.15 mM Spermine, 0.5 mM Spermidine, 0.1 M NaF, 0.1 M Na3VO4, 0.1 mM ZnCl2, 1 mM DTT, 1X cOmplete™, Mini, EDTA-free Protease Inhibitor Cocktail and 1U/µL Ribolock RNase Inhibitor) and resuspended using a 2mL Dounce homogenizer by 3 strokes with loose pestle and 2 strokes with tight pestle. Equal volume of 2X NUN buffer (50 mM HEPES-KOH pH 7.6, 600 mM NaCl, 2 M Urea, 2% NP-40, 1 mM DTT, 1X cOmplete™, Mini, EDTA-free Protease Inhibitor Cocktail and 1U/µL Ribolock RNase Inhibitor) was added to the nuclear suspension drop by drop while vortexing and the suspension was placed on ice for 20 min prior to spinning at 13,000 RPM for 30 min at 4°C. The supernatant was removed and TRI Reagent (Sigma, T9424) was added to the histone–DNA-Pol II-RNA pellet. The TRI Reagent–pellet suspension was incubated at 65°C with intermittent vortexing to dissolve the pellet, and RNA was extracted following the manufacturer’s protocol. Poly(A) depletion was performed with Dynabeads™ Oligo(dT)25 (ThermoFisher Scientific). The purification of nascent RNA was assessed by RT-PCR of CG12030, CG5059 and CG10802 genes which have slow rates of co-transcriptional splicing (Khodor et al., 2011); cDNA synthesis was performed using qScript cDNA synthesis kit (Quanta Biosciences, 95047-025).

### RNA-seq

Extracted RNA samples were quantified using a Nanodrop-8000 Spectrophotometer (ThermoFisher ND-8000-GL) to assess quality and to determine concentrations. Aliquots of each sample were diluted to ∼5ng/µl, and tested with an Agilent Tapestation 2200 (Agilent G2964AA) using High Sensitivity RNA ScreenTapes kit (Agilent 5067-5579) to determine the RNA Integrity Number.

Total-RNA (1µg) was first poly(A) selected using the NEBNext® Poly(A) mRNA Magnetic Isolation Module (New England Biolabs E7490L) prior to library construction. Nascent RNA samples (100 ng) were processed without poly(A) selection. RNA libraries were prepared using a NEBNext Ultra Directional RNA Library Prep Kit (New England Biolab E7420L) and NEBnext Multiplex Oligos for Illumina Dual Index Primers (New England Biolabs E7600S), following standard protocols. RNA libraries were checked for quality using the Tapestation 2200 (Agilent G2964AA) with High Sensitivity D1000 DNA ScreenTape (Agilent 5067-5584). Multiplexed libraries were sequenced (50-bp single-end reads) on a HiSeq4000 sequencer.

### CHIP-seq and RNA-seq data analysis

ChIP-seq and RNA-seq data were initially viewed and analysed using the Lasergene Genomics Suite version 14 (DNASTAR). Pre-processing, assembly and mapping of the sequencing reads in the FASTQ files were performed by the SeqMan NGen software of this package automatically after selecting the NCBI *D. melanogaster* Dm6 genome release and accompanying annotations. Assembly and alignment output files for each genome contig were then analysed with the ArrayStar and GenVision Pro software (from the same package) to view and compare data track on the genome. Profiles at selected regions were saved as high-resolution images.

To perform the metagene analyses, an index for Dm6 was downloaded from the HISAT2 website. HISAT2 v2.1.0 was then used to align the FASTQ files on it. The resulting SAM files were converted to BAM format, sorted, and indexed with Samtools 1.6. For the cytoplasmic RNA-seq data, the NCBI RefSeq gene annotations for Dm6 were downloaded as a GTF file from UCSC Table Browser (genome.ucsc.edu). Custom scripts were then used to produce read counts per gene in an HTSeq-count compatible format based on the GTF file. Transcript lengths were also obtained from the GTF file and used together with total mapped sequencing reads to convert counts into RPKM values. For both ChIP-seq and nascent RNA-seq data, the BAM files were converted to Bedgraph files. This was carried out with the genomeCoverageBed command and options -bga and -ibam from the Bedtools v2.26.0 suite. Custom Perl scripts were then used to filter the Dm6 annotations either for genes separated by a minimum distance to avoid overlapping signals or RNA-seq expression levels. Subsequently, custom scripts were used to extract the signal from the Bedgraph files for each entry in the filtered gene list. A single base resolution was used for flanking regions, while the signal in gene bodies was binned into 16 bins to take account of different gene lengths. Each dataset was normalized by the total mapped sequencing reads in that dataset. Cross-referencing between different datasets was done based on the ‘name’ field, after filtering the annotations for multiple entries with the identical name.

A custom script was used to extract from the Bedgraph files the sequencing read coverage for each exon/intron/exon region in the dm6 annotation file xon_fly_gene (downloaded from UCSC Table Browser). To normalise for any bias in the sequencing, the UPF1 signal of each exon or intron was divided by the average coverage in the input sample. The fold change of UPF1 signal/input signal in introns was compared to that of their flanking exons using Wilcox.test (two sided and unpaired) in R (www.r-project.org). This analysis was done using either all introns annotated in Dm6 (151623) or those longer than 100bp (76708), in either case flanking exons are significantly more enriched than introns. Here we have shown the result of analysis using just the longer introns as these were considered to be more informative because of the predicted lower resolution of ChIP at discriminating between closely adjacent sequences and because the lower sequencing coverage of shorter introns compared to longer introns. All ChIP-seq and RNA-seq raw sequencing data and Bedgraph files were deposited in the GEO repository (Accession No GSE116808)

## Acknowledgments

We thank Bob Michell for critically reading the manuscript and valuable discussions. Thanks also to Michael Rosbash and Michael Marr (USA) for providing a detailed Nascent RNA-seq protocol, Harald Saumweber (Germany) for the P11 antibody and Laszlo Tora (France) for the 7G5 antibody. Bloomington is also acknowledged for providing fly stocks. Thanks also to Alessandro Di Maio and the Birmingham Advanced Light Microscopy (BALM) facility; our School Drosophila research community, fly food facility and Shrikant Jondhale for fly stocks maintenance; our NGS facility; and, Mike Tomlinson for providing Odyssey infrared imaging system for Western blot detection. We thank also Pawel Grzechnik for reagents, and his and our group for help and continuous discussions. This project was funded by a Leverhulme Trust (RPG-2014-291) and BBSRC (BB/M022757/1) project grants, and at its start, Wellcome Trust (9340/Z/09/Z) to SB. DH was supported by BBSRC grants BB/M017982/1 and BB/L006340/1.

## Supplementary Figures and legends

**Figure S1.**
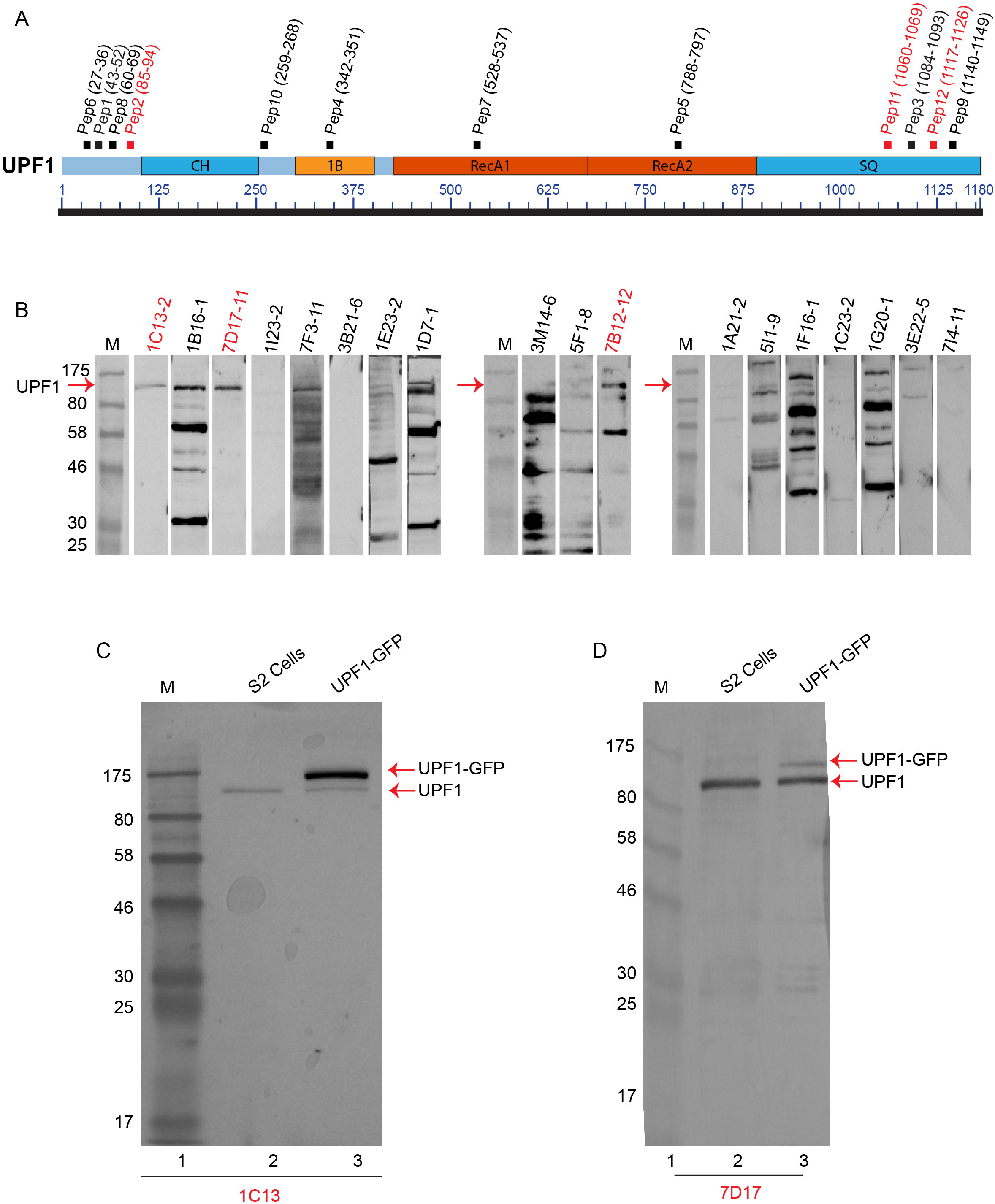
Generation of monoclonal antibodies against Drosophila UPF1. (**A**) Schematics of UPF1 showing its different structural domains. The peptides used as immunogens and their respective amino acid locations are given in brackets (sequences are in Table S1). The peptides indicated by red color produced the monoclonal antibodies with highest specificity, as shown below (**B**) Western blotting of S2 cell protein extracts, probed with 18 ascites induced with hybridomas previously screened for their reactivity to the corresponding peptides (terminal numbers correspond to different peptides indicated in A). Lanes labelled M show a molecular weight marker. (**C**) Western blotting of whole-cell S2 lysate with the UPF1 monoclonal antibody (mab) 1C13. From either normal (lane 2) or transfected S2 cells expressing UPF1-GFP (lane 3) in which the two bands correspond to either endogenous UPF1 or UPF1-GFP. (**D**) As in C, using mab 7D17. Western blotting with 7B12 is shown in Figure 1A.

**Figure S2.**
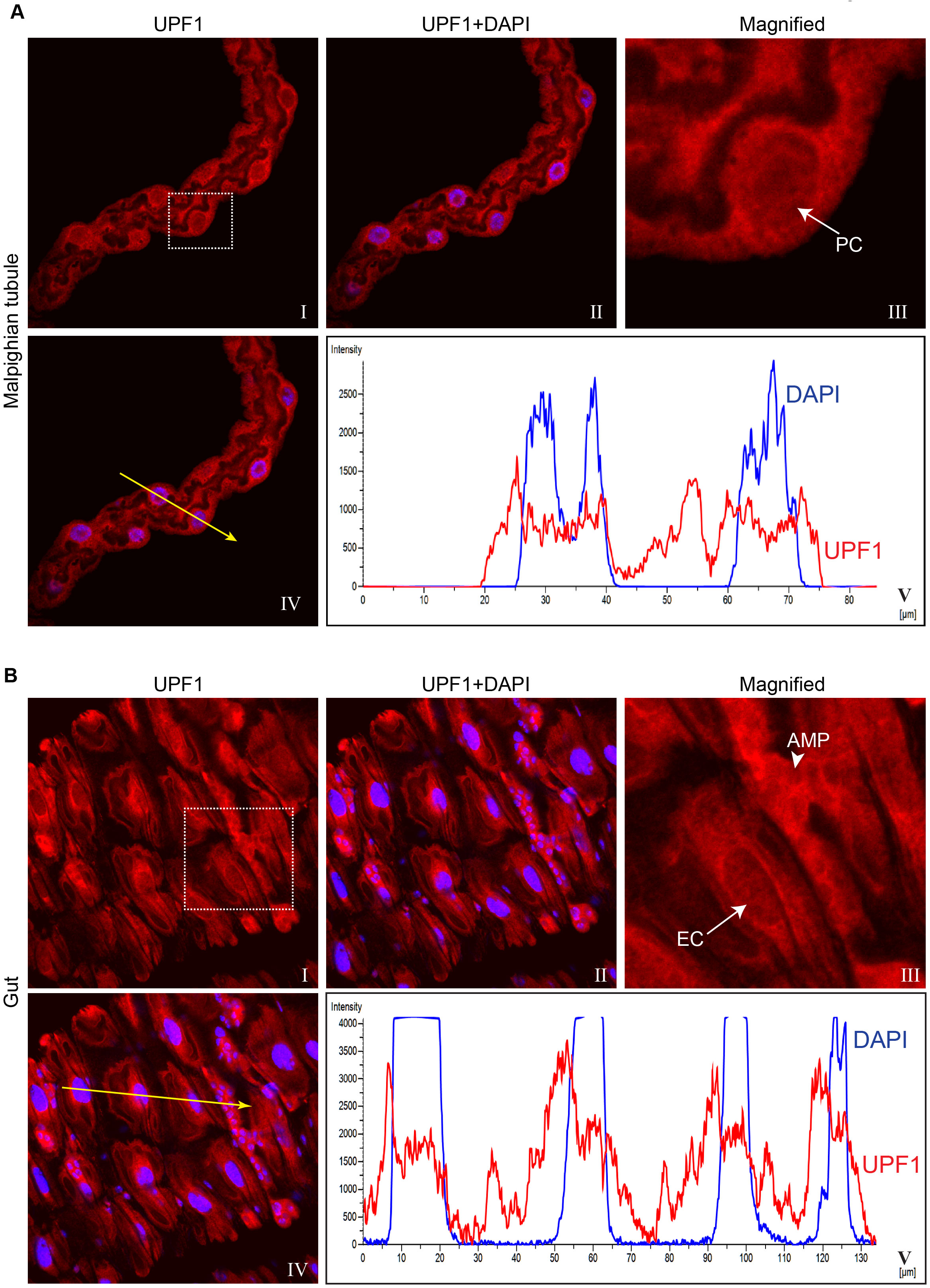
UPF1 subcellular localization in different larval tissues. (**A**) Fluorescence immunolocalization of UPF1 (Cy3, red) in 3^rd^ instar larval Malpighian tubule (I to IV). Panel III shows magnified view of boxed area in panel I, white arrow indicates UPF1 signal within nucleus of a Principal Cell (PC). Tissues were counter-stained with DAPI (blue). Line profiles (V) show both Cy3 and DAPI fluorescence intensities along the yellow line drawn (IV). (**B**) Immunolocalization of UPF1 (Cy3, red) in gut cells (I to IV). Panel III shows magnified view of boxed area panel I. Arrow and arrowhead in (III) indicate presence of UPF1 within nuclei of Enterocytes Cell (EC) and Adult Midgut Progenitor Cells (AMPs), respectively. The line profiles show both Cy3 and DAPI fluorescence intensities along the yellow line (IV) across both EC and AMPs cells (V).

**Figure S3.**
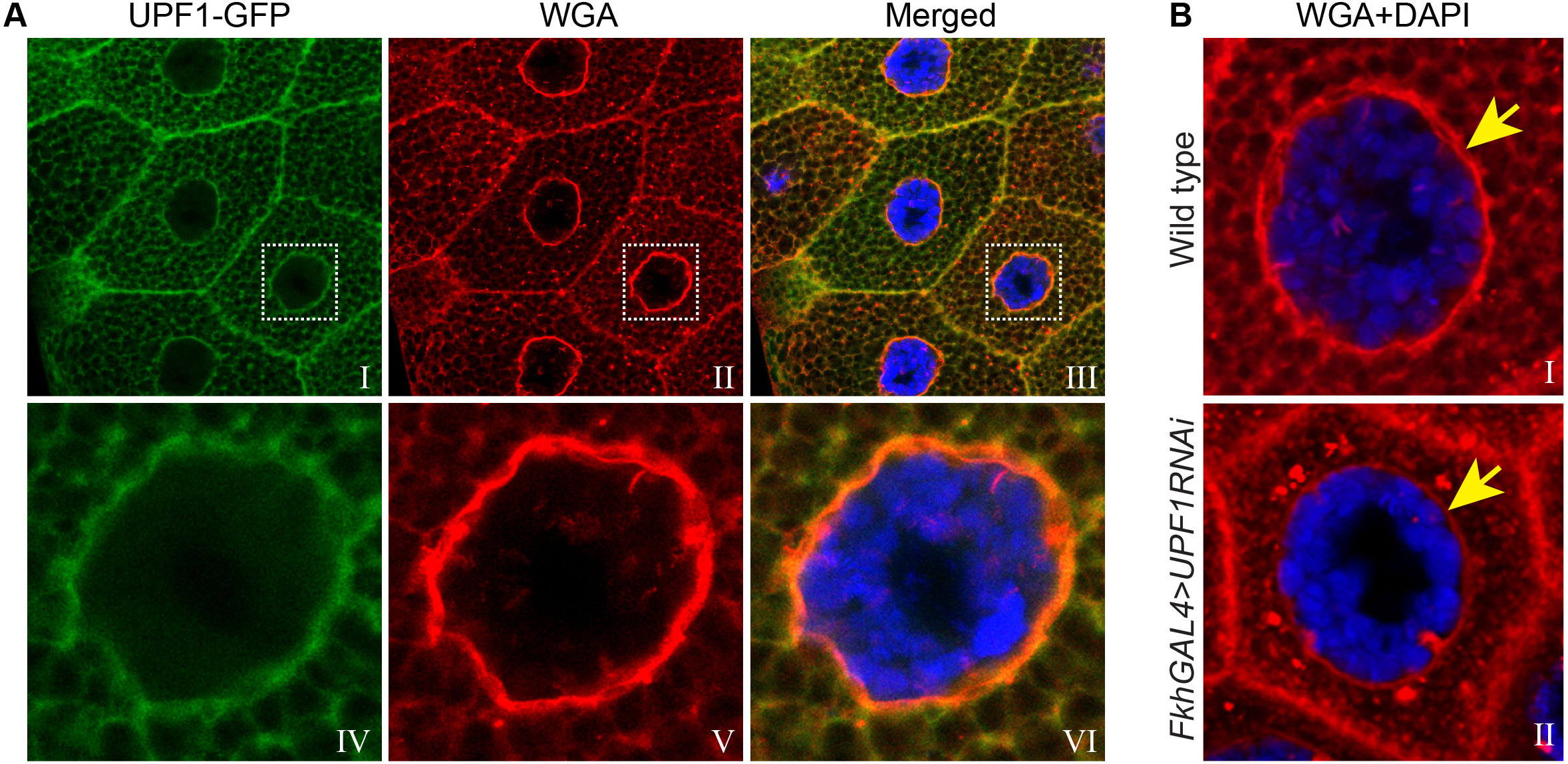
UPF1 may associate with nuclear membrane. (**A**) Fluorescence imaging of tetramethylrhodamine conjugated Wheat Germ Agglutinin (WGA, red) and UPF1-GFP (green) in salivary gland cells. Lower panels are magnified view of boxed area in upper panels. (**B**) Imaging of WGA (red) in wild type (upper panel) and FkhGAL4>UPF1-RNAi (lower panel) salivary gland cells. Yellow arrow indicates nuclear envelope. Cells were counter-stained with DAPI (blue).

**Figure S4.**
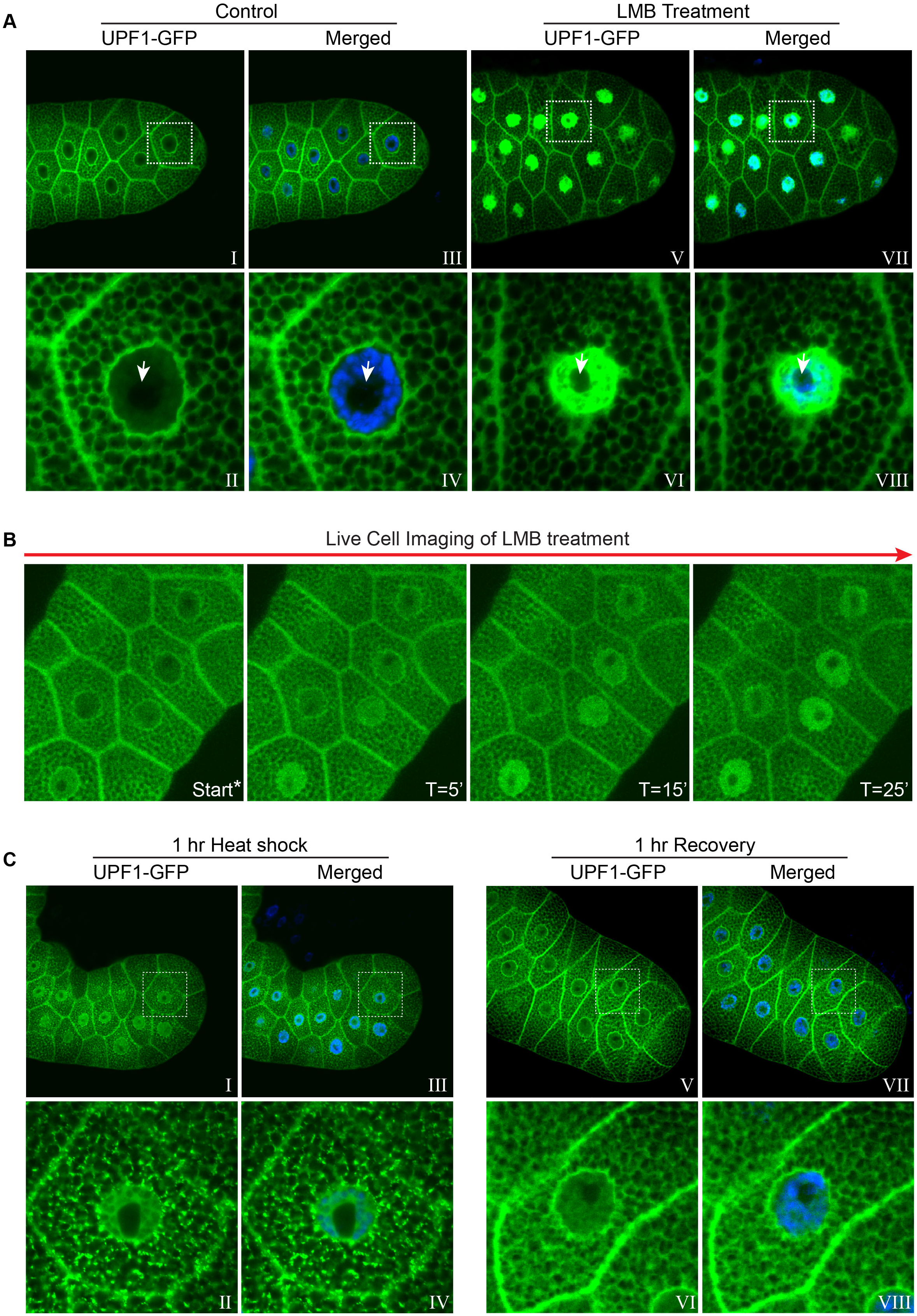
UPF1 is highly dynamic within both nucleus and cytoplasm. (**A**) Imaging of 3^rd^ instar larval salivary glands transgenically expressing UPF1-GFP (green), incubated for 1 hr in either normal M3 media (Control, I to IV) or supplemented with 100µM LMB (LMB, V to VIII). Panels II, IV, VI and VIII are magnified views of the boxed areas in panel I, III, V and VII, respectively. The arrows in II, IV, VI and VIII indicate the nucleoli. Nuclei were counter-stained with DAPI (blue). (**B**) Time-lapse live cell imaging showing changes in the cellular distribution of UPF1-GFP in 3^rd^ instar larval salivary gland at different time intervals of LMB treatment. Start* refers to the first image acquired straight after dissection and mounting of the tissues in a cavity slide, the procedure takes ∼5-6 minutes in which cells have been exposed to LMB. (**C**) Localization of UPF1-GFP (green) in salivary glands after 1 hr heat shock (I to IV) or after 1 hr recovery following heat shock (V to VIII). Lower panels are magnified views of boxed area in upper panel, with or without DAPI counter-staining (blue).

**Figure S5.**
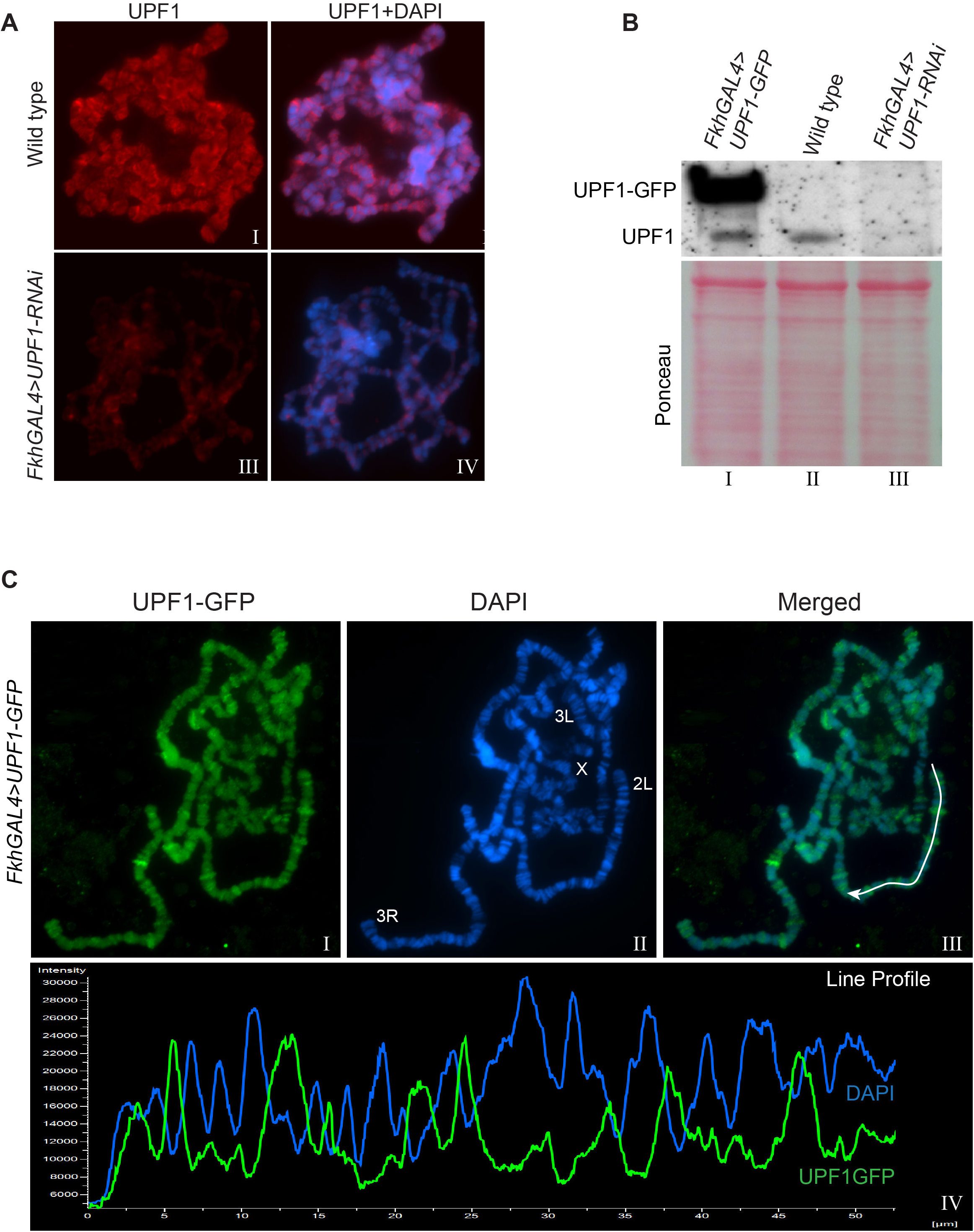
UPF1 RNAi depletes 7B12 mab signal. (**A**) Fluorescence immunolocalization of UPF1 using the 7B12 monoclonal antibody (Cy3, red) on polytene chromosomes (blue) of wild type (I, II) and UPF1-RNAi (III, IV) salivary glands. (**B**) Western blotting probed with 7B12 mab for protein extracts of 3^rd^ instar larval salivary gland from *FkhGAL4>UPF1-GFP* (lane I), wild type (lane II) and *FkhGAL4>UPF1-RNAi* (lane III). Ponceau staining of the same blot showing equal protein loading. (**C**) Immunolocalization of UPF1-GFP (FITC, green, I, III) on polytene chromosomes of *FkhGAL4>UPF1-GFP* salivary glands, detected using anti-GFP antibody. Chromosomes were counter stained with DAPI (blue, II, III). Line profiles in IV show both signal intensities along the white line traced on the chromosome arm in III. Note that UPF1 signal peaks at chromatin-decondensed regions characterised by low DAPI signal.

**Figure S6.**
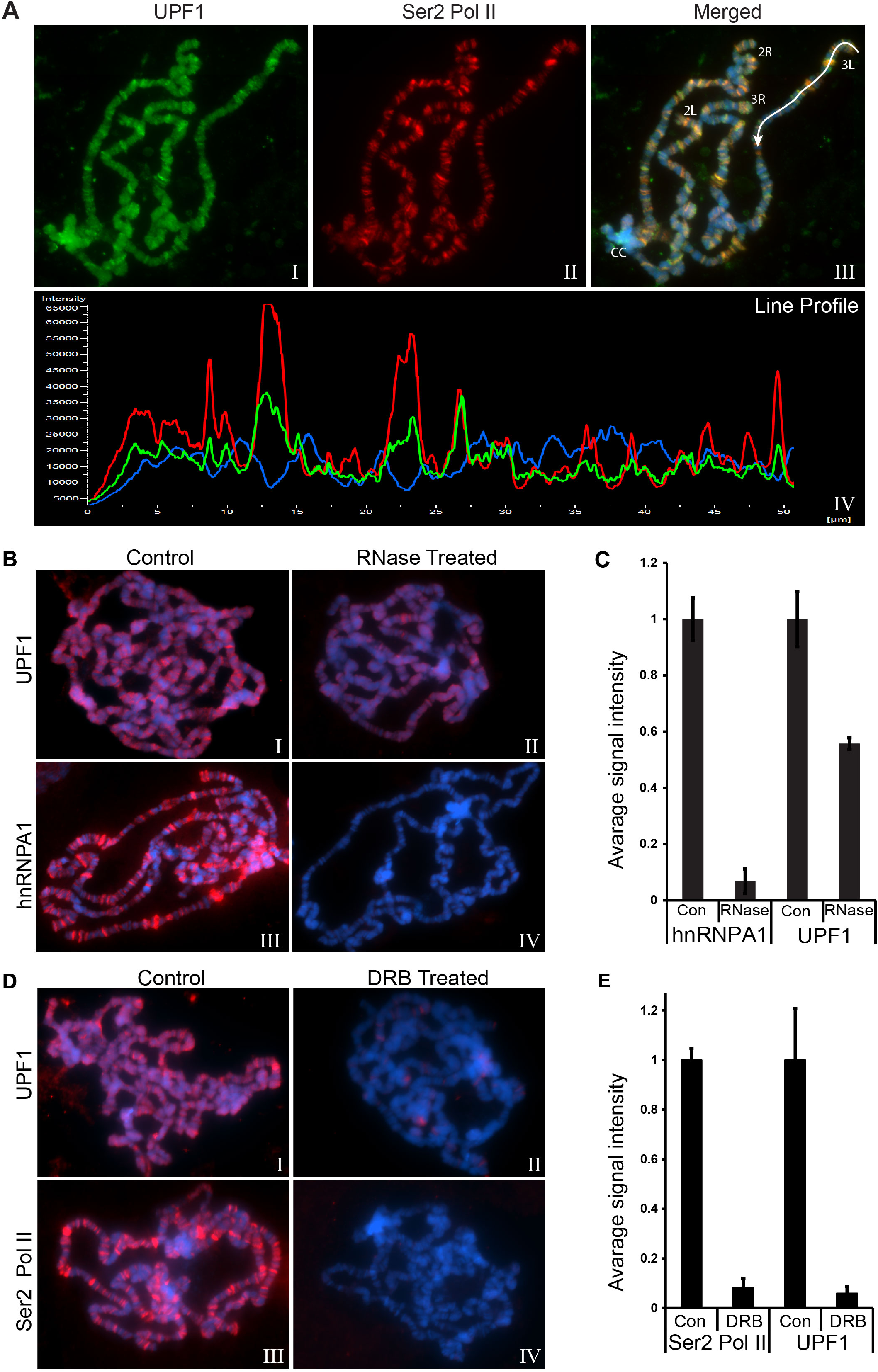
UPF1 chromosomal association is transcription and nascent RNA dependent. (**A**) Co-immunolocalization of UPF1 (FITC, green, I, III) and Ser2 Pol II (Cy3, red, II, III) at polytene chromosomes, counterstained with DAPI (blue). Line profiles in IV show all signal intensities along the white line drawn in III. (**B**) Immunolocalization of UPF1 (red, I and II) and hnRNPA1 (red, III and IV) at polytene chromosomes of either untreated (I and III) or RNase treated salivary glands (II and IV). (**C**) Graph shows normalized fluorescence intensity of the hnRNPA1 and UPF1 signals in control and after RNase treatment, based on mean intensities of 8-10 different nuclear spreads. (**D**) Immunolocalization of UPF1 (red, I and II) and Ser2 Pol II (red, III and IV) at polytene chromosomes from untreated (I and III) or DRB treated glands (II and IV). (**E**) Graph shows normalized fluorescence intensity of Ser2 Pol II and UPF1 in control and after DRB treatment, based on mean intensities of 8 different nuclear spreads.

**Figure S7.**
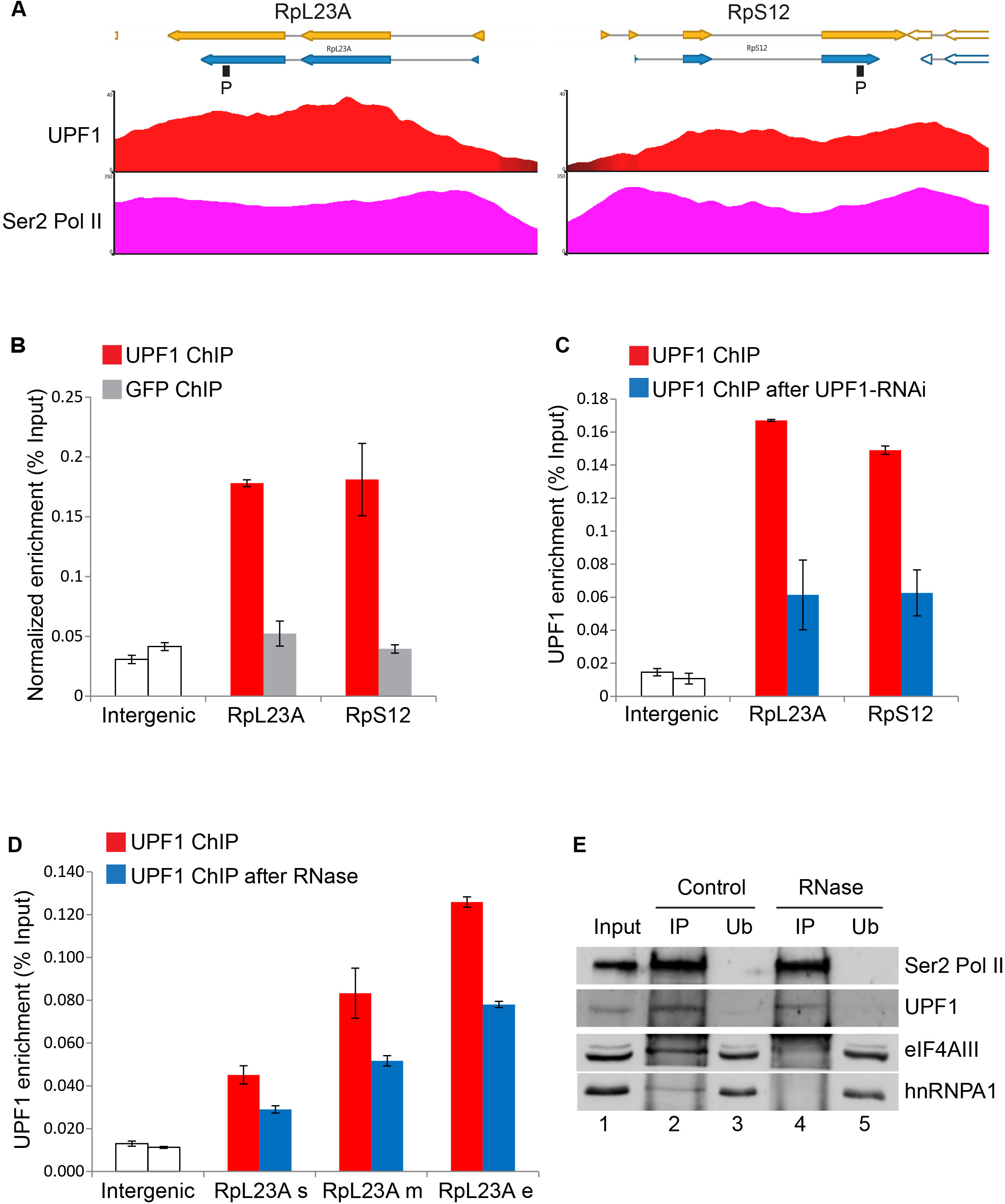
Real-time PCR validation of UPF1 ChIP association at selected genes. (**A**) UPF1 (red) and Ser2 Pol II (pink) ChIP-seq enrichment profiles at RpL23A (on left) and RpS12 (on right) gene loci. (**B**) Real-time PCR quantification of average ChIP signal of either endogenous UPF1 (red) or GFP (as negative control, grey) at the RpL23 and RpS12 genes in salivary glands expressing GFP. The locations of the primer pairs used are indicated by the black boxes (P) shown below the genes schematics in A. (**C**) Real-time PCR quantification of average UPF1 association in control (red) or UPF1-RNAi (blue) S2 cells. (**D**) Real-time PCR quantification of average UPF1 association at three distinct regions of RpL23 in S2 cells (same primers pairs as in Figure 3F) with (blue) or without (red) RNase A treatment. (**E**) Ser2 Pol II immunoprecipitation of S2 cell nuclear extracts using anti-Ser2 Pol II antibody (ab5095) and detection (same blot) of Ser2 Pol II, UPF1, eIF4AIII and hnRNPA1, in control (lanes 2-3) or RNase treated samples (lanes 4-5). IP refers to immunoprecipitated fractions, Ub to unbound fractions.

**Figure S8.**
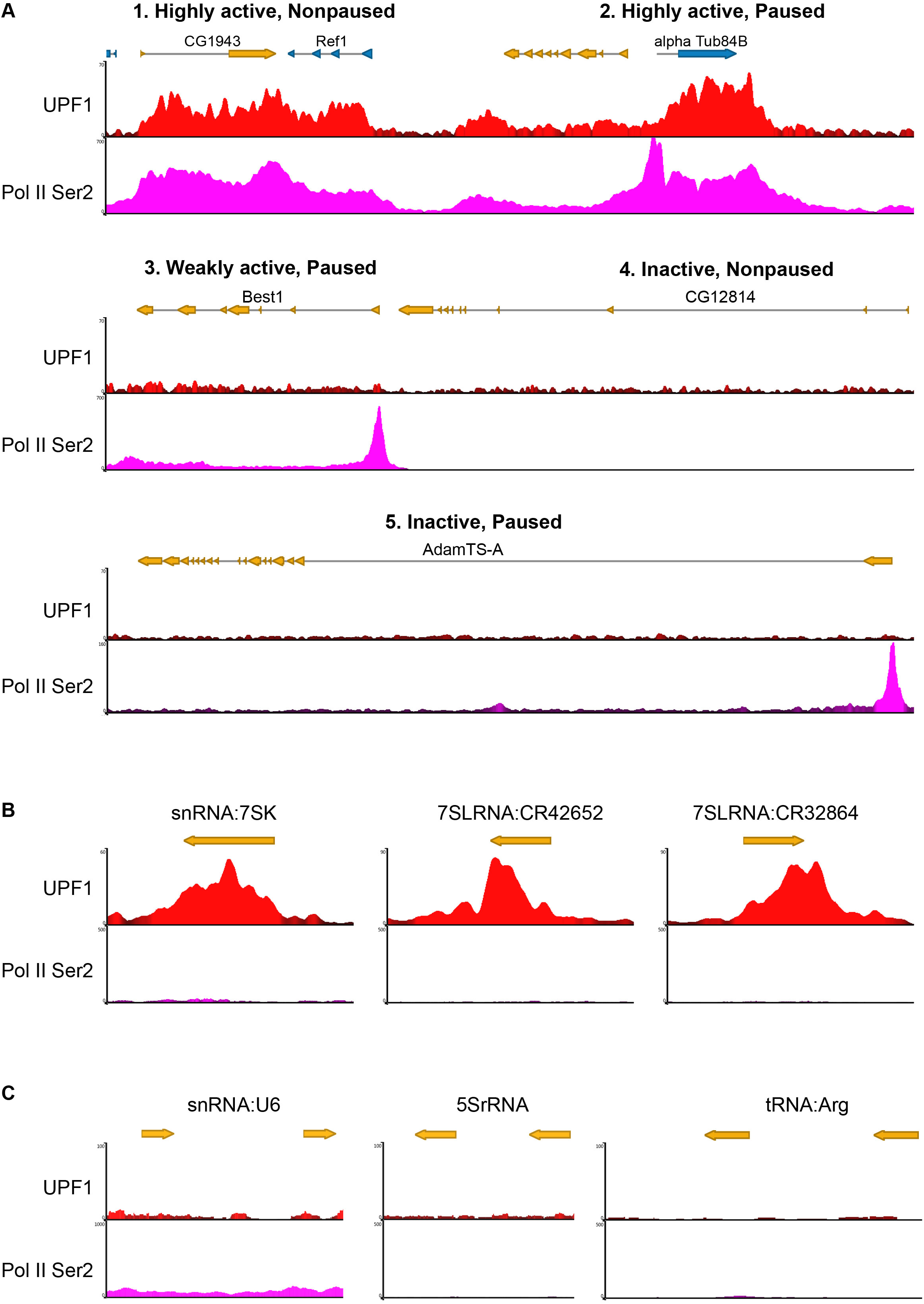
ChIP-seq profiles of UPF1 at representative Pol II genes and some Pol III loci. (**A**) ChIP-seq profiles of UPF1 (red) and Ser2 Pol II (pink) at different active genes characterised by either paused or not paused Pol II at the TSS (panels 1-3). Panels 4 and 5 show absence of Upf1 at two inactive genes, with or without paused Pol II. (**B**) ChIP-seq profile of UPF1 (red) and Ser2 Pol II (pink) at the three Pol III transcribing gene loci indicated. (**C**) ChIP-seq profiles showing no UPF1 signal at three highly transcribing Pol III genes (snRNPU6, 5SrRNA and tRNA:Arg).

**Figure S9.**
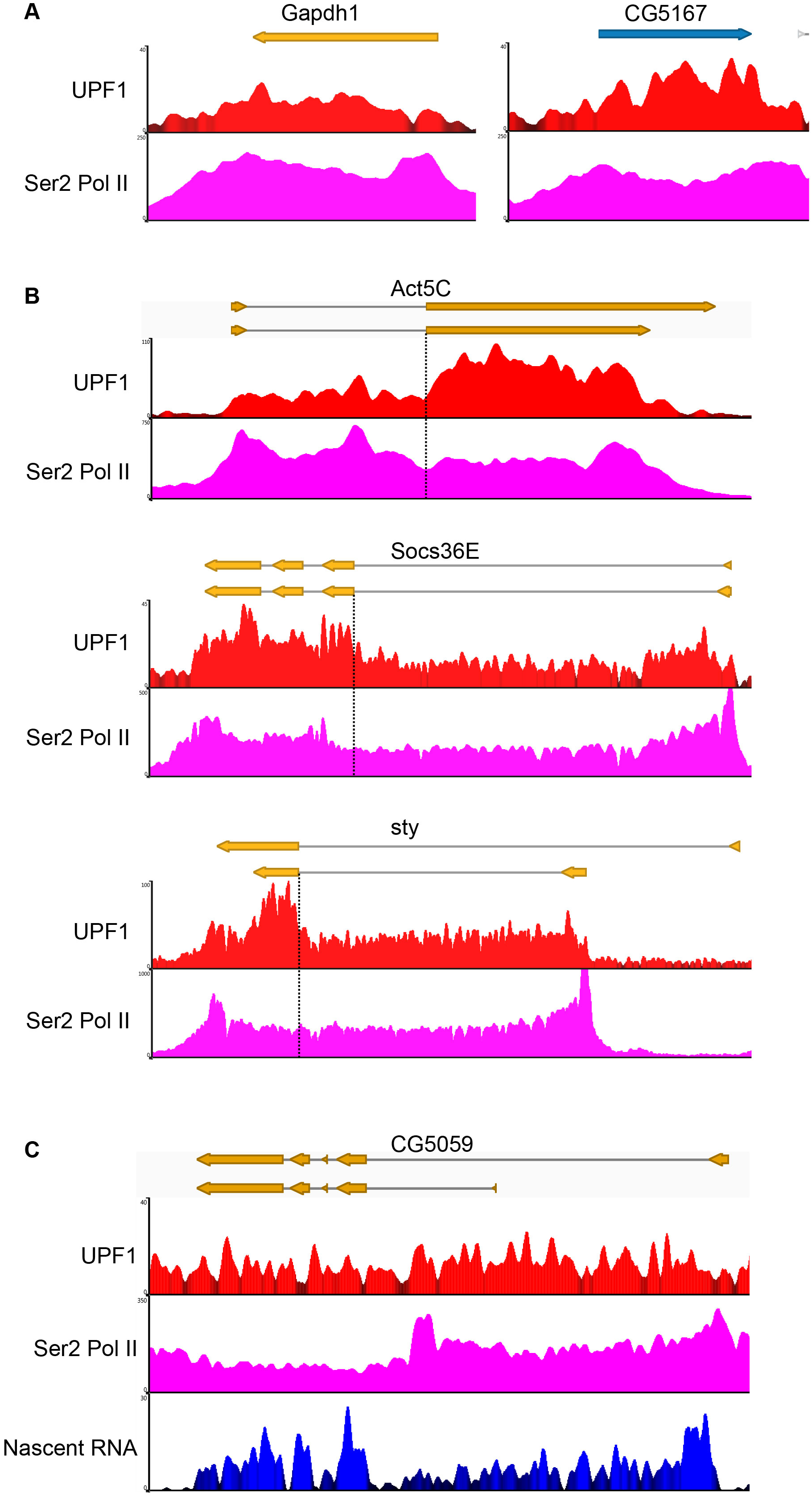
Additional examples of UPF1 ChIP-seq profiles at genes with or without introns. (**A**) ChIP-seq profiles showing enrichment of UPF1 (red) and Ser2 Pol II (pink) at genes without intron. (**B**) ChIP-seq profile showing enrichment of UPF1 (red) and Ser2 Pol II (pink) at other highly expressing intron-containing genes. Dotted lines demarcate regions around intron/exon borders at which UPF1 shows higher association with exons despite uniform Ser2 distribution. (**C**) UPF1 ChIP-seq profile (red) and Ser2 Pol II (pink) at CG5059 gene, which as indicated by the nascent RNA-seq profile underneath (blue) show high intronic sequencing reads, indicative of inefficient co-transcriptional splicing.

**Figure S10.**
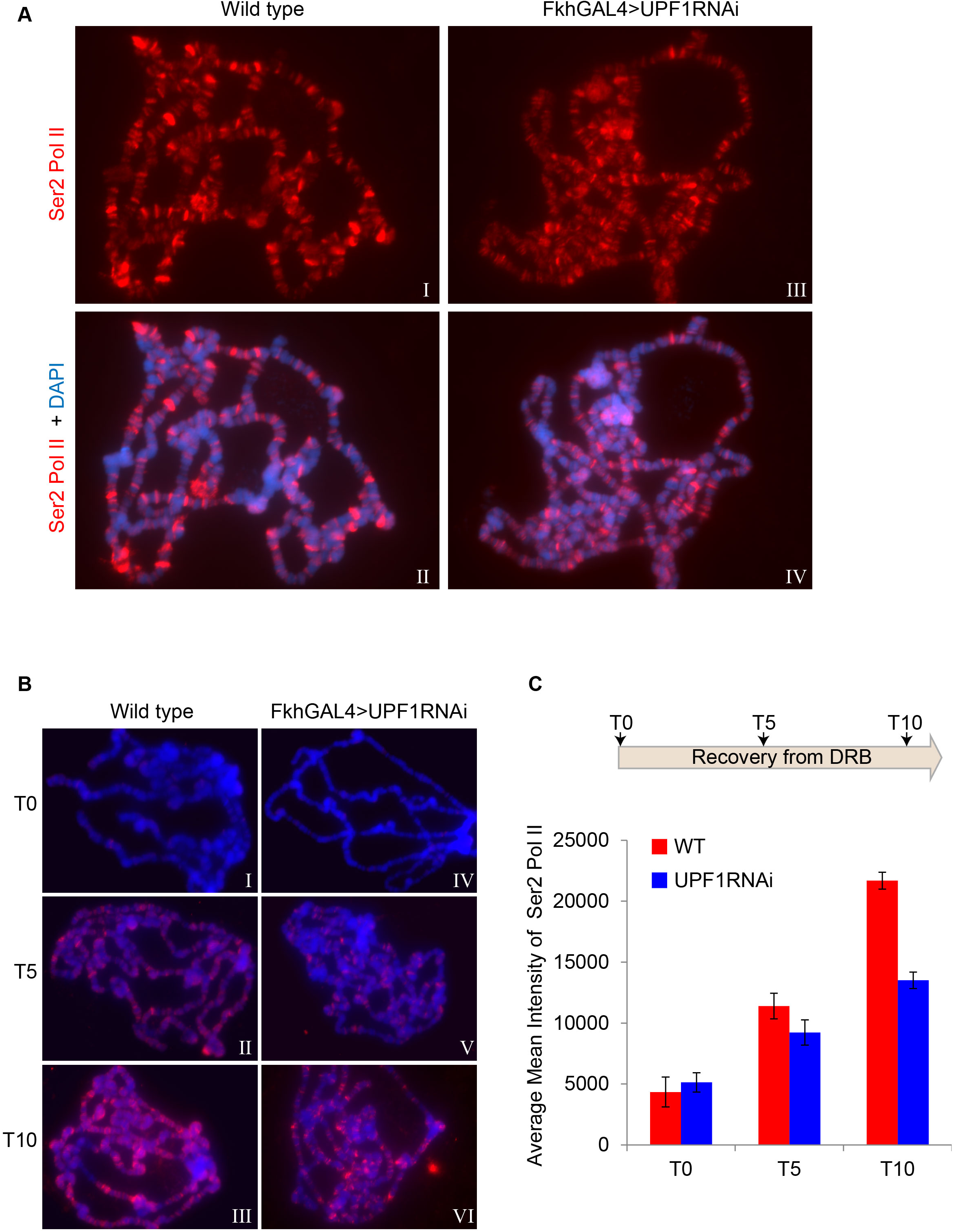
Depletion of UPF1 slows Pol II transcription recovery from DRB treatment. (**A**) Immunolocalization of Ser2 Pol II (Cy3, red) on polytene chromosomes (DAPI, blue) of wild type (I, II) or *FkhGAL4>UPF1-RNAi* (III, IV) 3^rd^ instar larval salivary glands. (**B**) Immunolocalization of Ser2 Pol II (Cy3, red) on polytene chromosomes (DAPI, blue) of wild type (I-III) or *FkhGAL4>UPF1-RNAi* (IV-VI) salivary glands, at different time points after recovery from DRB treatment for 0 min (T0), 5 min (T5) and 10 min (T10). (**C**) Top, schematics of the timeframe of the recovery from DRB treatment in S2 cells. Panel below shows average mean intensity of Ser2 Pol II signal on polytene chromosomes of wild type (red) or UPF1-RNAi (blue) salivary glands treated with DRB and at two time points after its removal. The mean of intensity was calculated by measuring signals in 10 different chromosome spreads.

**Table S1.** Peptides used for UPF1 antibody generation (positions refer to full length sequence FBpp0073433)

|  |  |  |  |
| --- | --- | --- | --- |
| No. | Start | End | Peptide |
| 1 | 43 | 52 | TSQSQTQNDQ |
| 2 | 85 | 94 | DEPGSSYVKE |
| 3 | 1084 | 1093 | PGGNKKTNKL |
| 4 | 342 | 351 | HYVGELYNPW |
| 5 | 788 | 797 | QYQGSLHSRL |
| 6 | 27 | 36 | DTQPTQYDYR |
| 7 | 528 | 537 | KSREAIDSPV |
| 8 | 60 | 69 | SAGDSHPRLA |
| 9 | 1140 | 1149 | SQQPELSQDF |
| 10 | 259 | 268 | KPGIDSEPAH |
| 11 | 1060 | 1069 | QTGNFSPGNS |
| 12 | 1117 | 1126 | AAPYSQHPMP |

**Table S2.** Most enriched transcription units by UPF1 ChIP-seq (black=Protein coding genes, red=Noncoding RNAs).

| No. | Transcript | Chromosome | Cytological Location | Strand | Gene | UPF1 Enrichment |
| --- | --- | --- | --- | --- | --- | --- |
| 1 | NR_002554 | chr2R | 60A3 | - | snoRNA:Psi18S-176 | 20.92784248 |
| 2 | NR_073860 | chr2R | 45F1 | + | CR43651 | 19.84962099 |
| 3 | NR_073861 | chr2R | 45F1 | + | CR43651 | 19.58870612 |
| 4 | NM_078901 | chr2R | 42A7 | - | Act42A | 16.53573972 |
| 5 | NM_166890 | chrX | 2B1 | - | sta | 15.7976555 |
| 6 | NM_001297856 | chrX | 2B1 | - | sta | 14.65188523 |
| 7 | NM_057402 | chrX | 2B1 | - | sta | 14.65188523 |
| 8 | NM_001316458 | chr3L | 70A6 | + | Nplp2 | 14.2141182 |
| 9 | NM_144173 | chr3L | 70A6 | + | Nplp2 | 14.2141182 |
| 10 | NM_166889 | chrX | 2B1 | - | sta | 14.0181416 |
| 11 | NM_206272 | chr3L | 63F5 | - | Ubi-p63E | 13.10522634 |
| 12 | NR_001992 | chr3R | 84A5 | - | 7SLRNA:CR32864 | 13.04190544 |
| 13 | NR_037753 | chr3R | 84A5 | - | 7SLRNA:CR42652 | 13.04190544 |
| 14 | NM_001274454 | chr3L | 63F5 | - | Ubi-p63E | 12.94301365 |
| 15 | NM_168043 | chr3L | 63F5 | - | Ubi-p63E | 12.86980148 |
| 16 | NM_079185 | chr3L | 63F5 | - | Ubi-p63E | 12.8624403 |
| 17 | NM_001299392 | chr2R | 48C5 | + | Ef1alpha48D | 12.7336405 |
| 18 | NR_002067 | chr3R | 93D | + | Hsromega | 12.6383098 |
| 19 | NR_002069 | chr3R | 93D | + | Hsromega | 12.6383098 |
| 20 | NM_001300130 | chr3L | 70A6 | + | Nplp2 | 12.582198 |
| 21 | NM_001316502 | chr3R | 86D8 | + | Tctp | 11.88147055 |
| 22 | NM_058027 | chr2R | 48C5 | + | Ef1alpha48D | 11.51812571 |
| 23 | NM_001014455 | chr2L | 21 E2 | + | PNUTS | 11.50911291 |
| 24 | NR_048444 | chr3R | 93D | + | Hsromega | 11.47018513 |
| 25 | NM_001297855 | chrX | 2B1 | + | rush | 11.45428524 |
| 26 | NM_130567 | chrX | 2B1 | + | rush | 11.36293998 |
| 27 | NM_170070 | chr3R | 94 E10-94 13 | - | pnt | 11.36070171 |
| 28 | NM_141791 | chr3R | 86D8 | + | Tctp | 11.3160895 |
| 29 | NM_001298659 | chr2L | 23F3-23F6 | + | Thor | 11.29086613 |
| 30 | NM_057284 | chr3R | 93 E13 | + | RpS3 | 10.9579823 |
| 31 | NM_001014454 | chr2L | 21 E2 | + | PNUTS | 10.86549487 |
| 32 | NM_001298210 | chrX | 10-E3 to 10 E4 | - | Hsc70-3 | 10.85467784 |
| 33 | NR_133429 | chr3R | 93D | + | Hsromega | 10.74841913 |
| 34 | NM_167307 | chrX | 10-E3 to 10 E4 | - | Hsc70-3 | 10.70276539 |
| 35 | NM_165850 | chr2R | 48C5 | + | Ef1alpha48D | 10.69805745 |
| 36 | NM_001202085 | chr3L | 61B2 | - | CG42846 | 10.5511492 |
| 37 | NM_001275943 | chr3R | 94 E10-94 13 | - | pnt | 10.42349043 |
| 38 | NM_079983 | chr2R | 57C3 | + | Xbp1 | 10.40232605 |
| 39 | NM_166427 | chr2R | 57C3 | + | Xbp1 | 10.39685831 |
| 40 | NM_001202063 | chr2R | 59B6 | - | CG3800 | 10.39235699 |
| 41 | NM_079175 | chr3L | 63B11 | + | Hsp83 | 10.35797394 |
| 42 | NM_001299393 | chr2R | 48C5 | + | Ef1alpha48D | 10.3424767 |
| 43 | NM_143714 | chrX | 5D2 | - | mab-21 | 10.22106132 |
| 44 | NM_167180 | chrX | 8C14 | - | His3.3B | 10.19996373 |
| 45 | NM_057947 | chr2L | 23F3-23F6 | + | Thor | 10.10779488 |
| 46 | NM_001103474 | chrX | 9D2 | - | spri | 10.09939045 |
| 47 | NM_001272490 | chrX | 9D2 | - | spri | 10.0968423 |
| 48 | NM_001272491 | chrX | 9D2 | - | spri | 10.0968423 |
| 49 | NM_001298516 | chrX | 18D3 | + | RpS10b | 10.05584386 |
| 50 | NM_001274433 | chr3L | 63B11 | + | Hsp83 | 9.962334214 |
| 51 | NM_167065 | chrX | 5 E4 | - | Ubi-p5E | 9.845904641 |
| 52 | NM_167306 | chrX | 10 E3-10 E4 | - | Hsc70-3 | 9.739493741 |
| 53 | NM_167308 | chrX | 10 E3-10 E4 | - | Hsc70-3 | 9.739493741 |
| 54 | NM_206223 | chr3L | 61B2 | + | CG33229 | 9.726778964 |
| 55 | NM_169625 | chr3R | 88 E4 | + | Hsc70-4 | 9.714744345 |
| 56 | NM_001300552 | chr3R | 94 E13 | + | RpS3 | 9.705903021 |
| 57 | NM_176503 | chr3R | 88 E4 | + | Hsc70-4 | 9.687864086 |
| 58 | NM_001347750 | chrX | 2B1 | + | mei-38 | 9.661728876 |
| 59 | NM_078497 | chrX | 5C7 | + | Act5C | 9.593373151 |
| 60 | NM_168070 | chr3L | 64A10 | - | ImpL2 | 9.571757664 |
| 61 | NM_134543 | chrX | 19C5 | + | I(1)G0004 | 9.506154816 |
| 62 | NM_169841 | chr3R | 91D3 | + | Xrp1 | 9.489137734 |
| 63 | NM_079879 | chr4 | 101 F1 | - | RpS3A | 9.472988193 |
| 64 | NM_166714 | chr4 | 101 F1 | - | RpS3A | 9.472988193 |
| 65 | NM_001275657 | chr3R | 88 E4 | + | Hsc70-4 | 9.464906214 |
| 66 | NM_001298008 | chrX | 5 E4 | - | Ubi-p5E | 9.44989544 |
| 67 | NM_169626 | chr3R | 88 E4 | + | Hsc70-4 | 9.391845758 |
| 68 | NM_176502 | chr3R | 88 E4 | + | Hsc70-4 | 9.391845758 |
| 69 | NM_176719 | chrX | 8C14 | - | His3.3B | 9.373510295 |
| 70 | NM_169627 | chr3R | 88 E4 | + | Hsc70-4 | 9.333994534 |
| 71 | NM_137895 | chr2R | 59B6 | - | CG3800 | 9.317056284 |
| 72 | NM_001272174 | chrX | 1C5 | + | Sec22 | 9.263596844 |
| 73 | NM_130507 | chrX | 1C5 | + | Sec22 | 9.263596844 |
| 74 | NM_164617 | chr2L | 25C1 | + | Col4a1 | 9.208538686 |
| 75 | NM_057786 | chrX | 1C4 | - | RpL22 | 9.203239849 |
| 76 | NM_164615 | chr2L | 25C1 | + | Col4a1 | 9.190852727 |
| 77 | NM_165869 | chr2R | 48 E8-48 E9 | - | RpS11 | 9.137225506 |
| 78 | NM_001014453 | chr2L | 21 E2 | + | PNUTS | 9.022927536 |
| 79 | NM_001272921 | chr2L | 21 E2 | + | PNUTS | 9.022927536 |
| 80 | NM_164616 | chr2L | 25C1 | + | Col4a1 | 9.013677552 |
| 81 | NM_001275501 | chr3R | 85 E1 | - | Calr | 9.013384298 |
| 82 | NM_001014725 | chrX | 5C7 | + | Act5C | 8.989781735 |
| 83 | NM_132656 | chrX | 11F1 | + | CG1673 | 8.981761268 |
| 84 | NM_079569 | chr3R | 85 E1 | - | Calr | 8.981419499 |
| 85 | NM_079632 | chr3R | 88 E4 | + | Hsc70-4 | 8.957987156 |
| 86 | NM_001273288 | chr2L | 85 E1 | - | Rack1 | 8.945814985 |
| 87 | NM_057921 | chr2L | 85 E1 | - | Rack1 | 8.945814985 |
| 88 | NM_132800 | chrX | 13C4 | - | CG15642 | 8.919897052 |
| 89 | NM_001014726 | chrX | 5C7 | + | Act5C | 8.909849299 |
| 90 | NM_001299958 | chr3L | 61B2 | + | CG33229 | 8.850750946 |
| 91 | NM_079355 | chr3L | 71B5 | - | Pdi | 8.830574597 |
| 92 | NM_080366 | chr2L | 39 E7 | - | EF2 | 8.804229445 |
| 93 | NM_001298794 | chr2L | 85 E1 | - | Rack1 | 8.793822499 |
| 94 | NM_165395 | chr2L | 39 E7 | - | EF2 | 8.793744101 |
| 95 | NM_165394 | chr2L | 39 E7 | - | EF2 | 8.732148442 |
| 96 | NR_123764 | chrX | 1C4 | + | CR44965 | 8.355359116 |
| 97 | NR_002545 | chr2R | 54A2 | + | snoRNA:U3:54Ab | 8.034844038 |
| 98 | NR_002544 | chr2R | 54A2 | + | snoRNA:U3:54Aa | 7.685229313 |
| 99 | NR_002493 | chr3R | 84D4 | - | snRNA:7SK | 7.602249655 |
| 100 | NR_073616 | chr4 | 102B7 | + | CR44027 | 7.389972024 |
| 101 | NR_133116 | chr2R | 50A1-A3 | - | CR44206 | 7.36480527 |
| 102 | NR_047918 | chr2L | 27F4 | + | mir-305 | 7.207345421 |

**Table S3.**
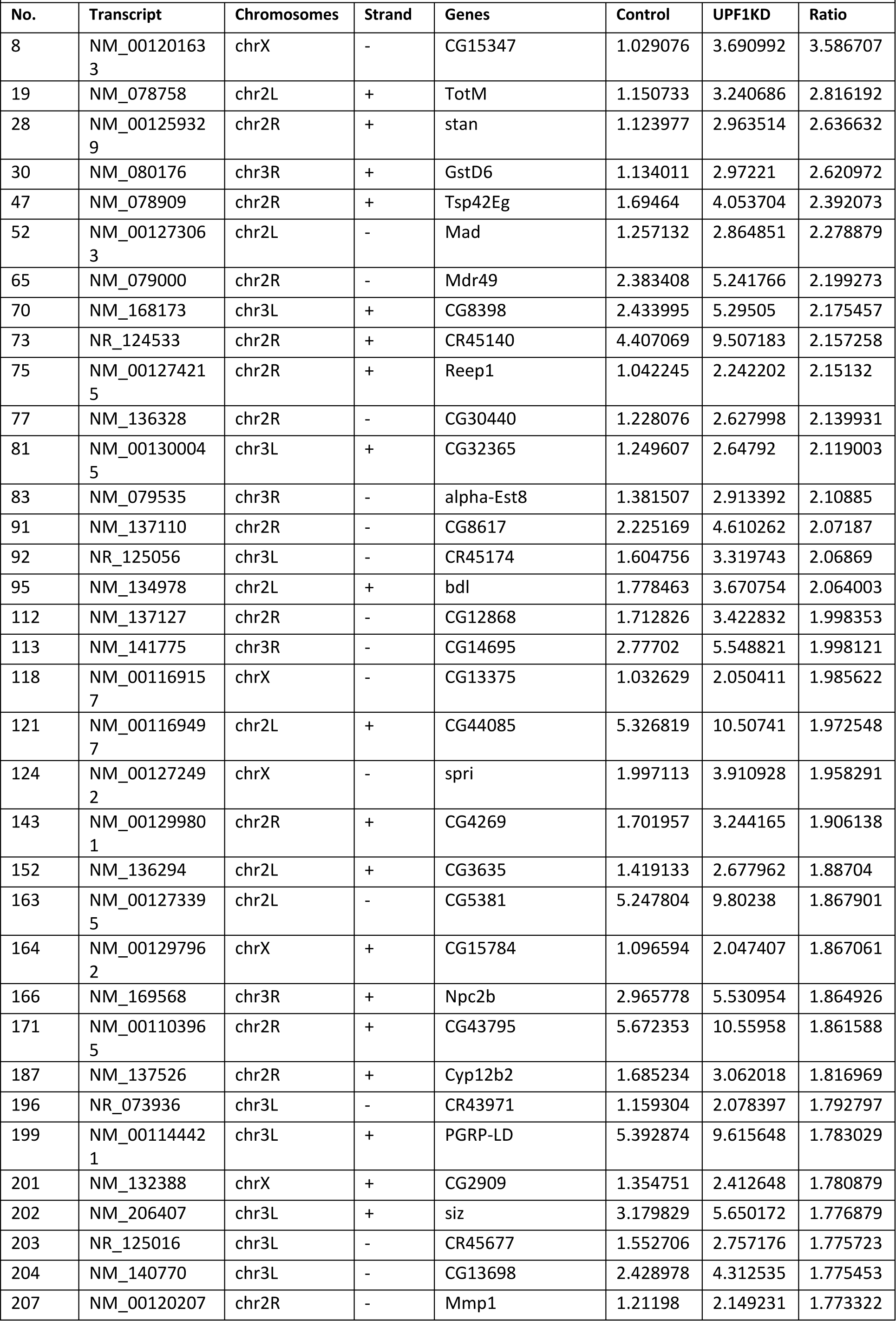
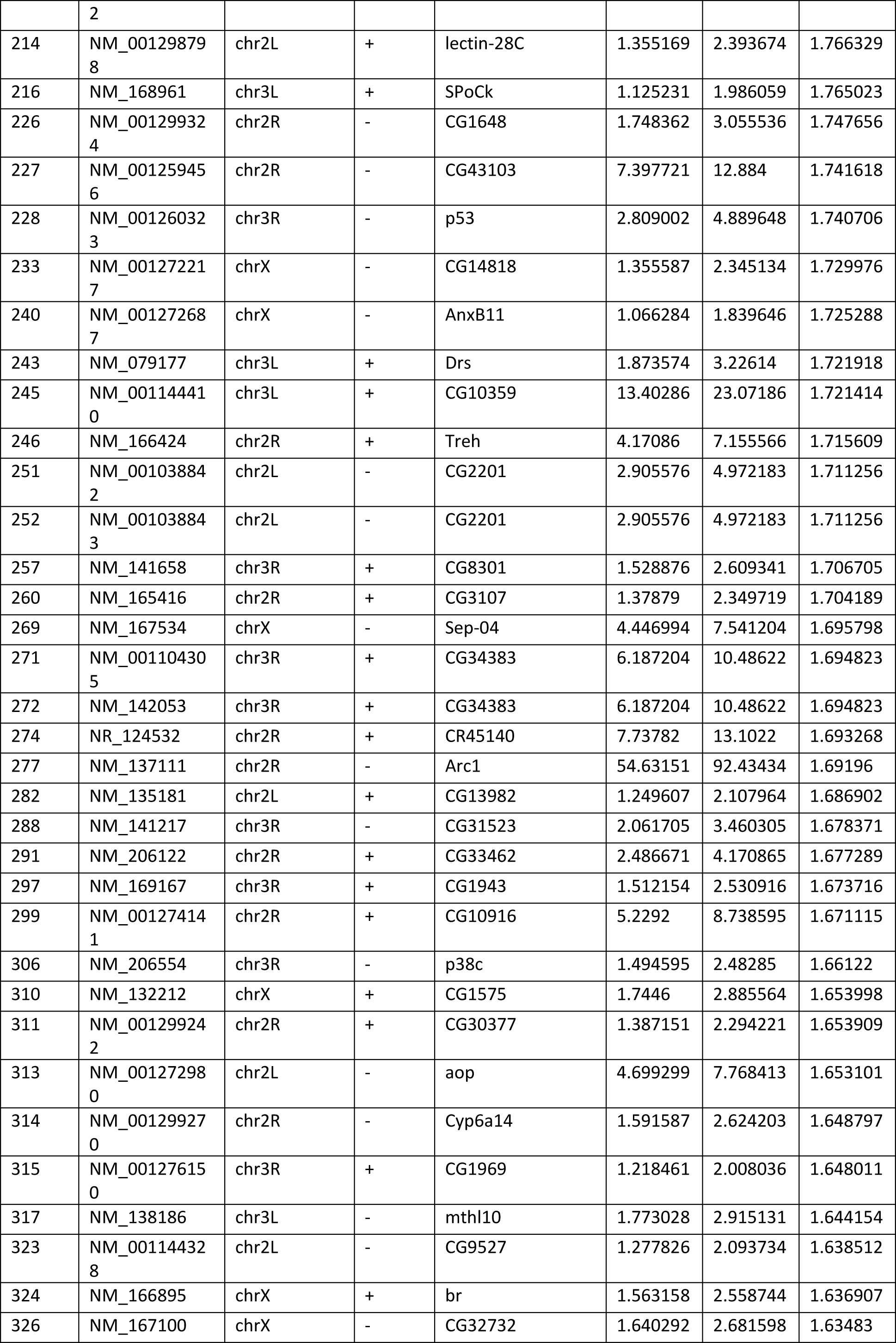
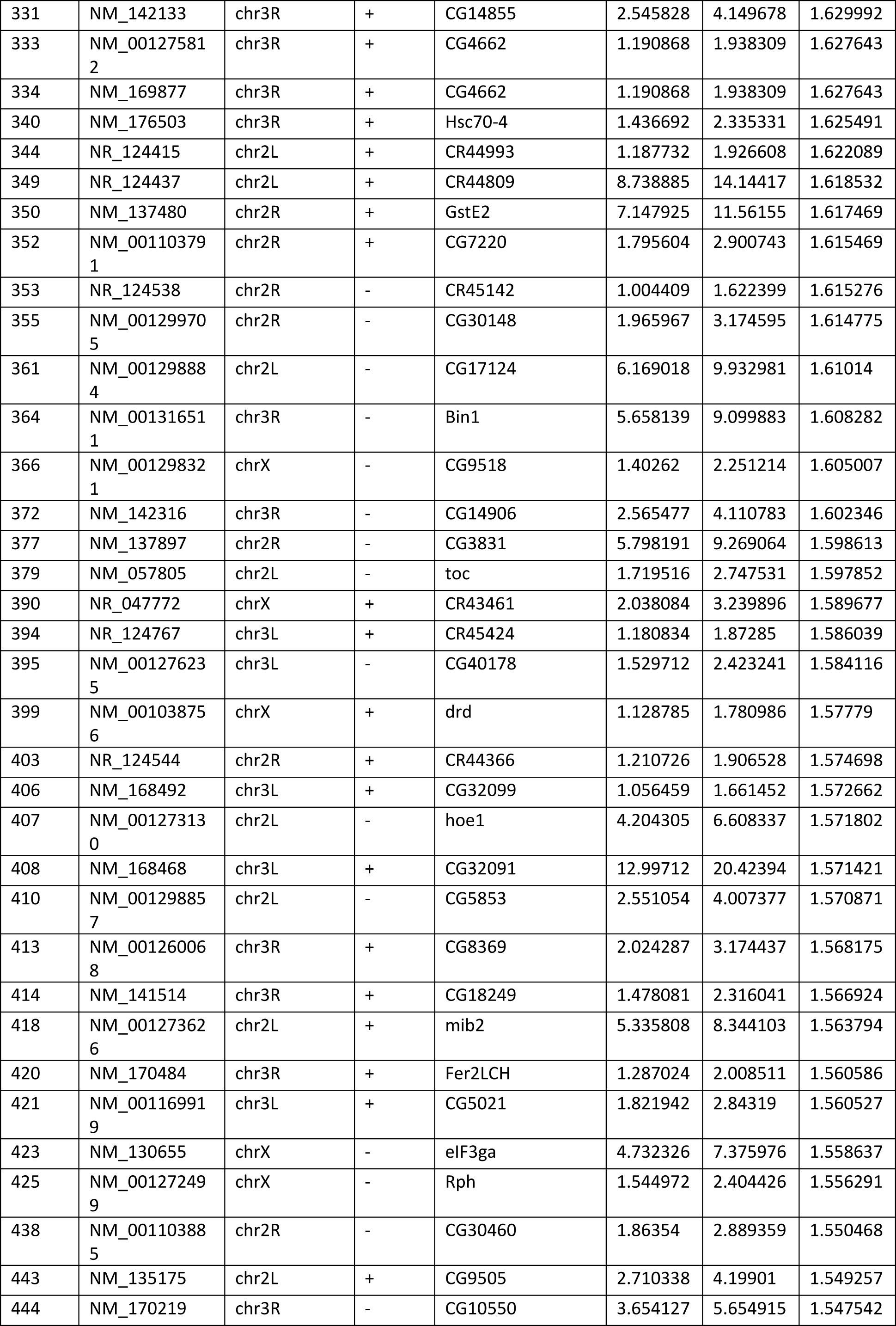
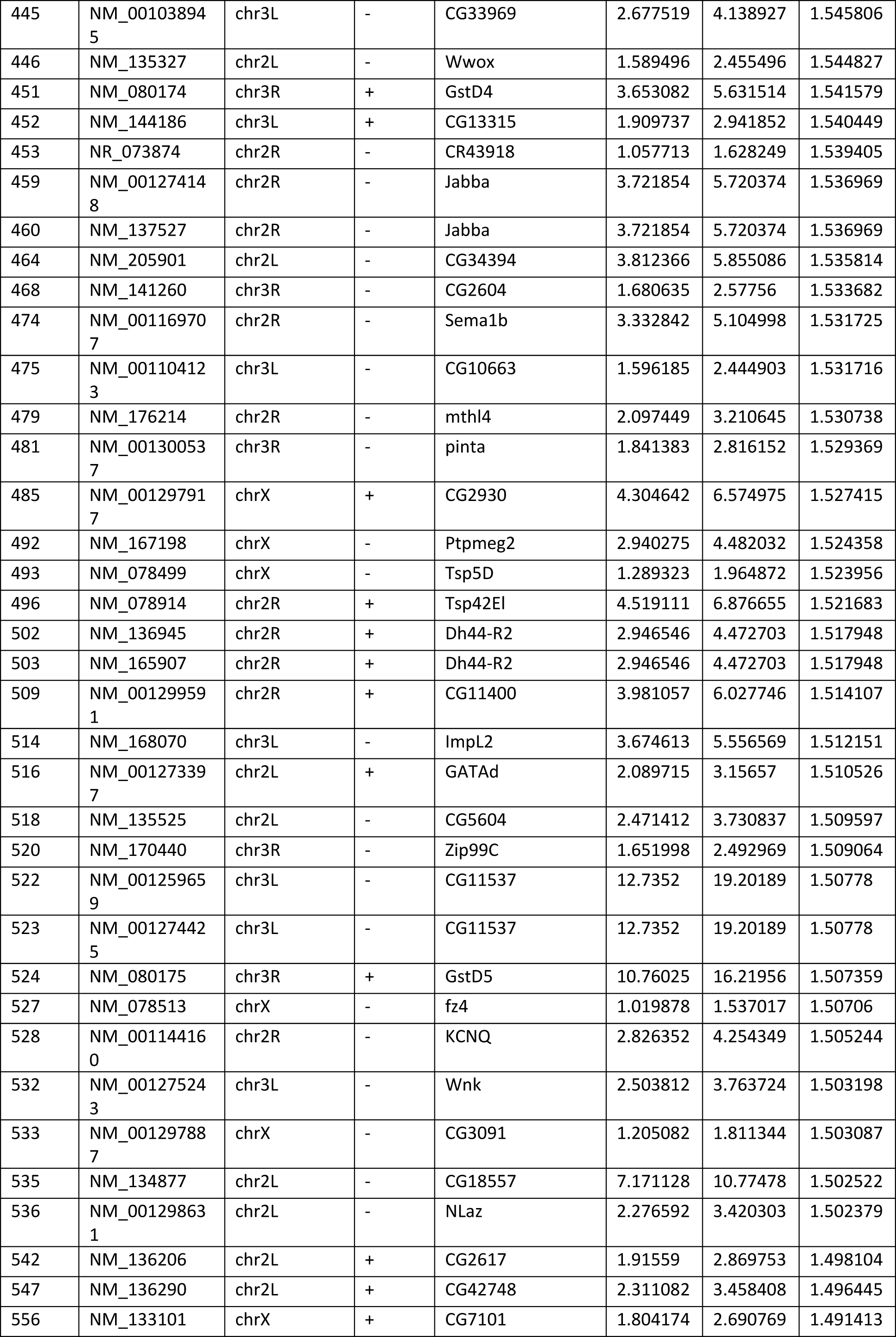
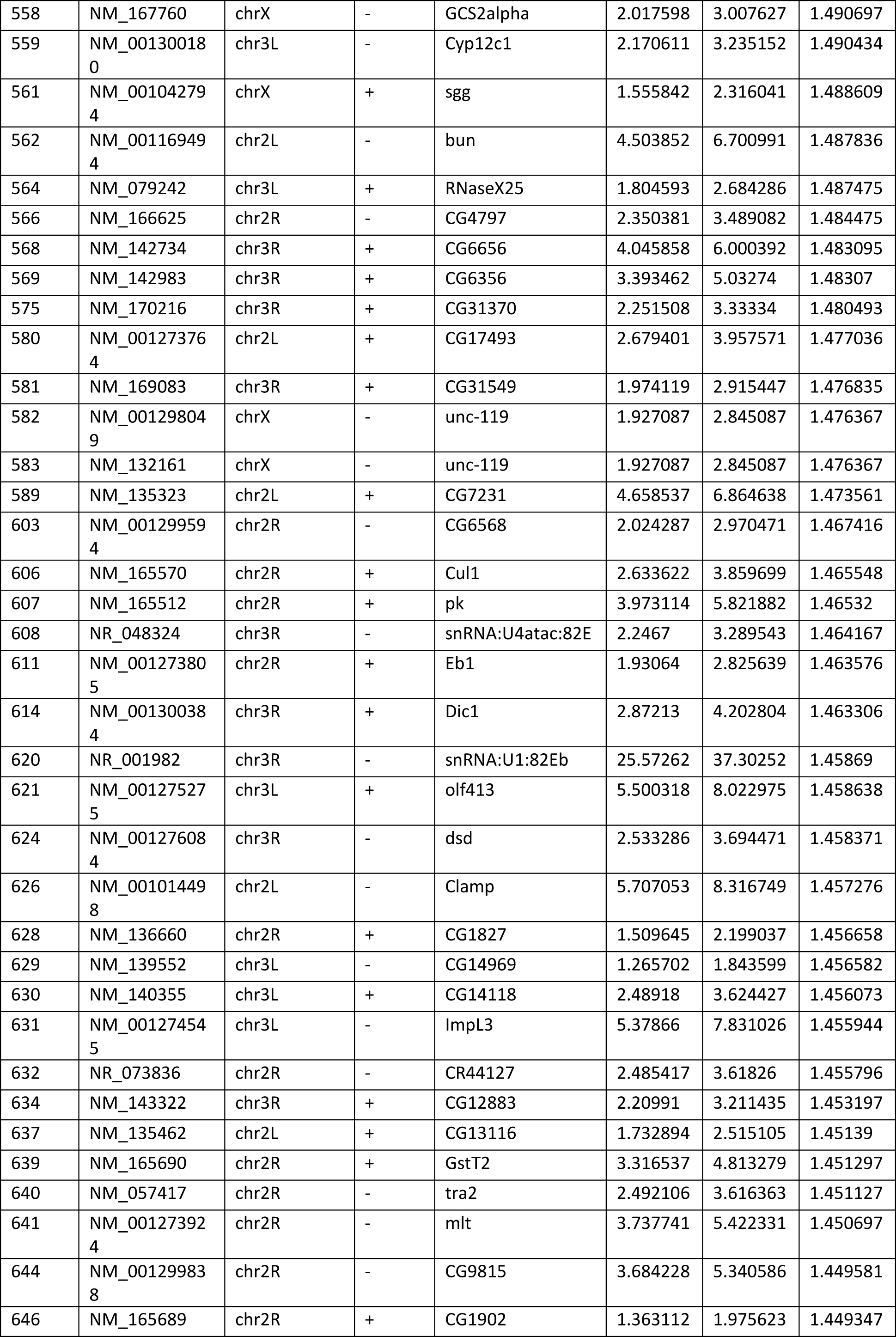
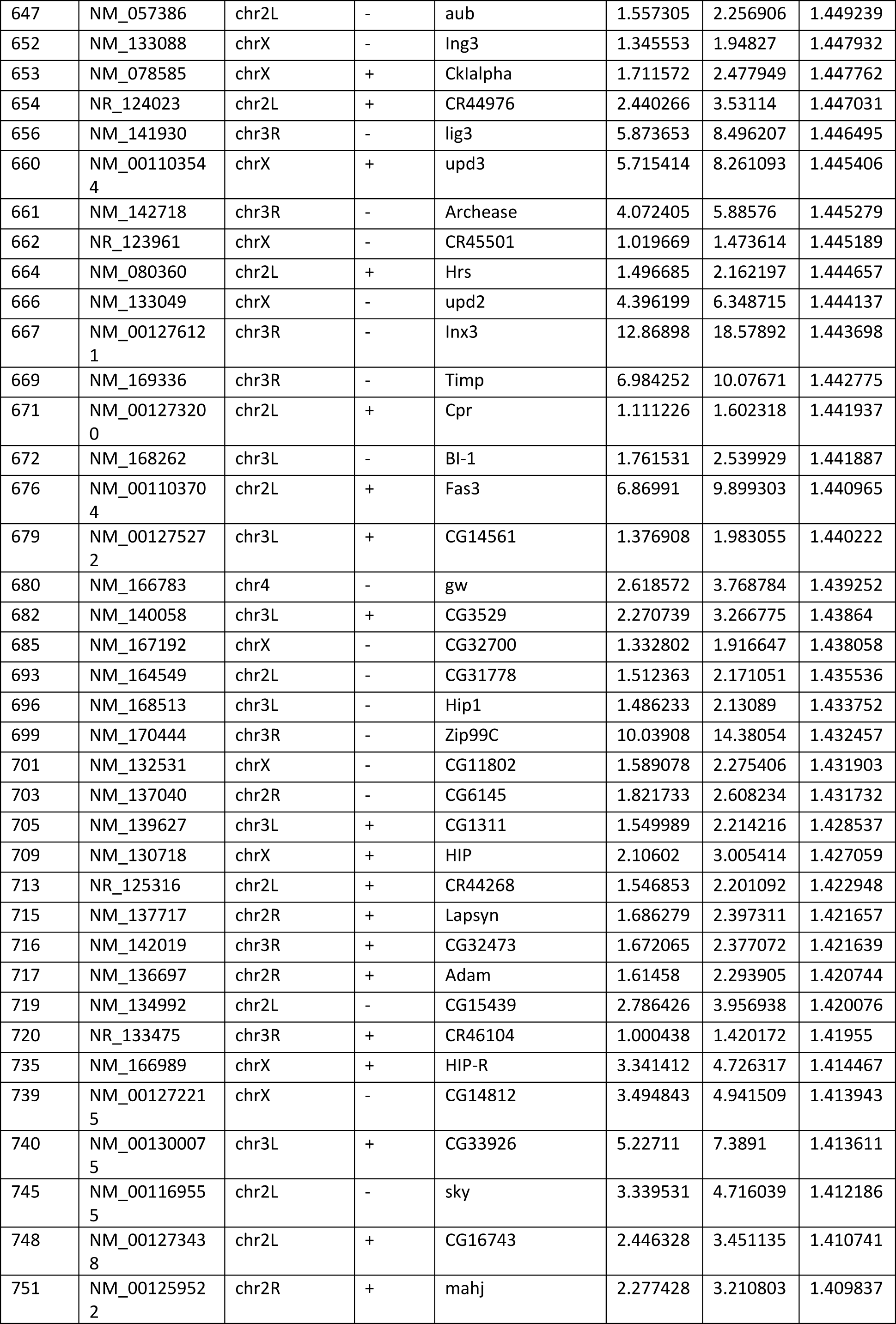
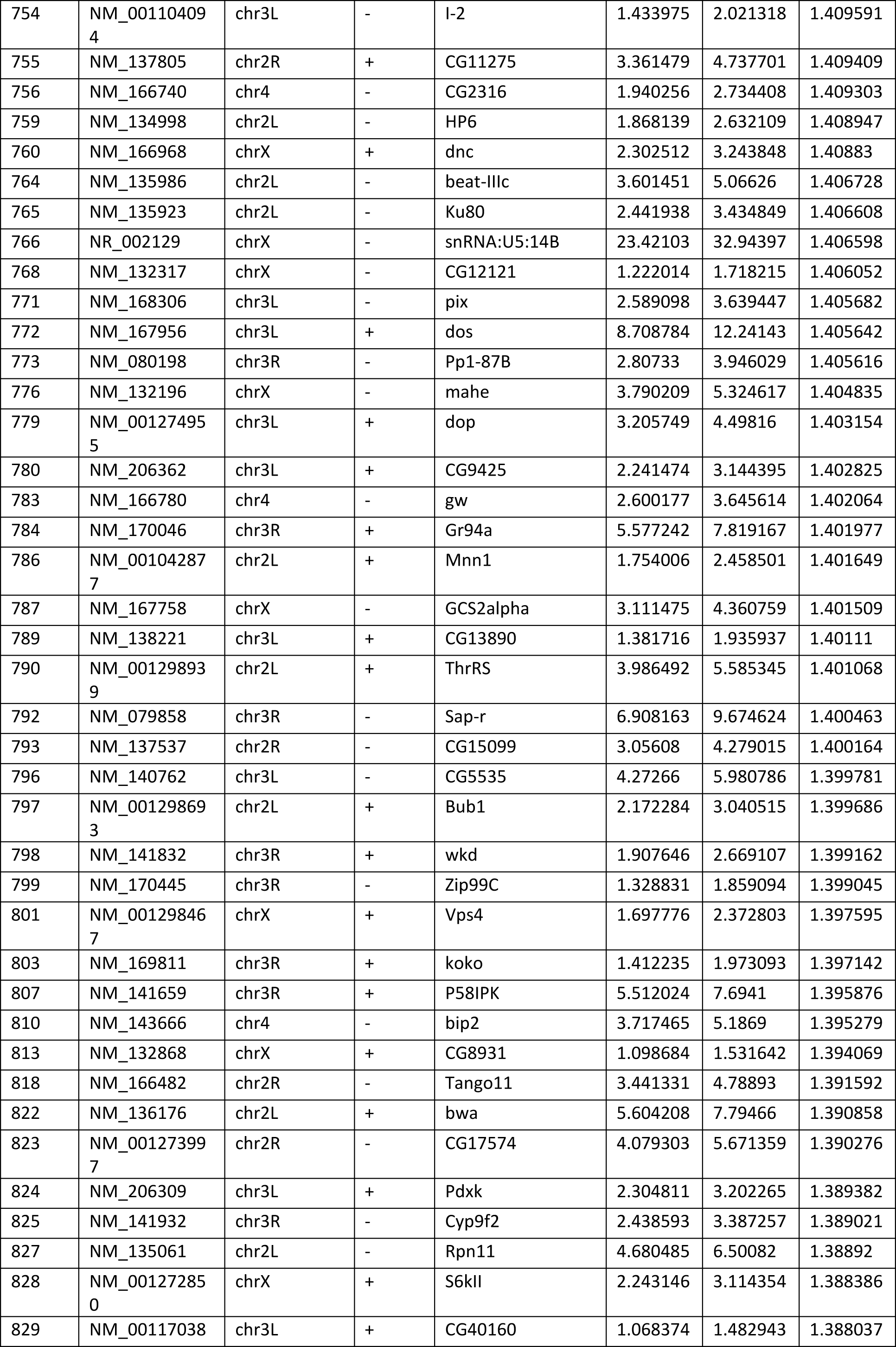
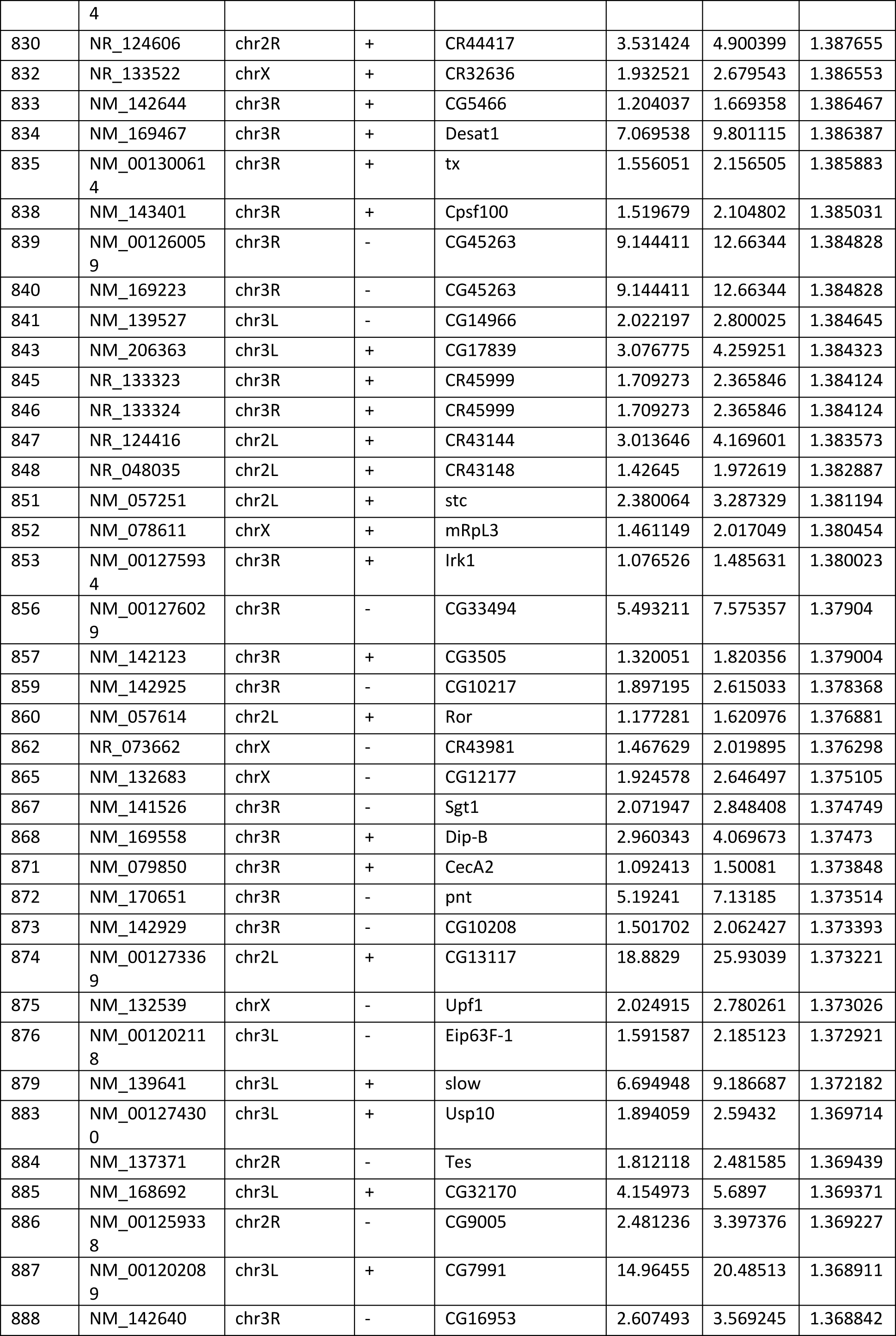
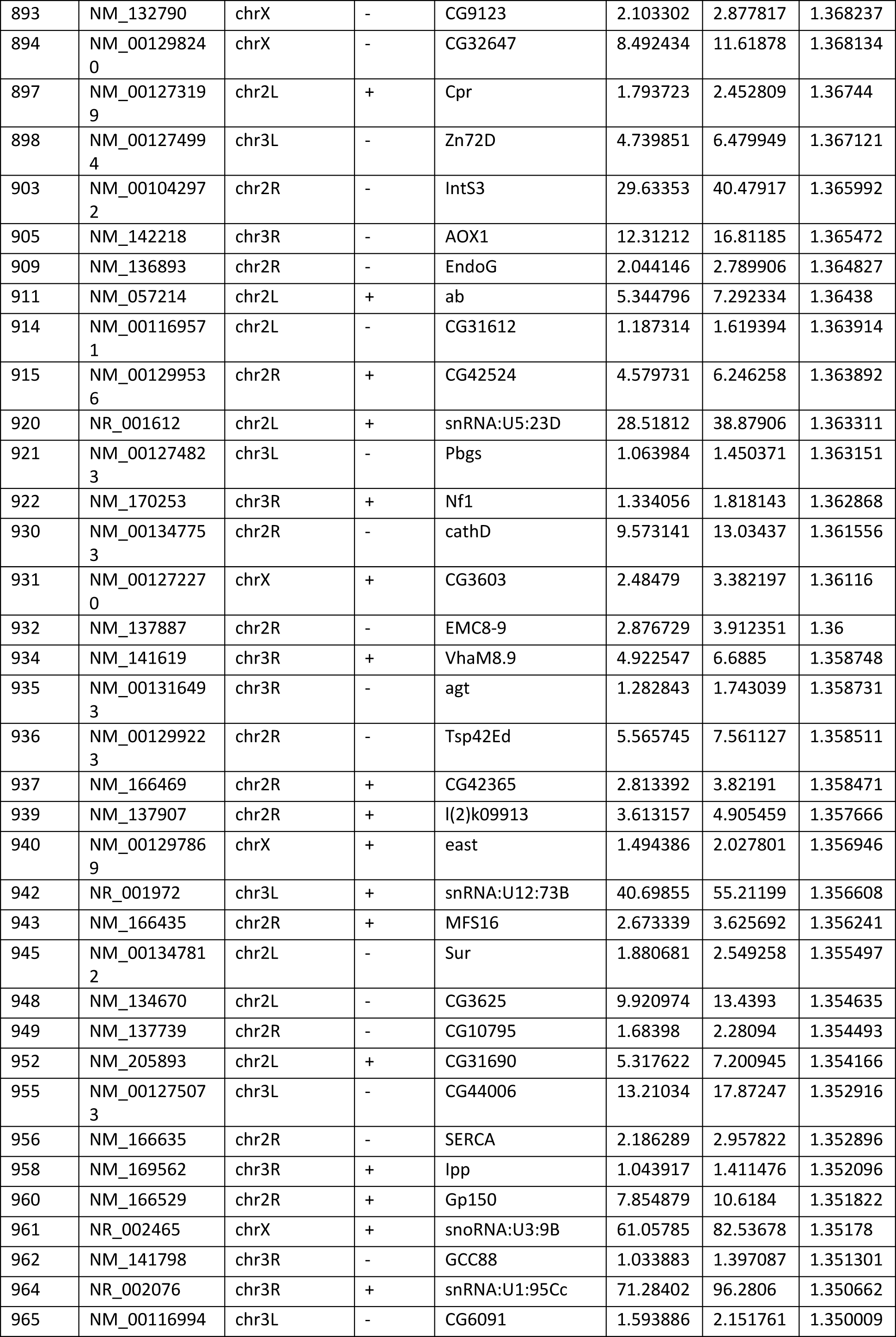
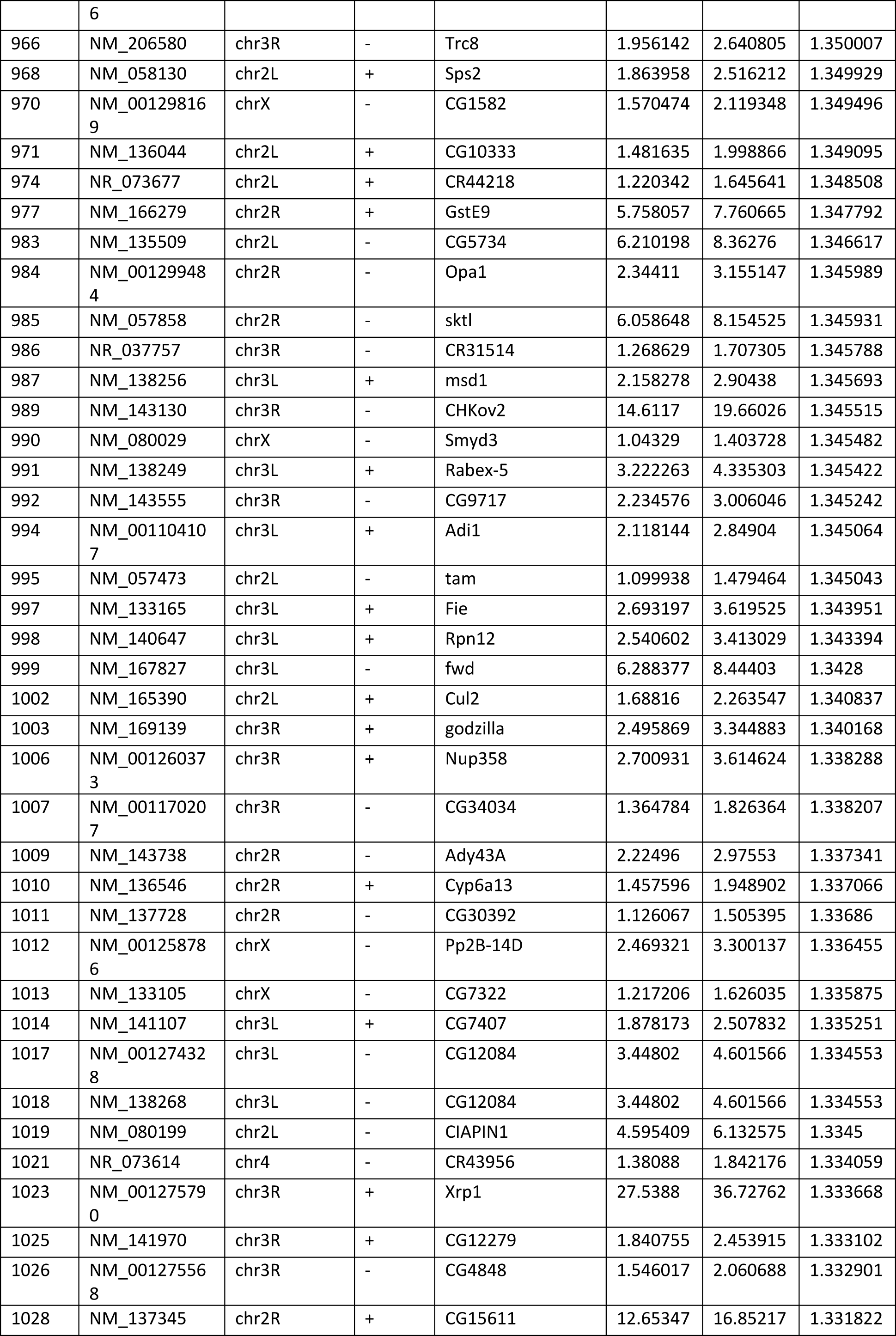
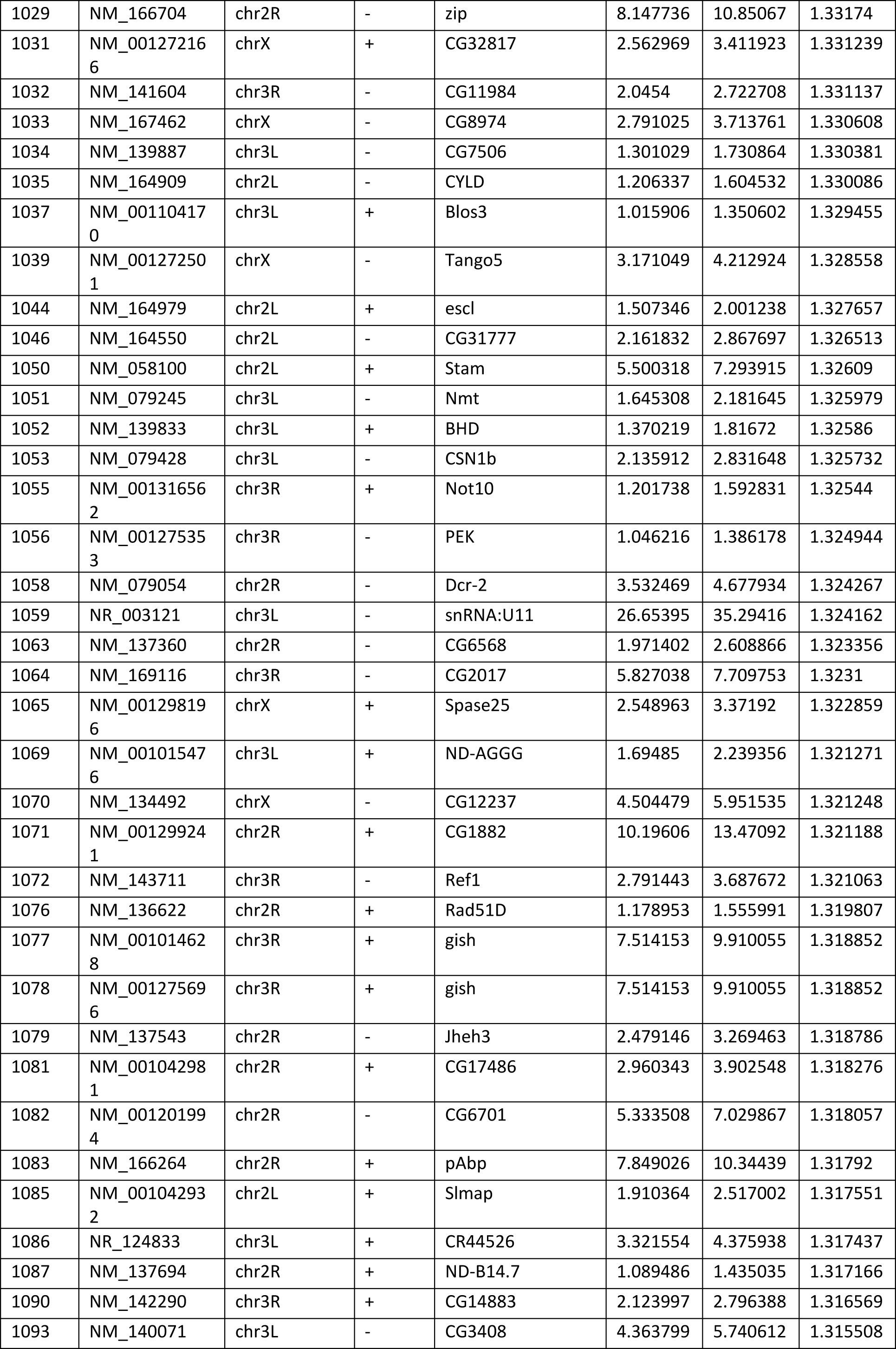
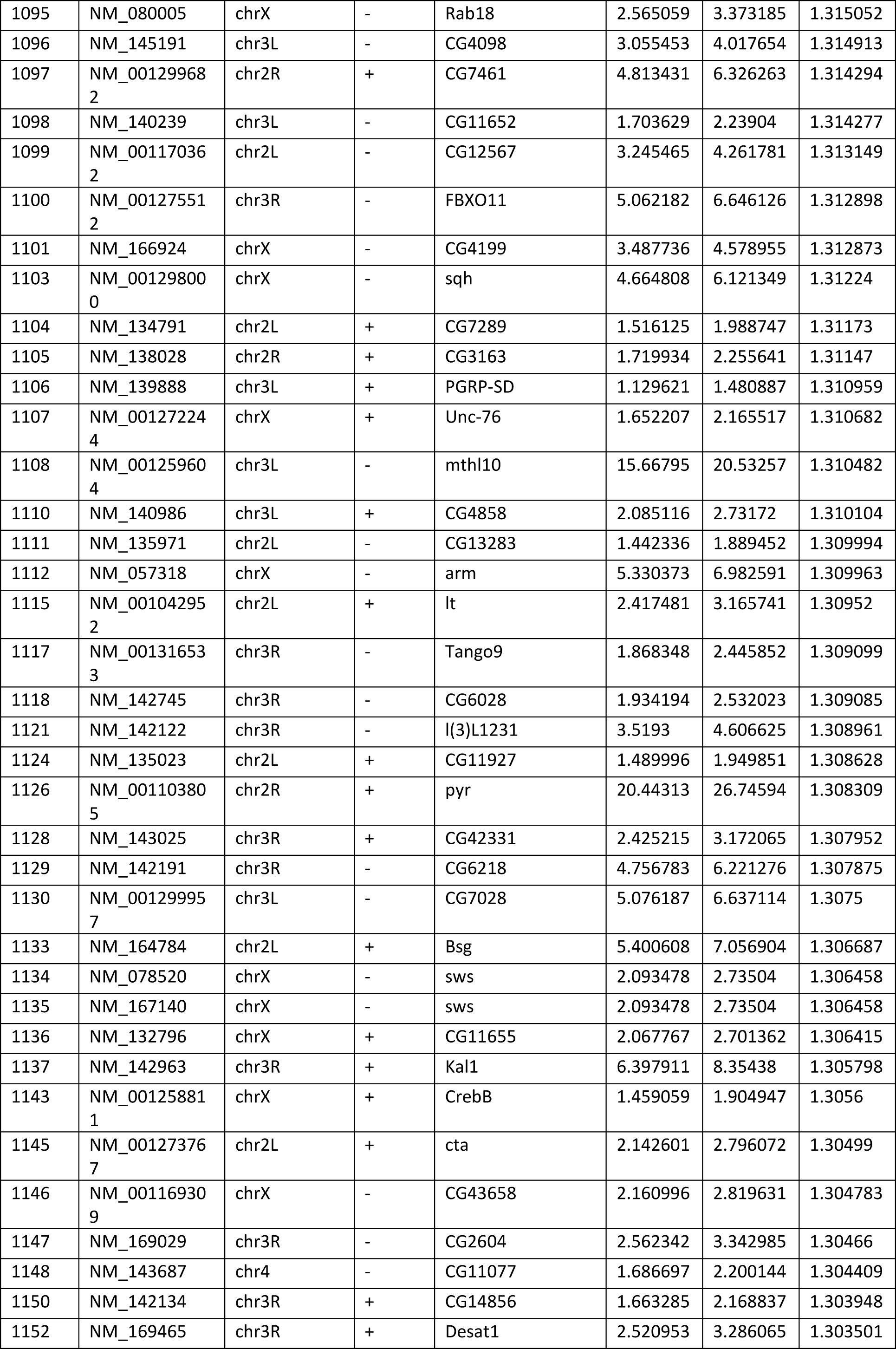
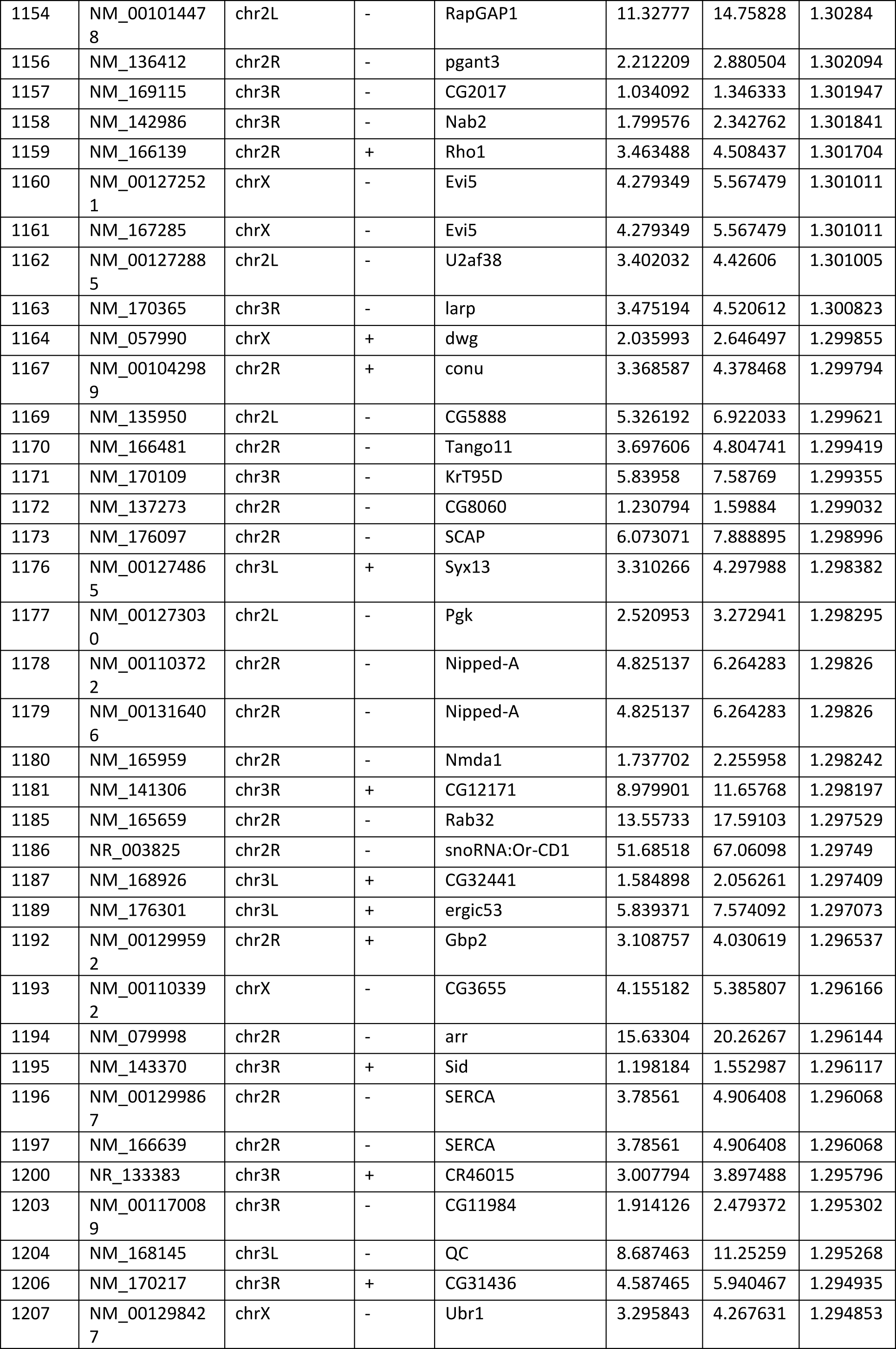
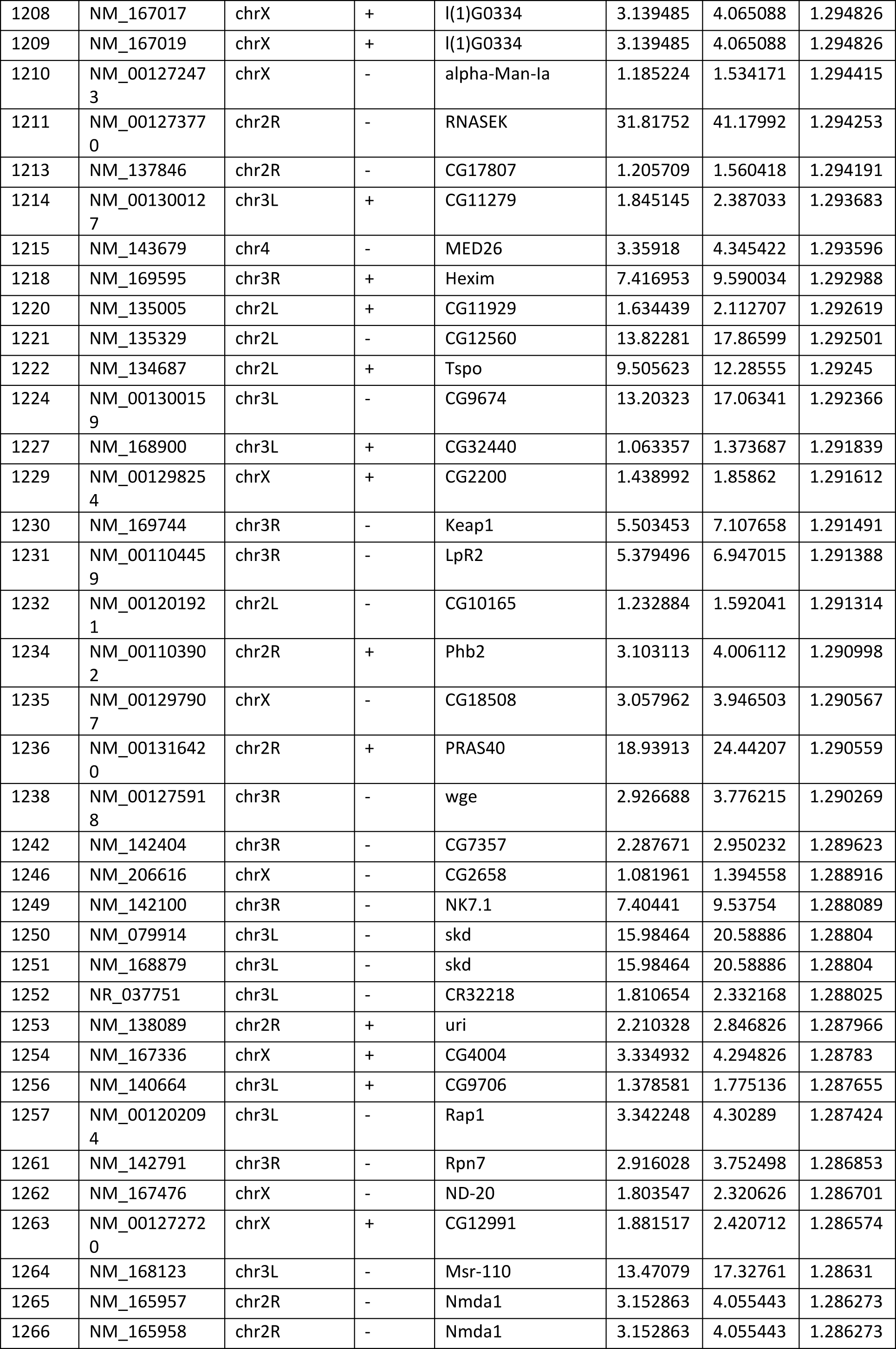
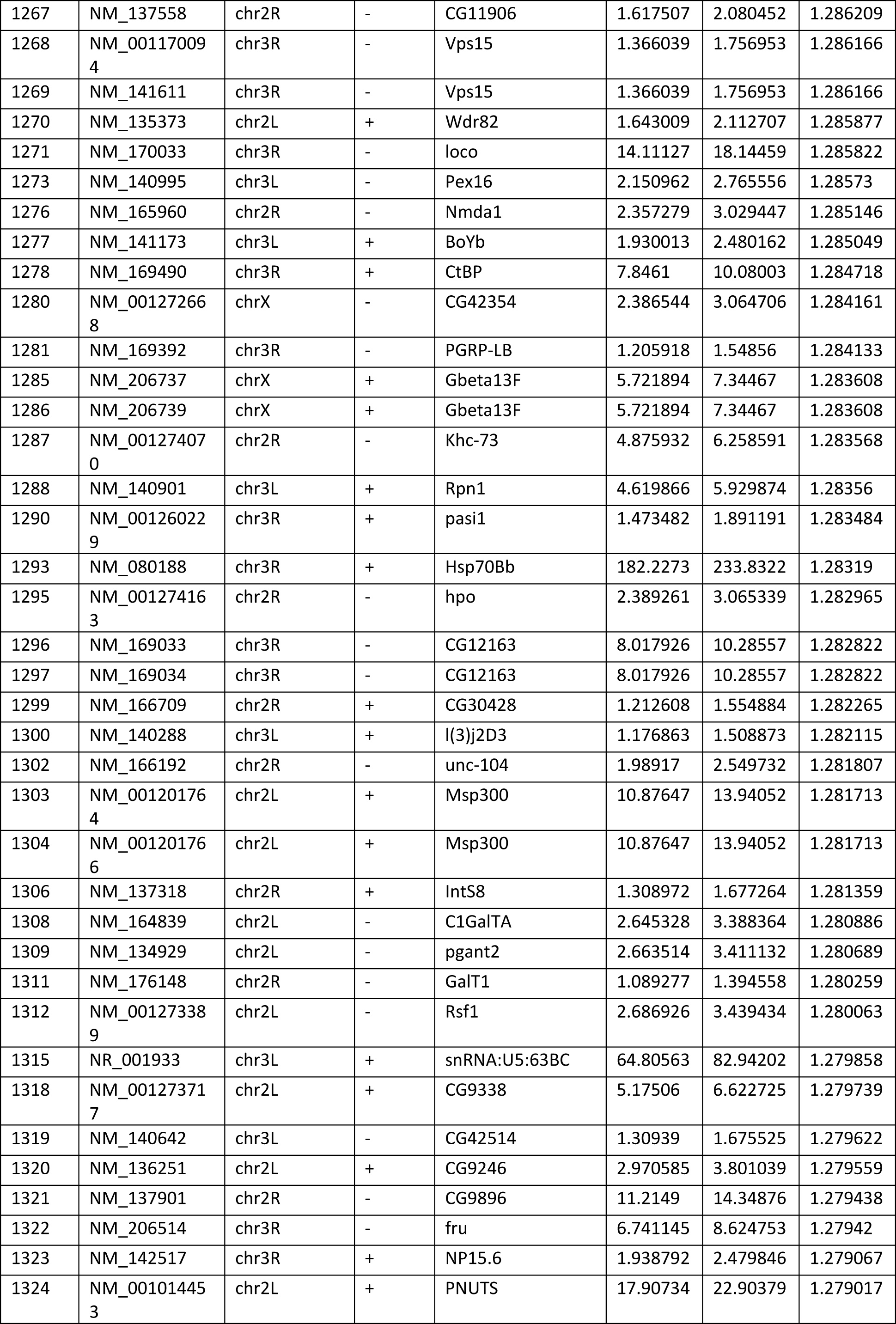
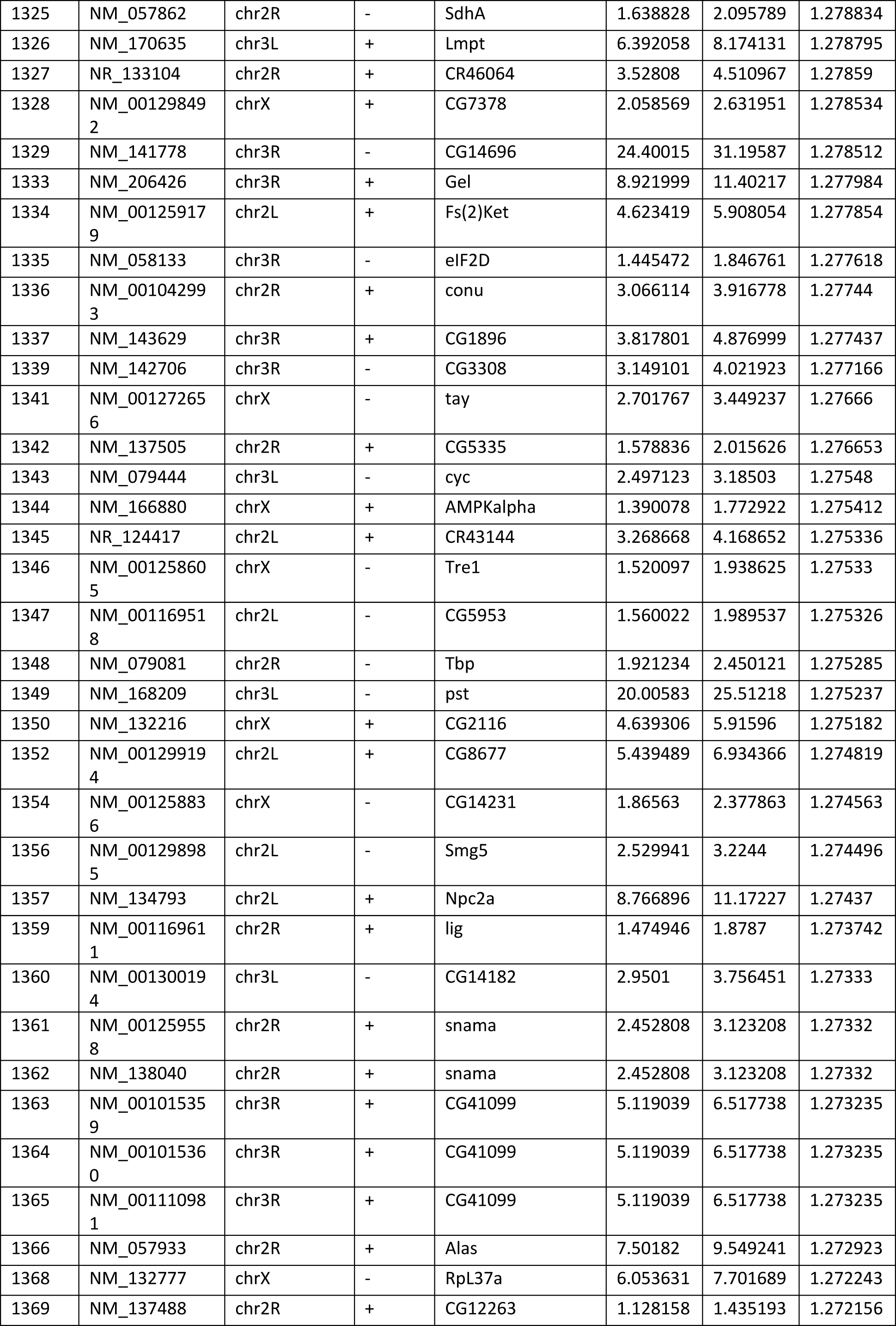
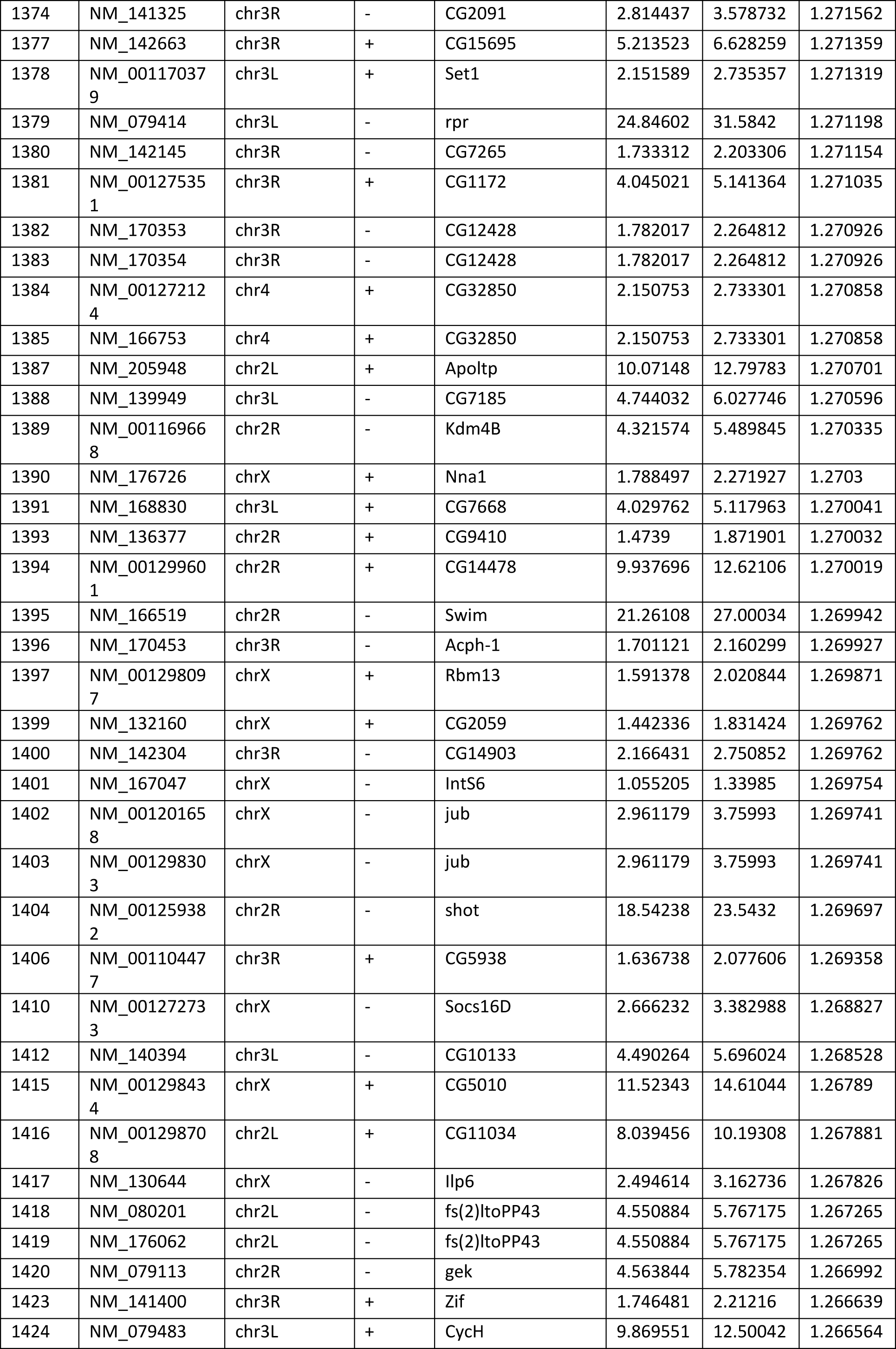
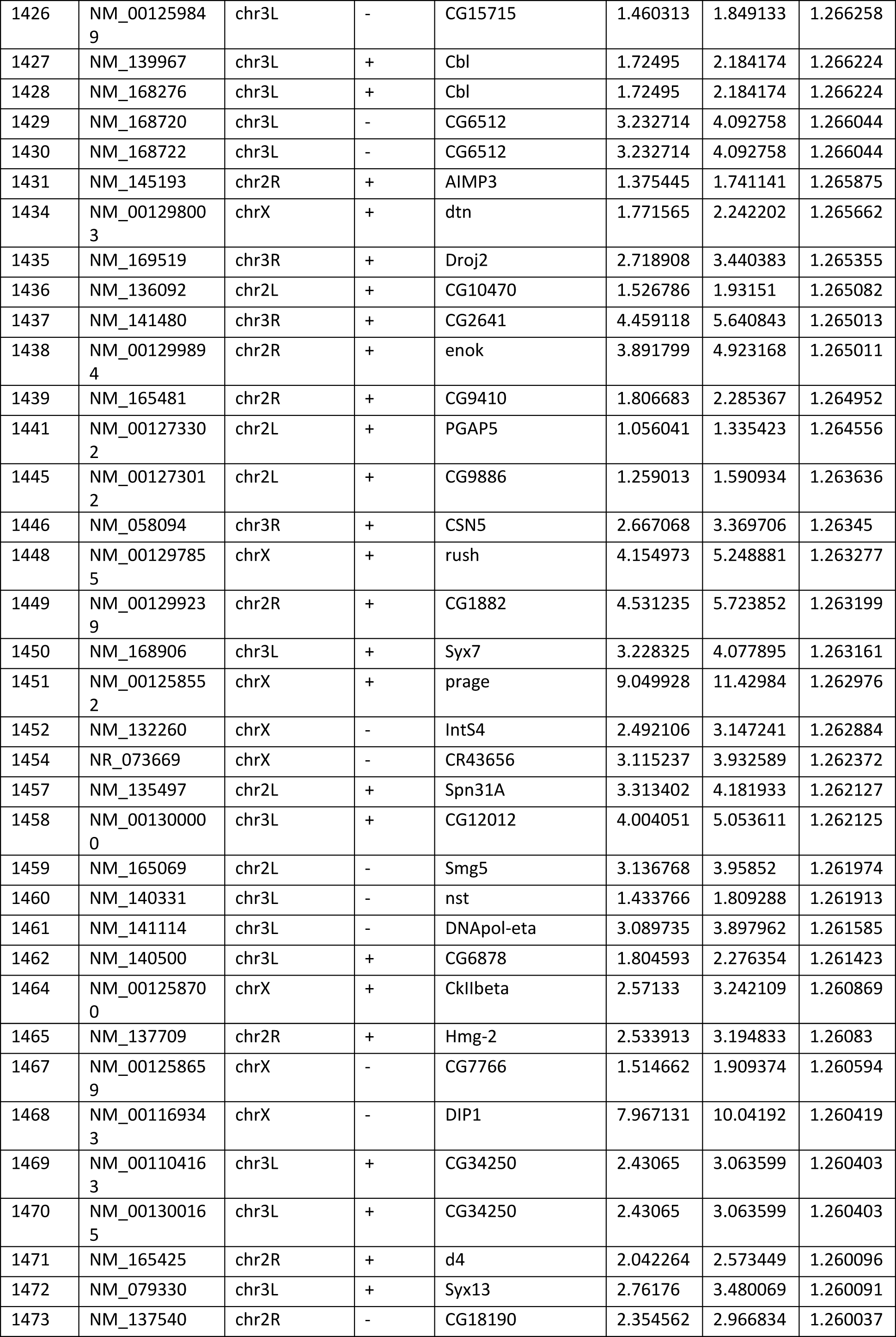
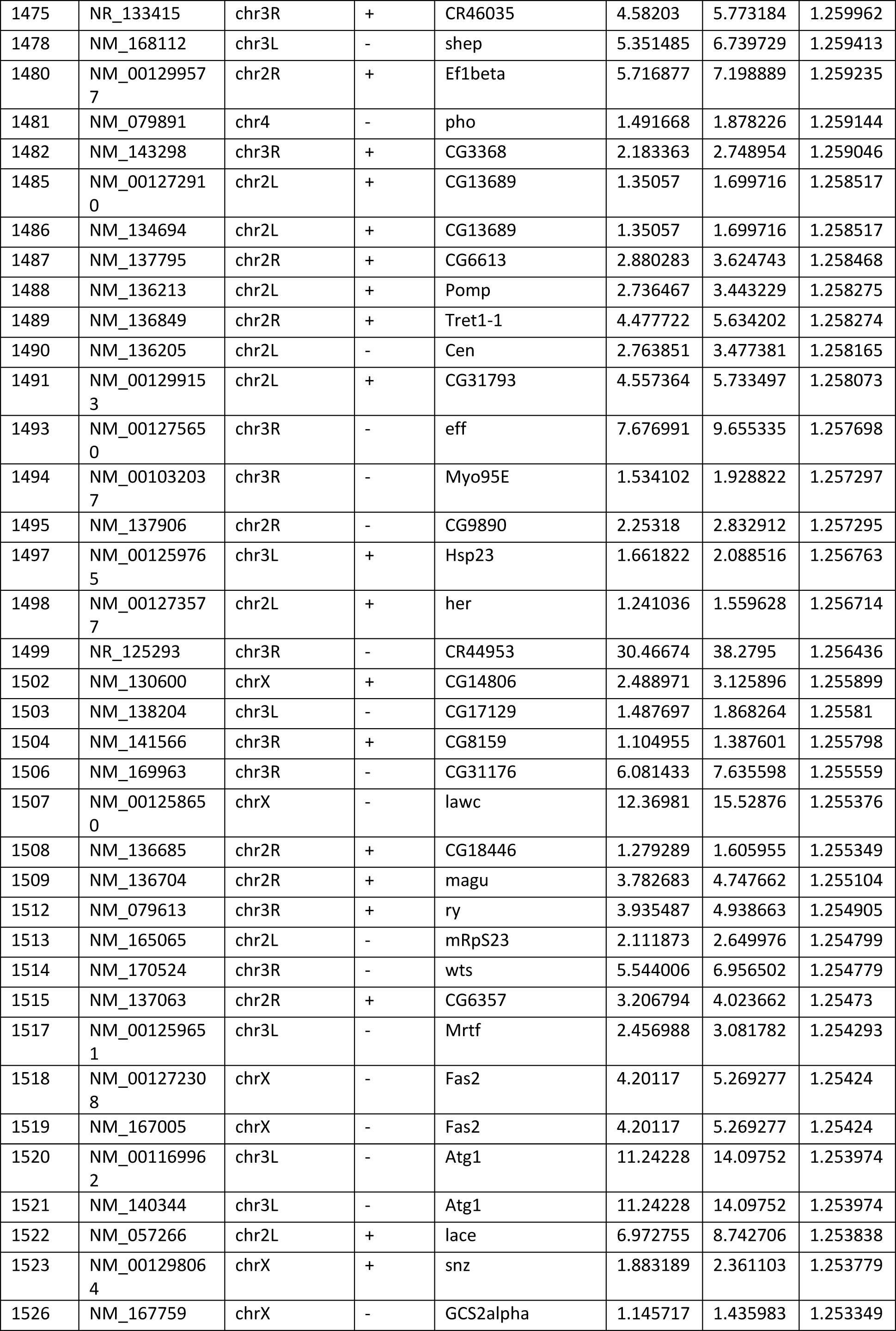
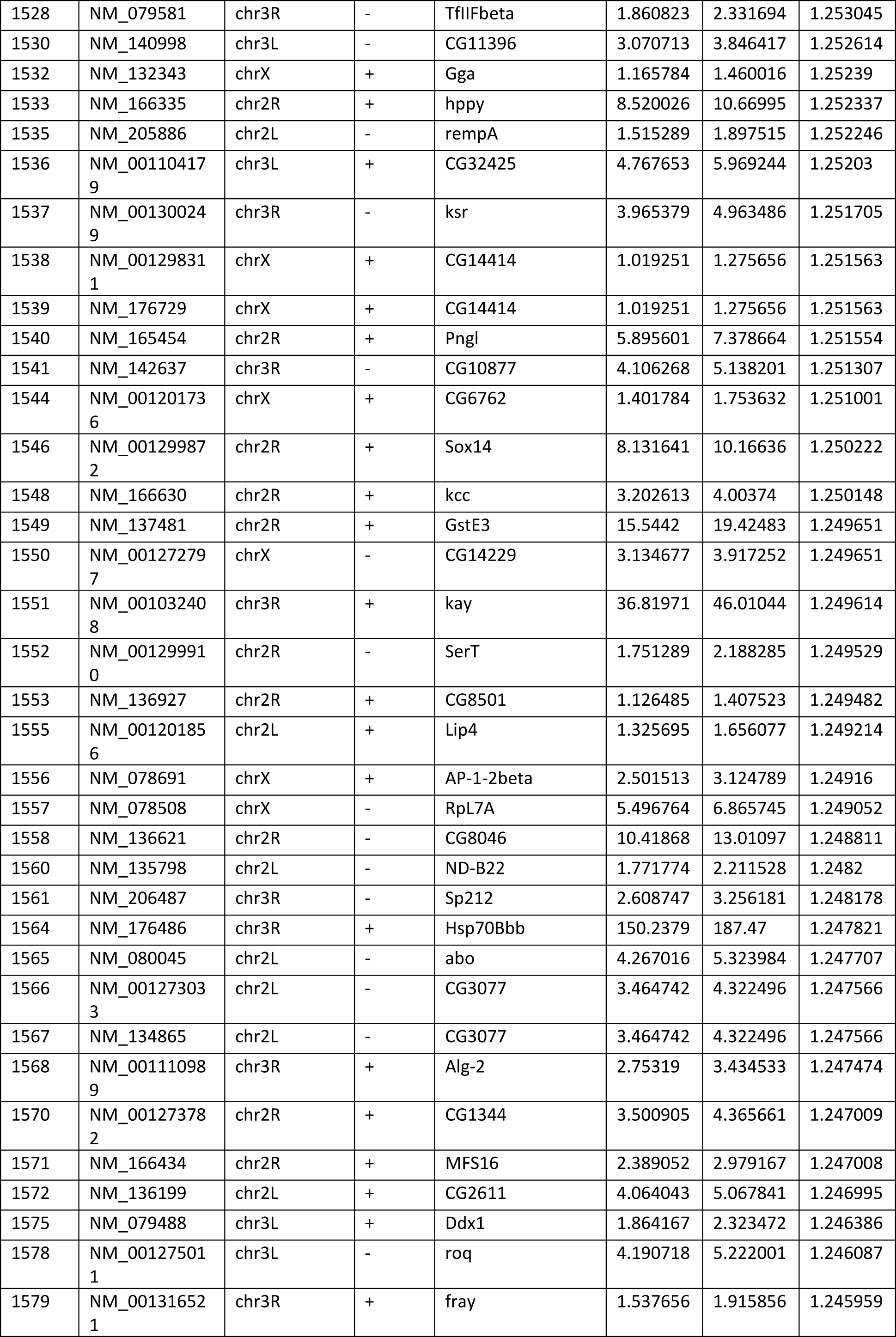
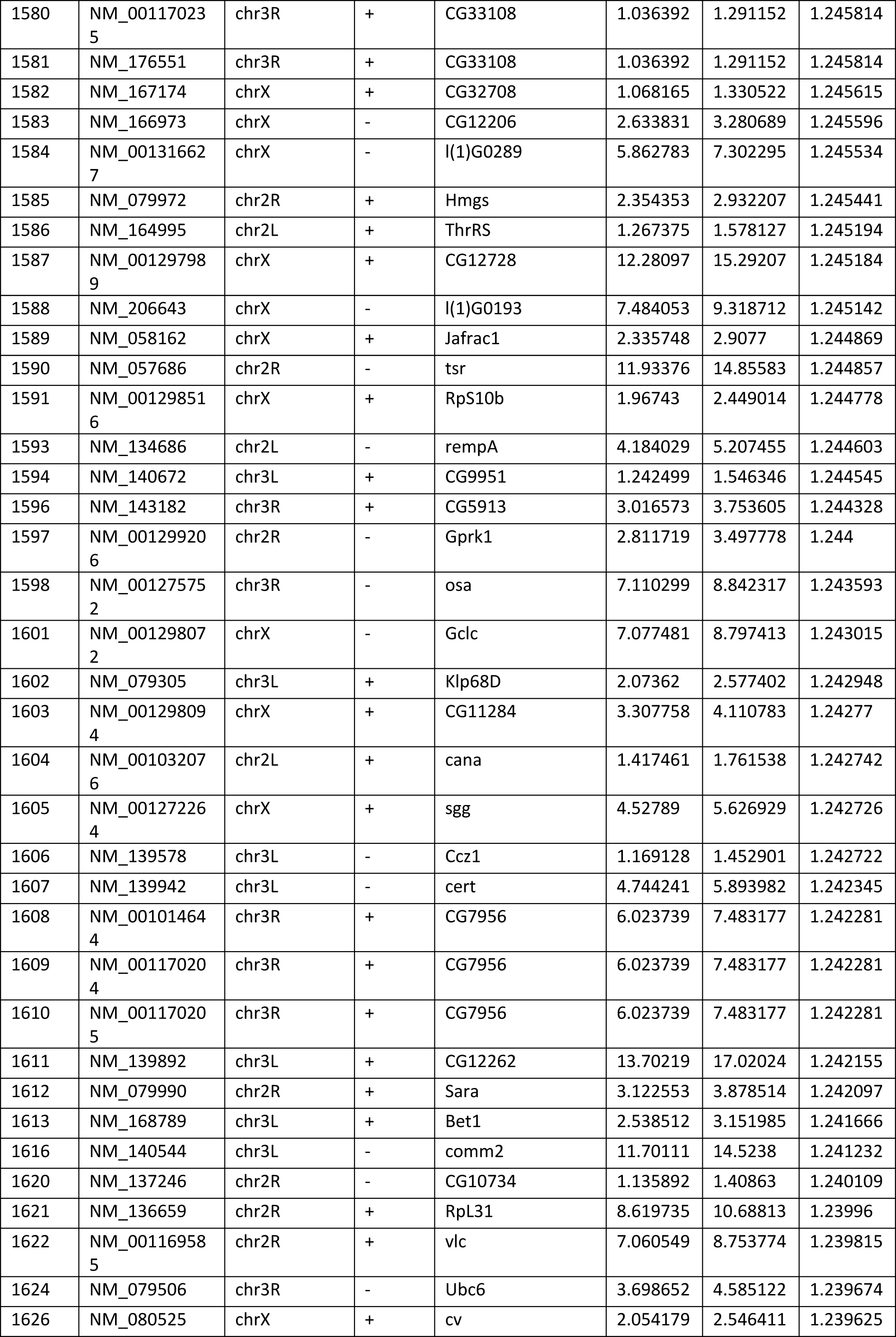
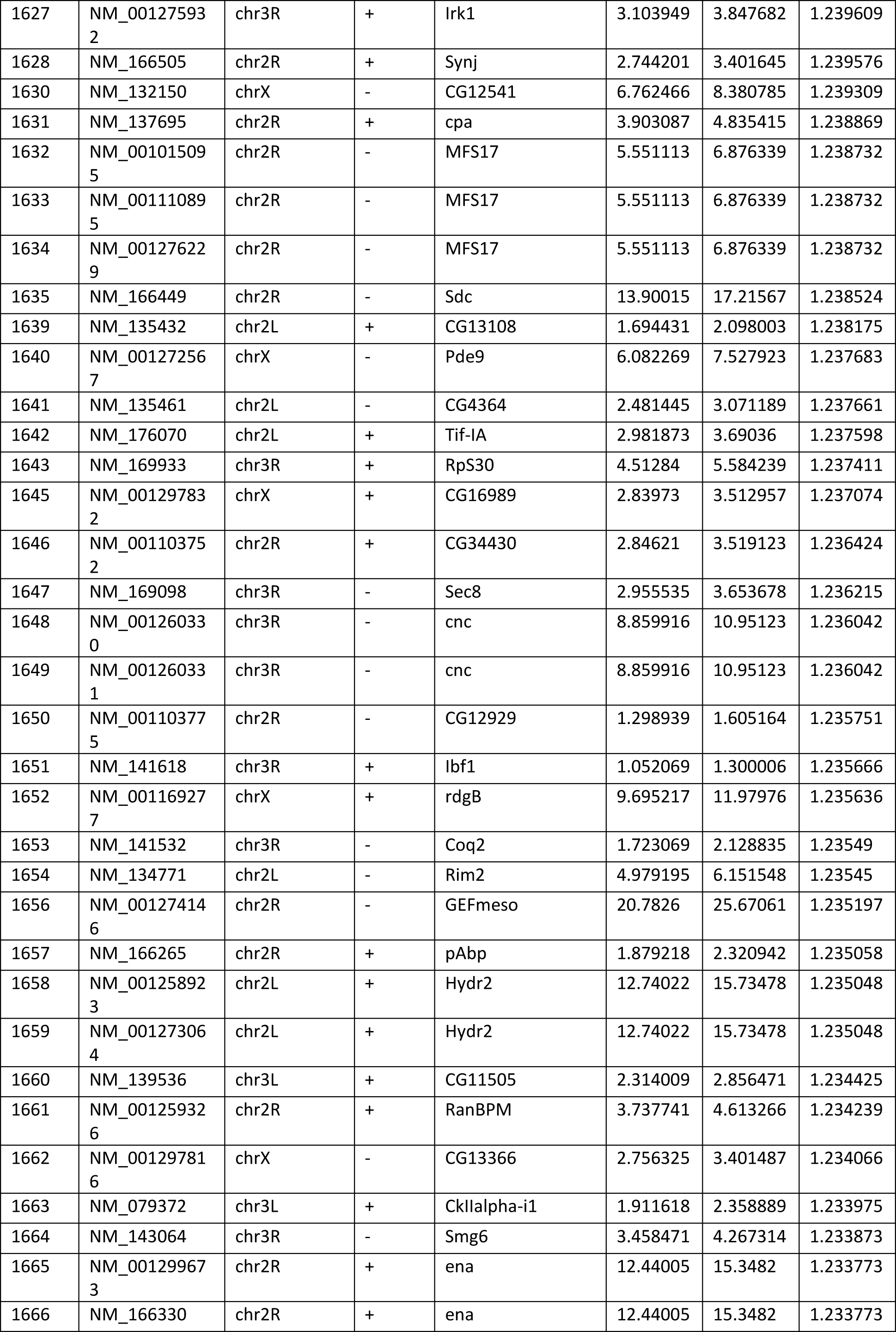
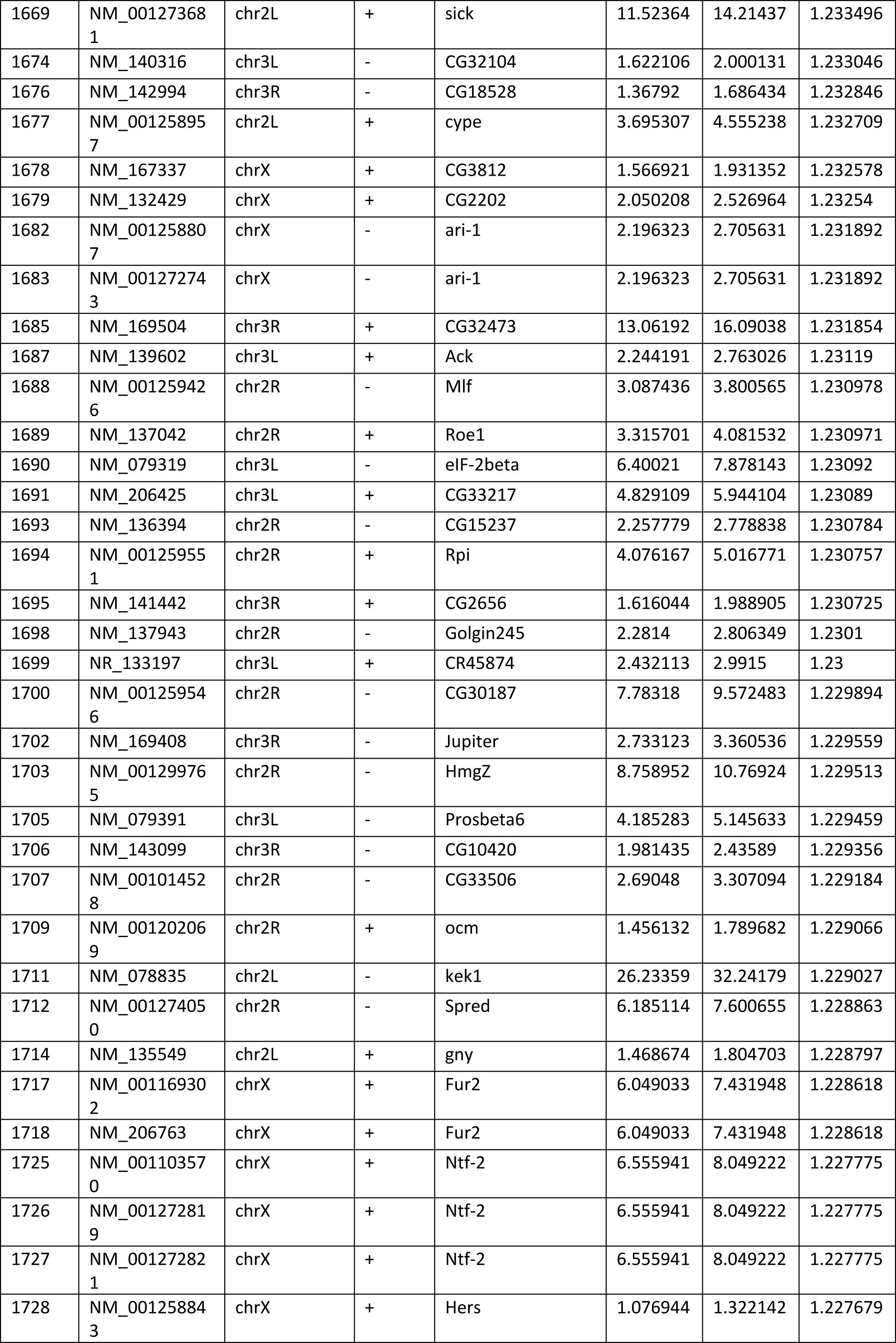
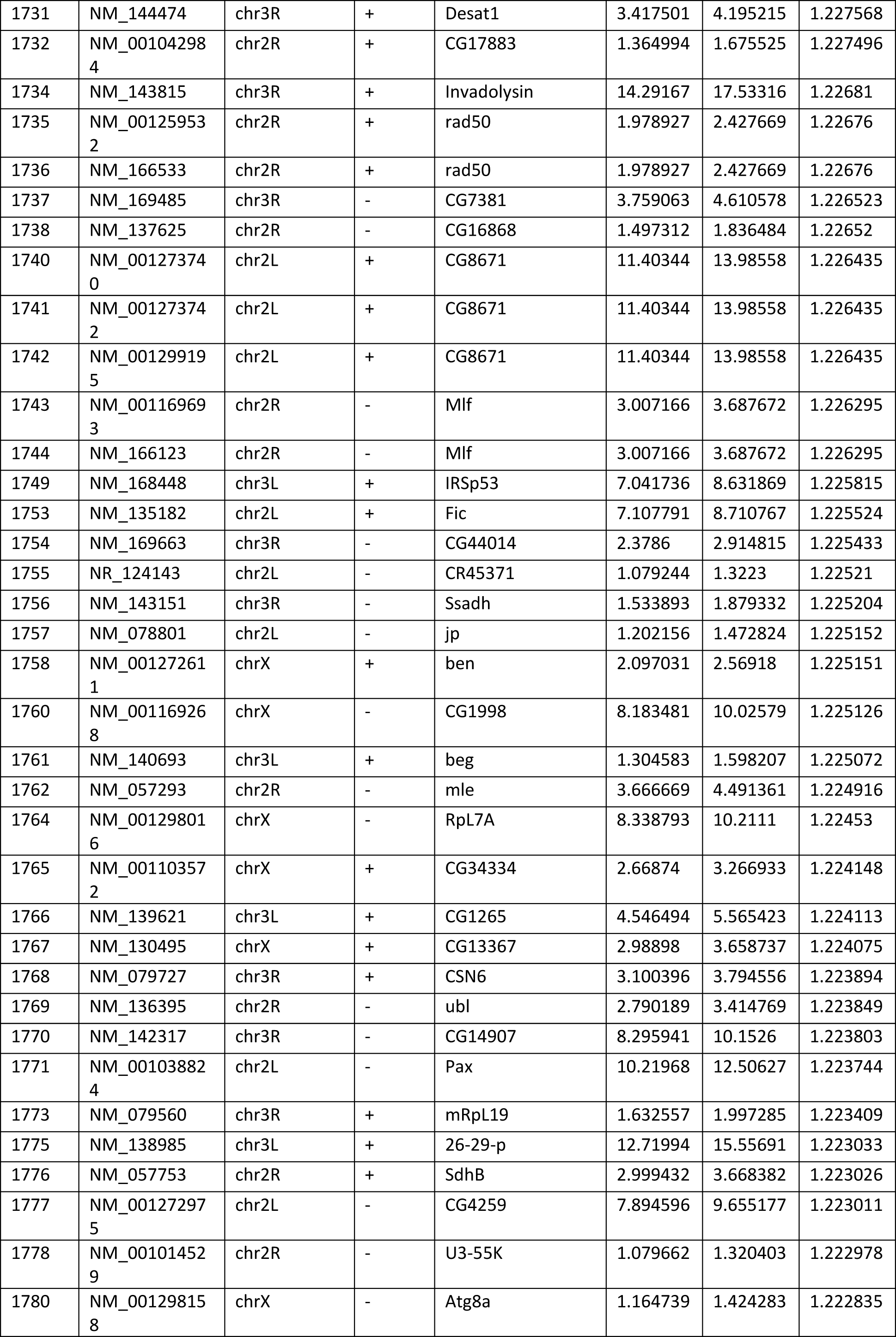
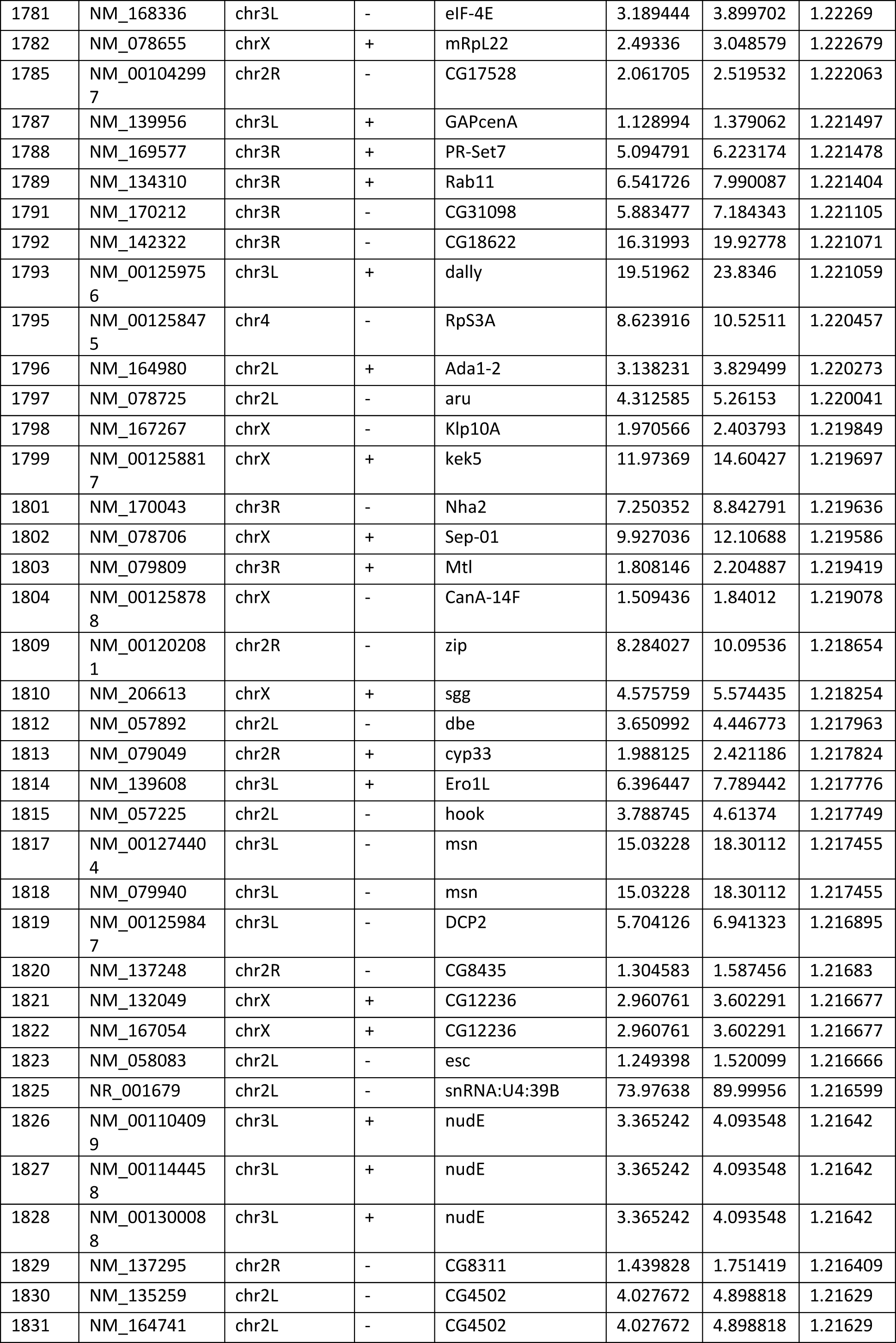
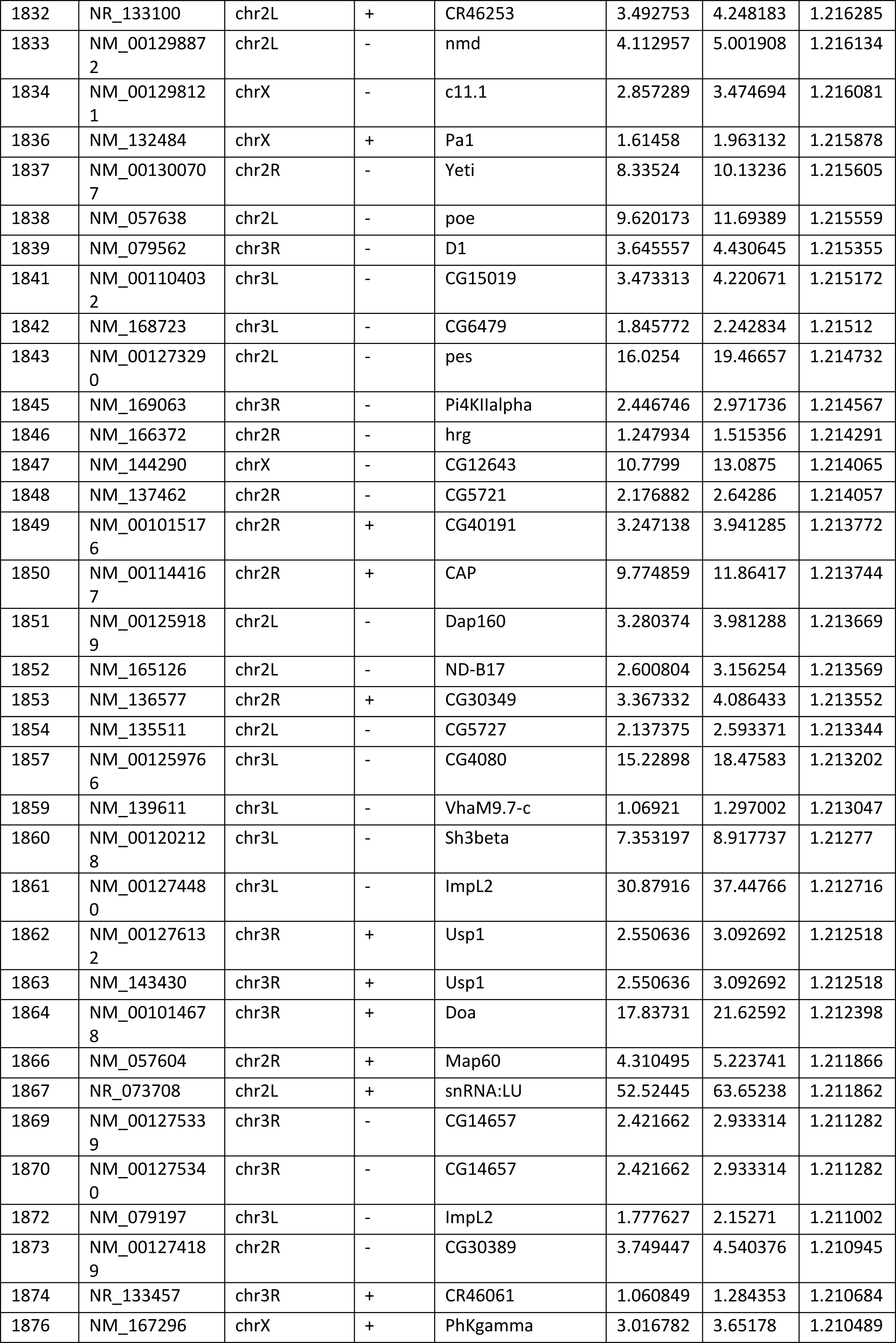
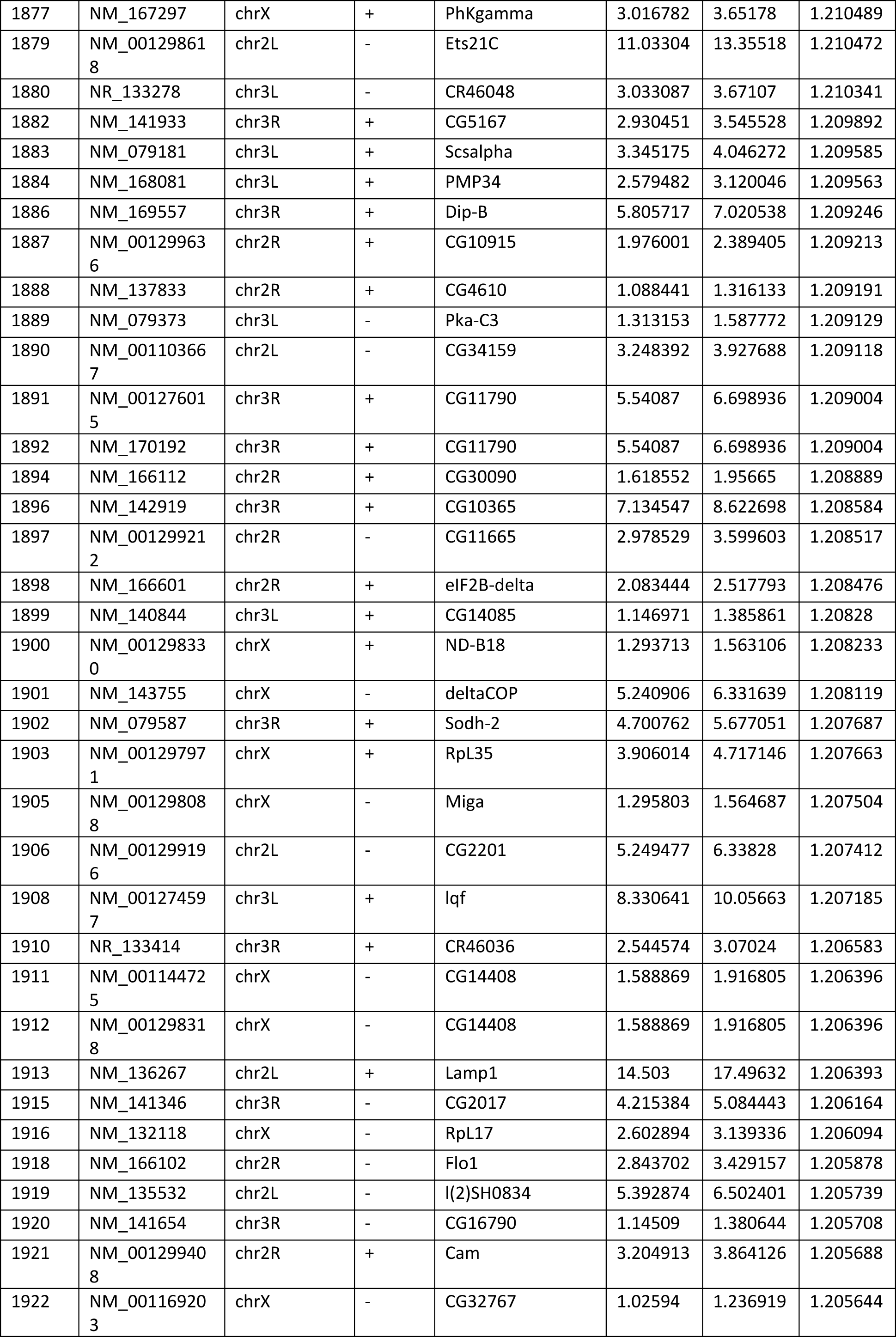
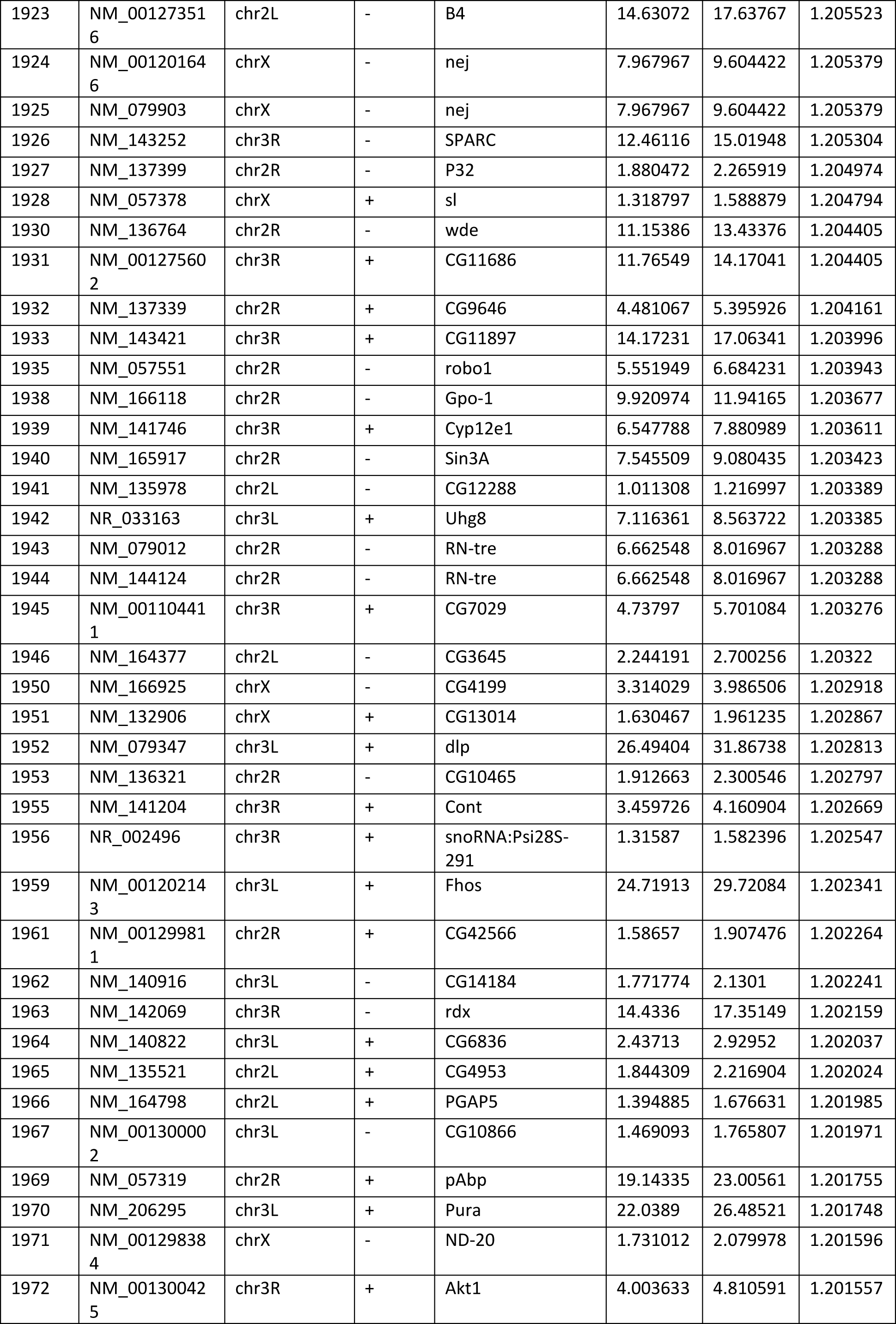
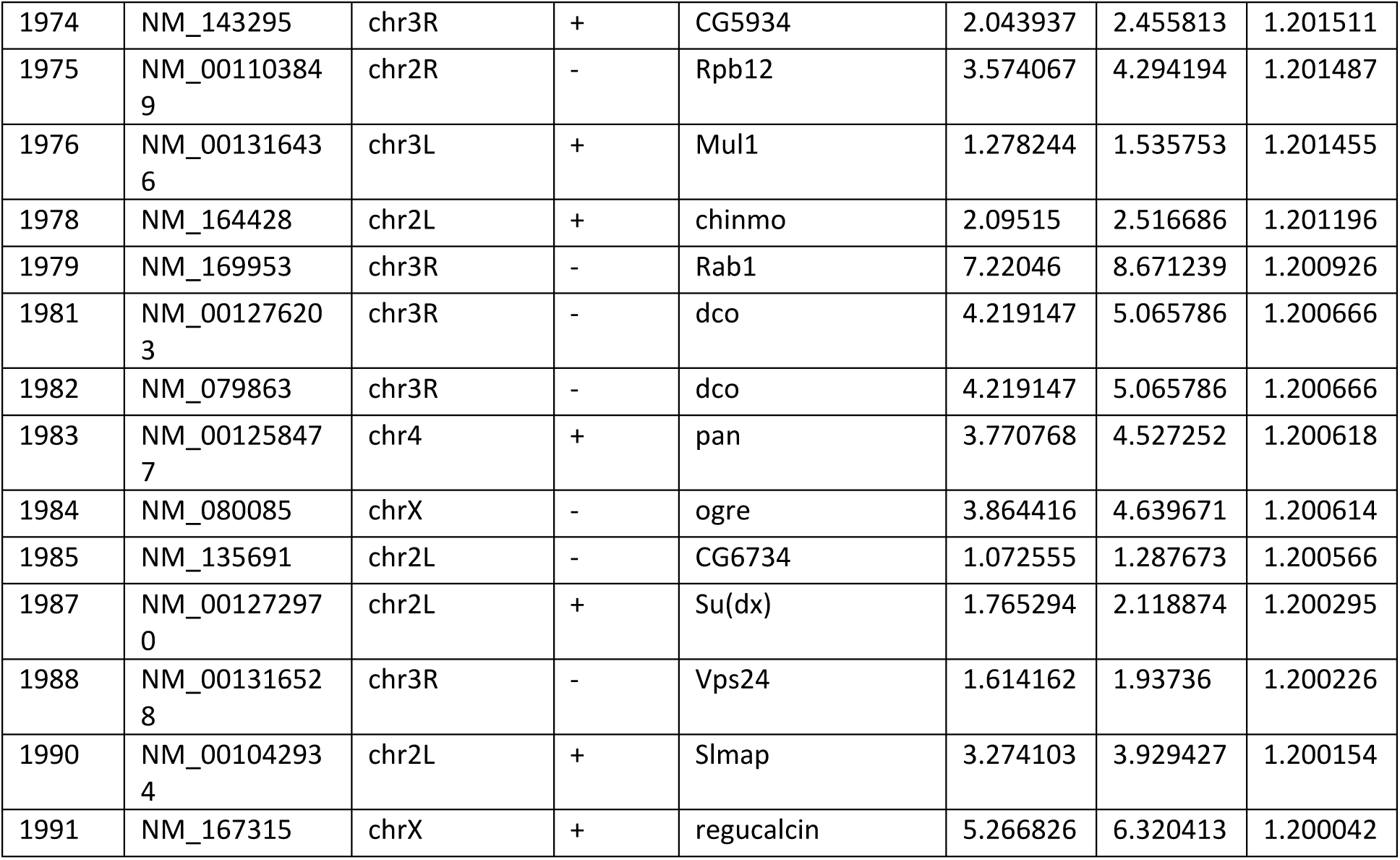
Transcription units with increased unphosphorylated Pol II at TSS in UPF1 depleted S2 cells.

**Table S4.**
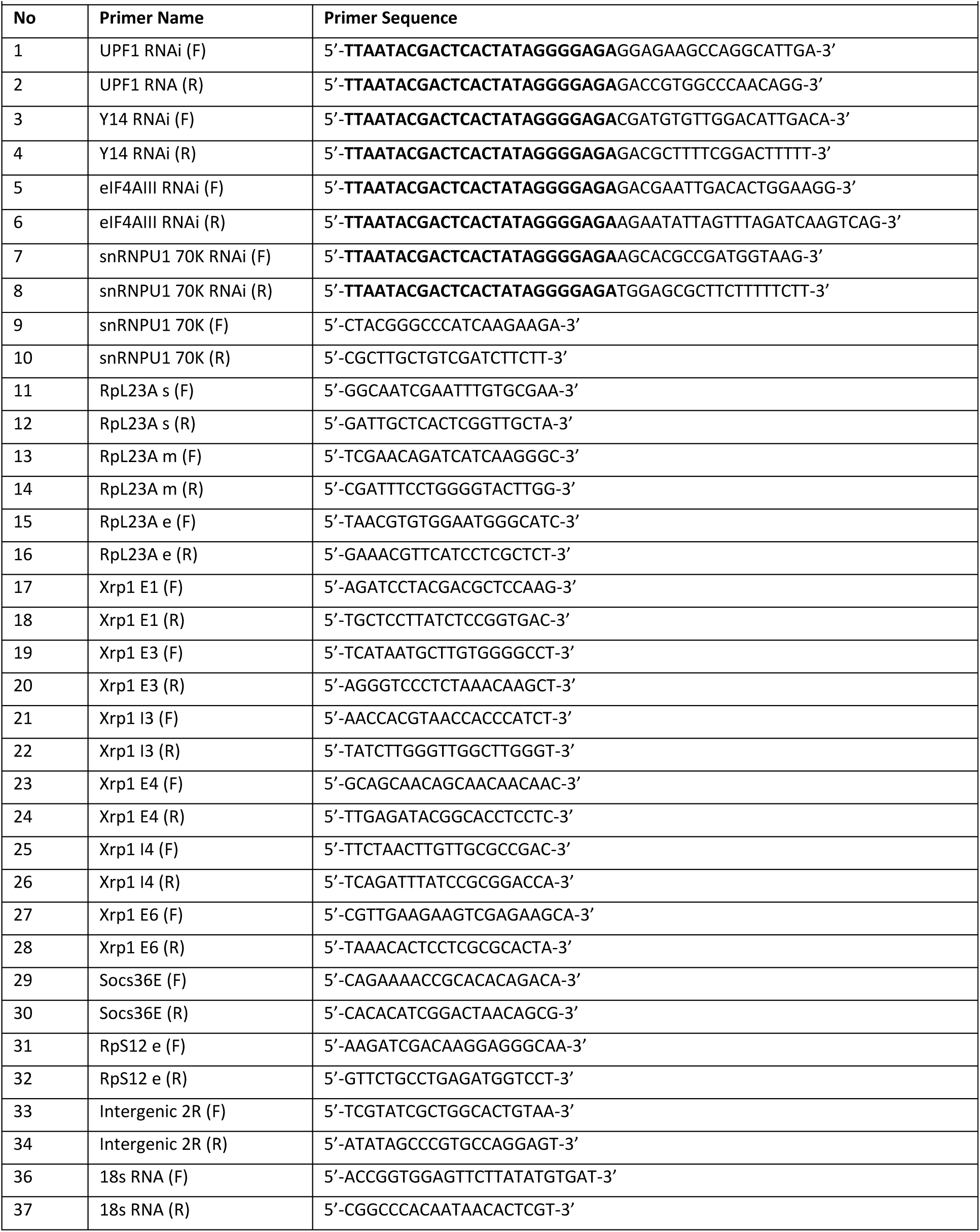
List of primers used in present study (bold=T7 promoter region)

